# Positive Modulators of *N*-Methyl-*D*-Aspartate Receptor: Structure-Activity Relationship Study on Steroidal C-17 and C-20 Oxime Ethers

**DOI:** 10.1101/2025.10.03.680025

**Authors:** Santosh Kumar Adla, Barbora Hrcka Krausová, Bohdan Kysilov, Karel Kudláček, Radko Souček, Miloš Budešínský, Jan Voldřich, Ladislav Vyklický, Eva Kudová

## Abstract

*N*-methyl-D-aspartate receptors (NMDARs) are crucial therapeutic targets, modulated by endogenous neurosteroids like pregnenolone sulfate (**PES**). This study investigates a novel structure-activity relationship approach focusing on the steroidal D-ring, employing the bioisosteric replacement of C-17 or C-20 keto groups with oximes and oxime ethers. We synthesized a series of pregn-5-ene and androst-5-ene derivatives (**11**–**23**) and evaluated their positive allosteric modulator (PAM) activity on recombinant rat GluN1/GluN2B receptors via patch-clamp in HEK293 cells. Our study revealed that pregnenolone-derived C-20 oxime ethers are potent and efficacious PAMs of NMDAR. Several analogues have been demonstrated as more potent than **PES** (E_max_ = 116%; EC_50_ = 21.7 µM). Compound **12** (C-20 ethyl oxime ether, C-3 hemiglutarate) displayed the highest efficacy, potentiating NMDAR currents over 6-fold more than **PES** (E_max_ = 673 ± 121%; EC_50_ = 8.7 ± 1.1 µM). Compound **17** (C-20 methyl oxime ether analogue) exhibited the highest potency, being over 3.5-fold more potent than **PES** (E_max_ = 503 ± 68%; EC_50_ = 6.1 ± 0.4 µM). In contrast, some C-17 analogues and derivatives with bulkier C-20 oxime substituents showed complex modulatory behavior. Promisingly, key compounds demonstrated favorable *in vitro* ADME profiles, including high metabolic stability and, for **12**, excellent thermodynamic solubility. These results validate C-20 oxime ether modification of the pregnenolone scaffold as an effective strategy for generating potent NMDAR PAMs with potentially superior efficacy and drug-like properties compared to endogenous modulators.

## Introduction

Endogenous neurosteroids such as 20-oxo-5β-pregnan-3α-yl sulfate (pregnanolone sulfate, **PAS**, Fig. 1A) and 20-oxo-pregn-5-en-3β-yl sulfate (pregnenolone sulfate, **PES**, Fig. 1A) are well-established modulators of *N*-methyl-D-aspartate receptors (NMDARs) (Korinek, Kapras et al. 2011, Kudova 2021). PES acts as a positive allosteric modulator (PAM) of NMDARs, whereas **PAS** functions as a negative allosteric modulator (NAM). NMDARs are heteromeric, Ca^2+^-permeable ion channels that play a critical role in synaptic plasticity and neurotransmission (Hansen, Wollmuth et al. 2021). Consequently, their activity has been identified as a promising pharmacological target for a range of neurological and psychiatric disorders, including epilepsy and autism (Gataullina, Bienvenu et al. 2019, Celli and Fornai 2021, Mangano, Riva et al. 2022), post-traumatic stress disorder (Jumaili, Trivedi et al. 2022), depression (Henter, Park et al. 2021), pain (Obeng, Hiranita et al. 2021, Meng and Shen 2022), stroke (Shen, Xiang et al. 2022) and various neurodegenerative diseases (Hardingham and Bading 2010).

**Fig. 1.**
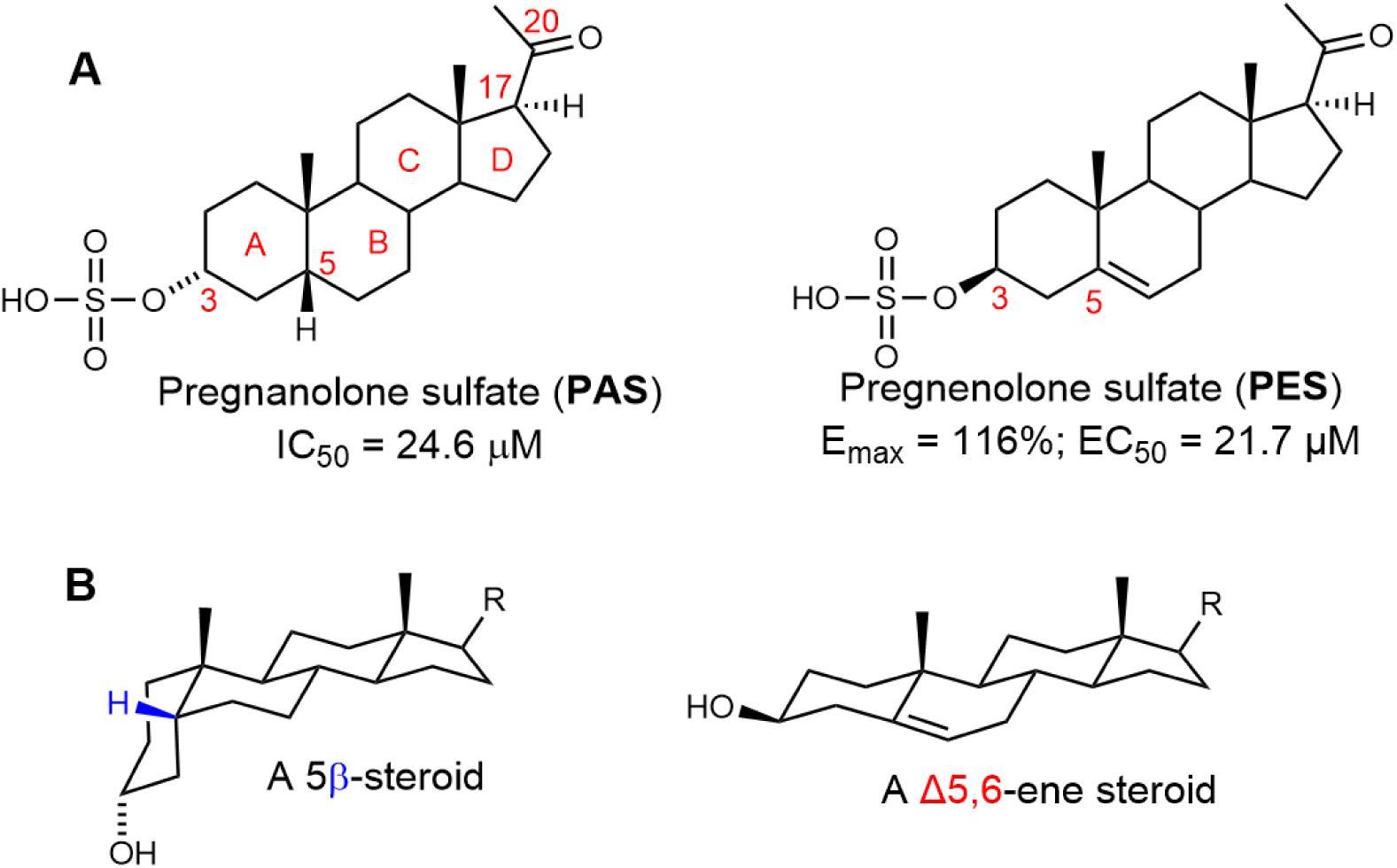
Steroidal positive and negative allosteric modulators of NMDA receptors. *(A)* Structures of endogenous neurosteroids pregnanolone sulfate (**PAS**) and pregnenolone sulfate (**PES**) with ring letters and skeleton numbering relevant for this publication. *(B)* Simplified visualization of a bent molecule of 5β- and planar Δ5,6-ene steroids.

Mutations in NMDA receptor subunit genes are well-documented neurodevelopmental and neuropsychiatric disorders (Hamdan, Gauthier et al. 2011, Tarabeux, Kebir et al. 2011, Hu, Chen et al. 2016, Fedele, Newcombe et al. 2018, Tang, Liu et al. 2020, Elmasri, Lotti et al. 2022). Many disease-associated de novo variants result in loss-of-function phenotypes with reduced receptor activity and impaired glutamatergic signalling. Consequently, pharmacological strategies to enhance NMDAR function are a promising therapeutic direction.

Beyond these endogenous neurosteroids, numerous synthetic neuroactive steroid analogues have been shown to allosterically modulate NMDARs (Kudova, Chodounska et al. 2015, Slavikova, Chodounska et al. 2016, Adla, Slavikova et al. 2017, Adla, Slavikova et al. 2018, Krausova, Slavikova et al. 2018, Smidkova, Hajek et al. 2019). The mode of action of these compounds is primarily dictated by the structural features of the steroidal skeleton. Specifically, NMDAR inhibition is associated with a bent steroid ring system characterized by 3α-hydroxy-5β-stereochemistry (Fig. 1B). In contrast, potentiation of NMDARs is predominantly linked to a planar skeleton with a 3β-hydroxy-Δ5,6-ene configuration (featuring a double bond in the B-ring, Fig. 1B), a structural motif that has been extensively studied (Weaver, Land et al. 2000, Krausova, Slavikova et al. 2018).

As part of our ongoing research on neurosteroid modulators of NMDARs, we have conducted several structure–activity relationship (SAR) studies to elucidate the pharmacophore requirements for steroidal PAMs and NAMs. Our findings indicate that NMDAR modulatory activity is highly dependent on the presence of a charged substituent, such as a sulfate moiety (Borovska, Vyklicky et al. 2012). Specifically, replacing the sulfate group with uncharged substituents abolishes NAM activity, whereas substitution with dicarboxylic acid esters generally preserves activity (Borovska, Vyklicky et al. 2012, Krausova, Slavikova et al. 2018).

Further, we have explored the impact of non-lipophilic substituents at the C-17 position of the steroidal D-ring on *in vitro* activity (Kudova, Chodounska et al. 2015, Krausova, Slavikova et al. 2018). Our studies demonstrated that the allosteric modulatory effect - either positive or negative - correlates with the lipophilicity of the compound. For example, compound **1** (Fig. 2) featuring an isobutyl chain at C-17, inhibited currents of GluN1/GluN2B receptors with an IC_50_ of 90 nM (Kudova, Chodounska et al. 2015). Similarly, removing the C-17 substituent in compound **2** resulted in potentiation of GluN1/GluN2B receptor responses, with an E_max_ of 452% and an EC_50_ of 7.4 µM (Krausova, Slavikova et al. 2018).

Interestingly, neither our SAR studies nor those by other researchers have extensively investigated polar modifications of the steroidal D-ring. Based on literature data, substitution of C-17 with a ketone is expected to yield inactive compounds, as observed for dehydroepiandrosterone and dehydroepiandrosterone sulfate (Yaghoubi, Malayev et al. 1998). Contrary to this expectation, our 2018 study (Krausova, Slavikova et al. 2018) revealed that compounds **5**–**10** function as PAMs with moderate efficacy. While compounds within this structural series exhibited similar maximal efficacy (E_max_ ranging from 154% to 226%), their potency varied significantly, with EC_50_ values between 16 and 151 µM.

**Fig. 2.**
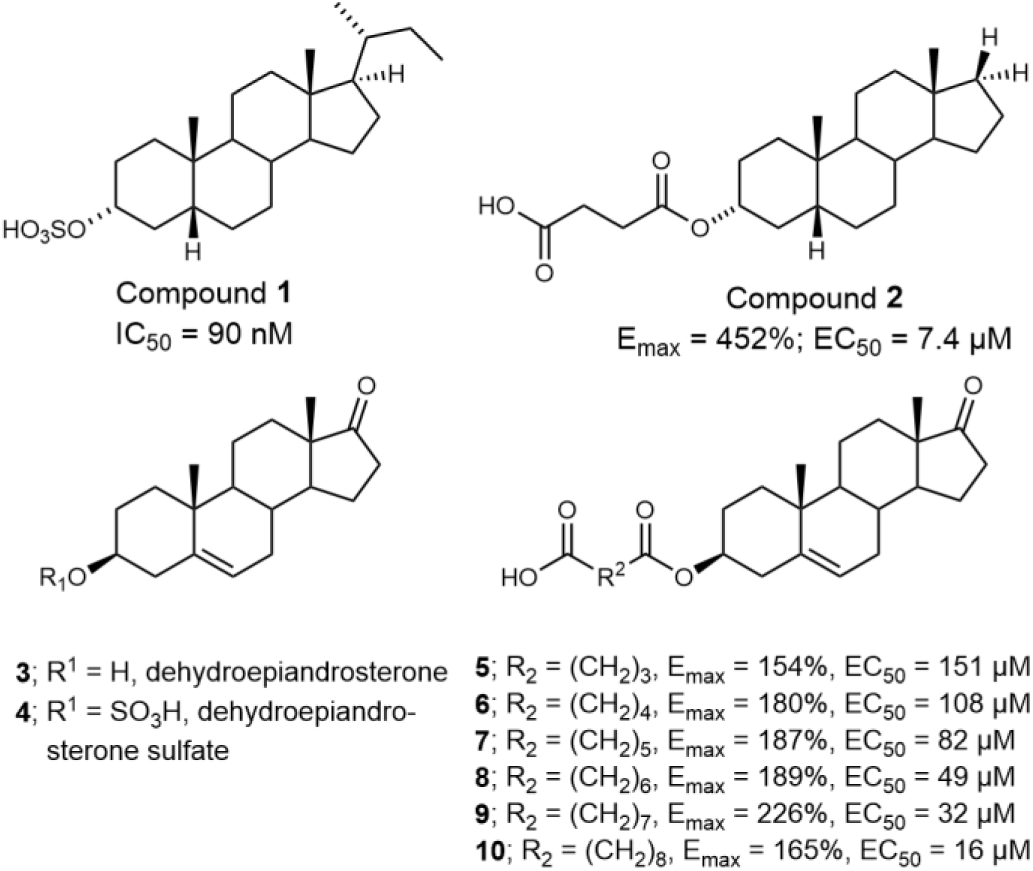
Structures of PAMs and NAMs of NMDARs.

The replacement of the keto group with a suitable surrogate, or (bio)isostere, can represent an effective strategy for SAR studies targeting novel analogues as such modification should preserve at least similar biological activities of the parent compound (Patani and LaVoie 1996, Meanwell 2011). For our study, the replacement of C-17 and C-20-ketone groups with a lipophilic-like surrogate was proposed.

The modification of a ketone into an aliphatic oxime ether was selected as the most promising candidate for further evaluation. Interestingly, according to the literature, the electronic distribution of aliphatic oxime ether derivatives can mimic that of aromatic groups (Fig. 3). A seminal study from 1985 (Macchia, Balsamo et al. 1985) reported that a series of methyleneaminoxy methyl derivatives (C=NOCH_2_) exhibited *in vitro* activity on β-adrenoceptors comparable to their aromatic analogues. The bioisosteric potential of the oxime ether moiety as a replacement for aryl groups has been extensively reviewed (Patani and LaVoie 1996).

**Fig. 3.**
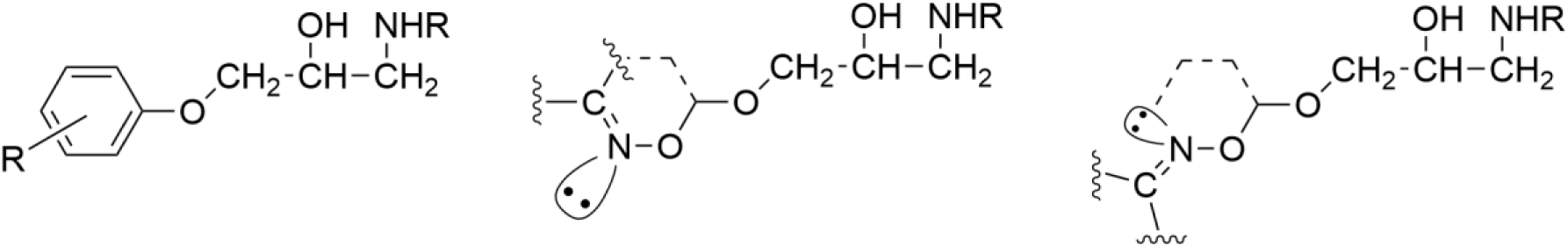
Proposed electronic distribution of aliphatic oxime ethers and their aromatic analogue from Macchia et al (Macchia, Balsamo et al. 1985).

Oximes are widely studied nitrogen- and oxygen-containing structural motifs with diverse biological and pharmacological applications (Dhuguru, Zviagin et al. 2022), including indications related to central nervous system (CNS) disorders, as they are capable of permeating the blood-brain barrier (Lorke, Kalasz et al. 2008, Faiz Norrrahim, Idayu Abdul Razak et al. 2020). Notably, oximes have been extensively investigated as antidotes for organophosphate poisoning, with pyridine-2-aldoxime (pralidoxime) being the only FDA-approved treatment for this condition to date (Dhuguru, Zviagin et al. 2022). Additionally, golexanolone, a novel neurosteroid-based γ-aminobutyric acid receptor (GABA_A_R) antagonist, is currently in development for the treatment of cognitive impairment associated with hepatic encephalopathy (Johansson, Mansson et al. 2018, Montagnese, Lauridsen et al. 2021, Mincheva, Gimenez-Garzo et al. 2022).

In this study, we report the synthesis of C-17 and C-20 oximes and oxime ethers (compounds **11**–**23**) and evaluate their biological activity on recombinant GluN1/GluN2B receptors expressed in human embryonic kidney (HEK293) cells. Additionally, key pharmacokinetic properties were assessed *in vitro*, including stability in rat liver microsomes and parallel artificial membrane permeability (PAMPA). For the most potent compound, **12**, further evaluations were conducted to determine plasma stability, stability in primary rat hepatocytes, and thermodynamic solubility.

## Results and discussion

### Chemistry

The proof-of-concept of our hypothesis of biologically active steroidal oximes and oxime ethers was evaluated on two basic steroidal skeletons. Compounds **11**–**13** were prepared from 3β-hydroxy-pregn-5-en-20-one (pregnenolone) and compounds **14**–**16** (Fig. 4) were prepared from 3β-hydroxy-androst-5-en-17-one (dehydroepiandrosterone, DHEA, **3**).

**Fig. 4.**
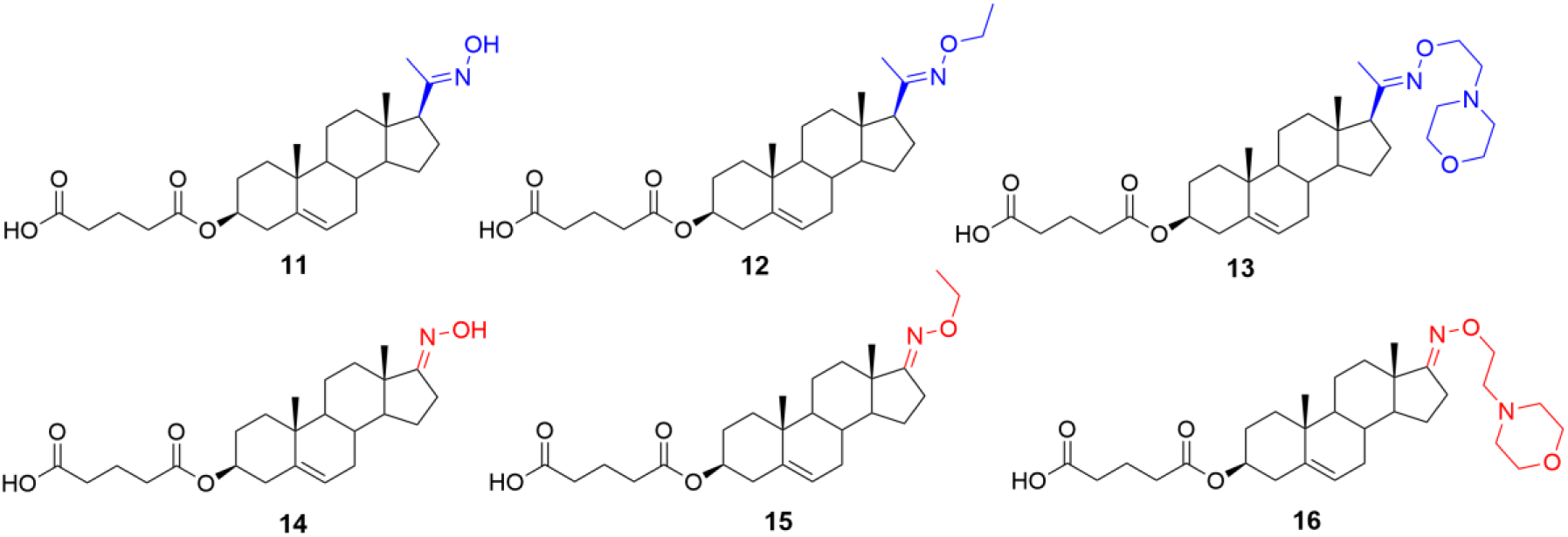
Proof-of-concept molecules **11**–**16**.

A steroidal skeleton with a C-3 hemiglutarate ester moiety was selected based on our previous SAR study on pregn-5-en and androst-5-ene derivatives (Krausova, Slavikova et al. 2018). Oximation at positions C-6 or C-7 is well documented in the literature, as hydroxyimino steroids represent a distinct class of antineoplastic agents (Deive, Rodriguez et al. 2001, Carrasco-Carballo, Guadalupe Hernandez-Linares et al. 2021). Following a review of synthetic strategies for steroidal ketoximes and oxime ethers, we evaluated two methods for introducing the hydroxyimino group at C-20 of pregnenolone (Scheme 1): *(i)* reaction with hydroxylamine hydrochloride in aqueous ethanol in the presence of sodium acetate (NaOAc)(Acharya and Bansal 2014) and *(ii)* reaction with hydroxylamine hydrochloride in ethanolic pyridine or pyridine with triethylamine (Nahar, Sarker et al. 2008, Sikharulidze, Nadaraia et al. 2010, Guthrie, Stein et al. 2012). Both approaches provided comparable isolated yields exceeding 90% (synthesis of compound **24**). The treatment of *O*-alkyl hydroxylamine hydrochloride with sodium acetate was subsequently applied to the synthesis of all target compounds.

Next, we explored two synthetic sequences for the introduction of the hydroxyimino and hemiester moieties (Scheme 1) in the synthesis of compound **12**. This approach was designed to generate a parent compound that could be further modified at C-3 with ester linkers of varying lengths or novel oxime ethers. Compound **12** was synthesized via oximation of the C-20 ketone of pregnenolone using hydroxylamine hydrochloride in aqueous ethanol with NaOAc, yielding compound **24** (98%). This was followed by esterification of the C-3 hydroxyl group with glutaric acid in the presence of EDCI, DMAP, and DIPEA, affording compound **12** in 89% yield.

Alternatively, pregnenolone was first esterified at C-3 with glutaric anhydride and DMAP in pyridine, yielding compound **25** (57%). Subsequent treatment of compound **25** with *O*-ethylhydroxylamine hydrochloride (NH₂OEt·HCl/NaOAc) resulted in compound **12** (64%). Additionally, compound **25** was treated with NH₂OEt·HCl/NaOAc to yield compound **11** (82%), while compound **13** was prepared using freshly synthesized 4-[2-(aminooxy)ethyl]morpholine in 62% yield (Christensen, Erichsen et al. 2013).

**Scheme 1.**
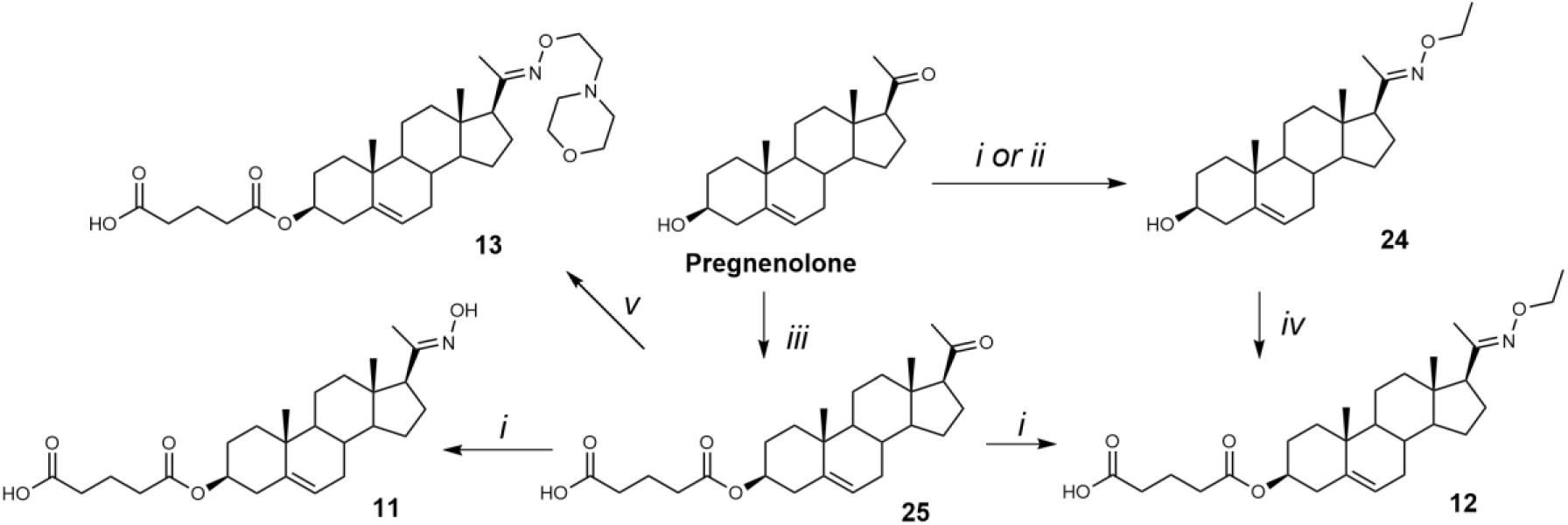
Synthesis of compounds **11**, **12**, and **13**. Reagents and conditions: (i) NH_2_OEt.HCl, NaOAc, EtOH/H_2_O, 95 °C; (ii) NH_2_OEt.HCl, pyridine, Et_3_N, EtOH/H_2_O, 95 °C; (iii) glutaric anhydride, DMAP, pyridine, 105 °C; (iv) glutaric acid, EDCI, DMAP, DIPEA, DCM, rt; (v) 4-[2-(aminooxy)ethyl]morpholine, NaOAc, EtOH/H_2_O, 95 °C.

Since both synthetic approaches yielded comparable isolated yields, we prioritized the sequence involving hemiester formation followed by oxime synthesis for the preparation of compounds **15** and **16** (Scheme 2). This decision was further supported by our previous experience with low-yielding esterification steps in the synthesis of various hemiesters (Krausova, Slavikova et al. 2018). Accordingly, DHEA (**3**) was first esterified with a hemiglutarate moiety at C-3, affording compound **26** in 50% yield. Subsequent oximation with NH₂OEt·HCl/NaOAc yielded compound **14** in 85% yield. Finally, compounds **15** and **16** were synthesized by treating DHEA with NH₂OEt·HCl and 4-[2-(aminooxy)ethyl]morpholine/NaOAc, respectively, yielding compound **15** (52%) and compound **16** (77%).

**Scheme 2.**
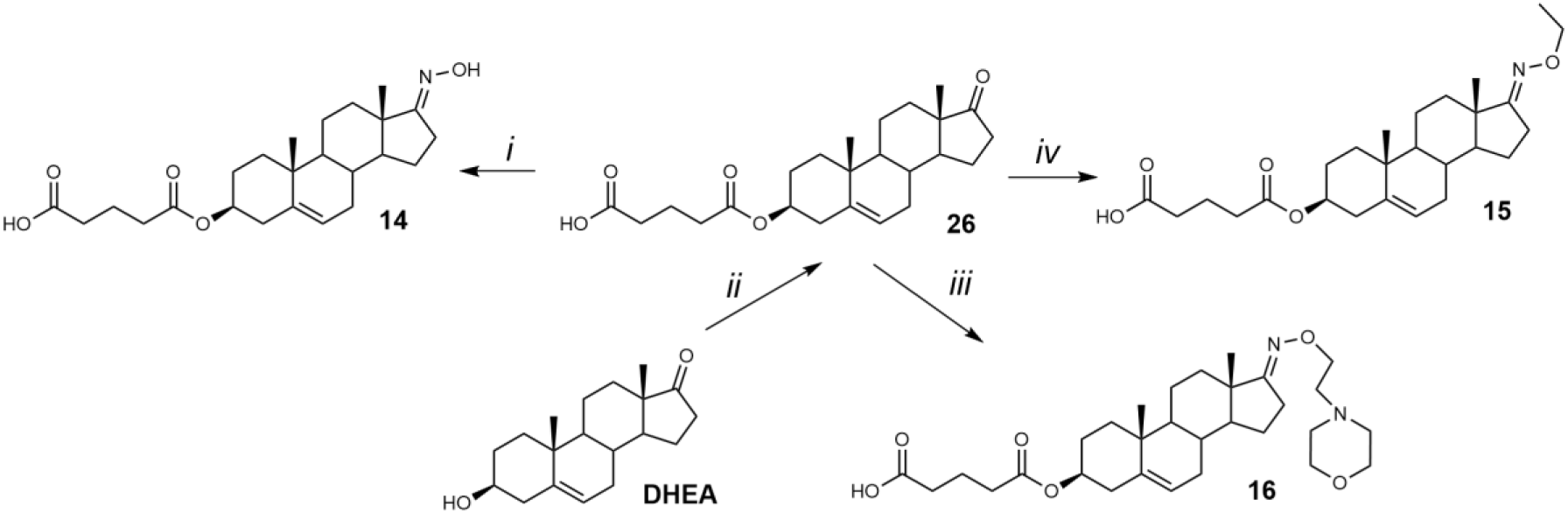
Synthesis of compounds **14**, **15**, and **16**. Reagents and conditions: (i) NH_2_OEt.HCl, NaOAc, EtOH/H_2_O, 95 °C; (ii) glutaric anhydride, DMAP, pyridine, 105 °C; (iii) 4-[2-(aminooxy)ethyl]morpholine, NaOAc, EtOH/H_2_O, 95 °C; (iv) NH_2_O-ethyl.HCl, NaOAc, EtOH/H_2_O, 95 °C.

The biological activity of compounds **11**–**16** (Fig. 4) was evaluated on recombinant GluN1/GluN2B receptors expressed in HEK293 cells. Our results demonstrated that pregnenolone analogues (**11**–**13**) exhibited greater activity than their DHEA-derived counterparts. The reference compound – pregnenolone sulfate - produced a potentiation with a maximal response (E_max_) near baseline (116%), and a potency of EC_50_ = 21.7 μM). Several compounds of series **11**–**16** displayed enhanced efficacy compared to PES. Pregnenolone derivatives **11** and **12** both substantially increased the current amplitude, with **12** showing the greatest potentiation (673 ± 121%) and EC_50_ value of 8.7 μM. DHEA derivative **15** also displayed marked potentiation (441 ± 76%), statistically significant, but with lower potency (EC_50_ = 22.7 μM), aligning more closely with **PES**. Consequently, further compound development focused exclusively on modifying the pregnenolone skeleton. Based on these findings, we designed a series of pregnenolone oxime ethers **17**–**21** (Fig. 5).

**Fig. 5.**
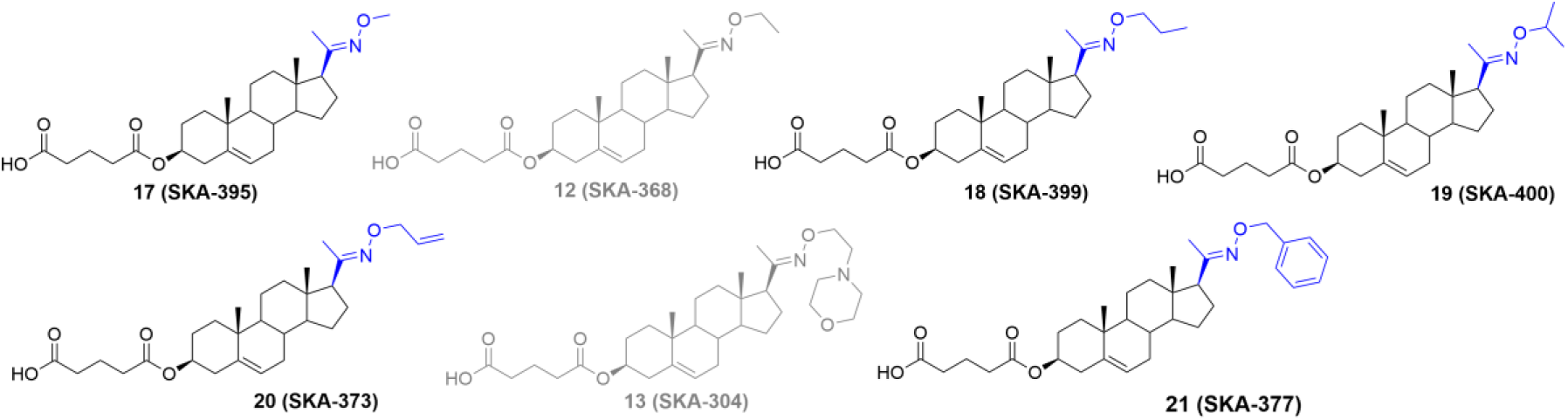
A series of oxime ethers modifications of pregnenolone 3-hemiglutarate **12**, **13**, and **17**–**21**.

Compounds **17**–**21** were synthesized by reacting pregnenolone 3-hemiglutarate (**25**) with the corresponding oxime reagent in the presence of NaOAc (Scheme 3). Compound **17** was obtained using *O*-methyl hydroxylamine hydrochloride (NH₂OCH₃·HCl) in 73% yield, compound **18** with *O*-propyl hydroxylamine hydrochloride (NH₂OCH₂CH₃·HCl) in 55% yield, and compound **19** with *O*-isopropyl hydroxylamine hydrochloride (NH₂OCH(CH₃)₂·HCl) in 49% yield. Treatment with *O*-allyl hydroxylamine hydrochloride (NH₂OCH₂CH=CH₂·HCl) afforded compound **20** in 63% yield, while reaction with *O*-benzyl hydroxylamine hydrochloride (NH₂OCH₂C₆H₅·HCl) produced compound **21** in 89% yield.

**Scheme 3.**
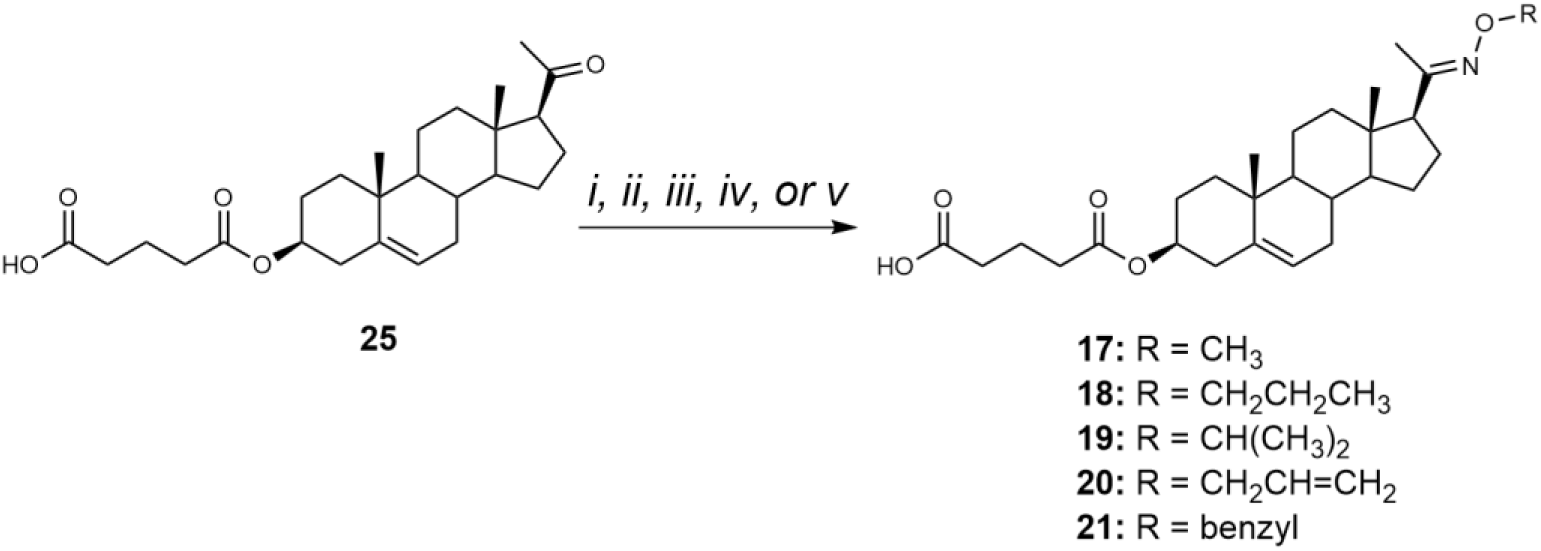
Synthesis of compounds **17**–**21**. Reagents and conditions: (i) For **17**: NH_2_O-methyl.HCl, NaOAc, EtOH/H_2_O, 95 °C; (ii) For **18**: NH_2_O-propyl.HCl, NaOAc, EtOH/H_2_O, 95 °C; (iii) For **19**: NH_2_O-isopropyl.HCl, NaOAc, EtOH/H_2_O, 95 °C; (iv) For **20**: NH_2_O-allyl.HCl, NaOAc, EtOH/H_2_O, 95 °C; (v) For **21**: NH_2_O-benzyl.HCl, NaOAc, EtOH/H_2_O, 95 °C.

The biological activity of compounds **17**–**21** (Fig. 5) was evaluated on recombinant GluN1/GluN2B receptors expressed in HEK293 cells. Our results showed that the structural modifications in compounds **17**–**21** did not enhance the PAM effect compared to compound **12**, yet compound **17** showed the greatest potentiation (503 ± 85%) and EC_50_ value of 6.1 μM. Consequently, we synthesized analogues of compound **12** with hemiester linkers of varying lengths (Scheme 4).

**Scheme 4.**
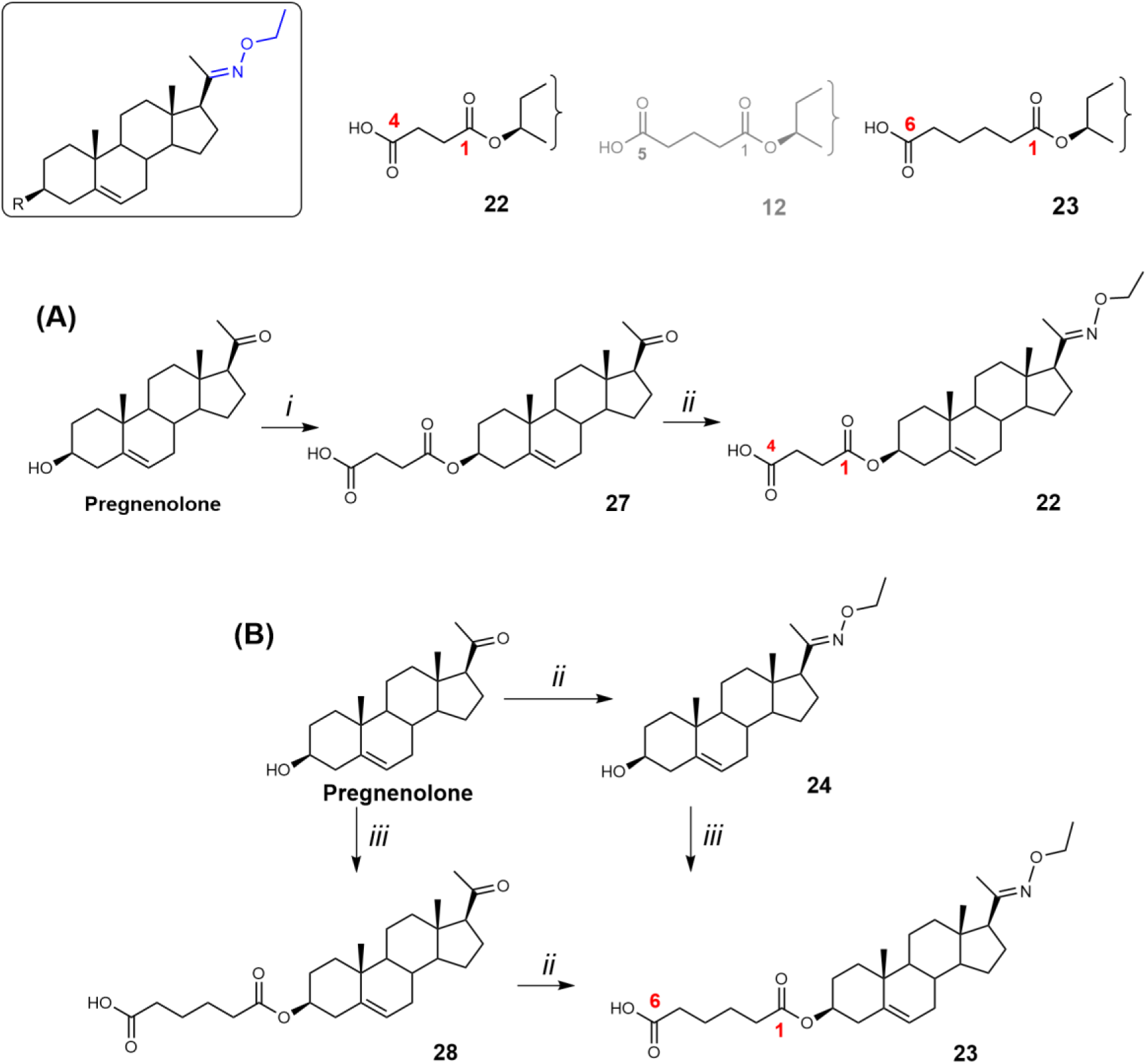
Synthesis of compounds **22** and **23**. Reagents and conditions: (i) succinic acid, EDCI, DMAP, DIPEA, DCM, rt; (ii) NH_2_-ethyl.HCl, NaOAc, EtOH/H_2_O, 95 °C; (iii) adipic acid, EDCI, DMAP, DIPEA, DCM, rt.

The C-3 hemisuccinate (compound **22**) was prepared via a two-step synthesis (Scheme 4). First, the C-3 hydroxy group of pregnenolone was esterified with succinic acid using EDCI, DMAP, and DIPEA, affording compound **27** in 57% yield. Subsequent oximation with NH₂OEt·HCl/NaOAc yielded compound **22** in 71% yield. The C-3 hemiadipate (compound **23**) was synthesized analogously. Pregnenolone was esterified with adipic acid in the presence of EDCI, DMAP, and DIPEA, giving compound **28** in a yield of 35%. Similarly, an attempt to esterify compound **24** yielded only 13% of compound **23**, with 40% of the starting material recovered. These low esterification yields are consistent with our previously published results (Krausova, Slavikova et al. 2018).

Finally, the biological activity of compounds **22** and **23** was evaluated on recombinant GluN1/GluN2B receptors expressed in HEK293 cells. Our results showed that the structural modifications in compounds **22** and **23** did not enhance the PAM effect compared to compound **12**.

**Fig. 6.**
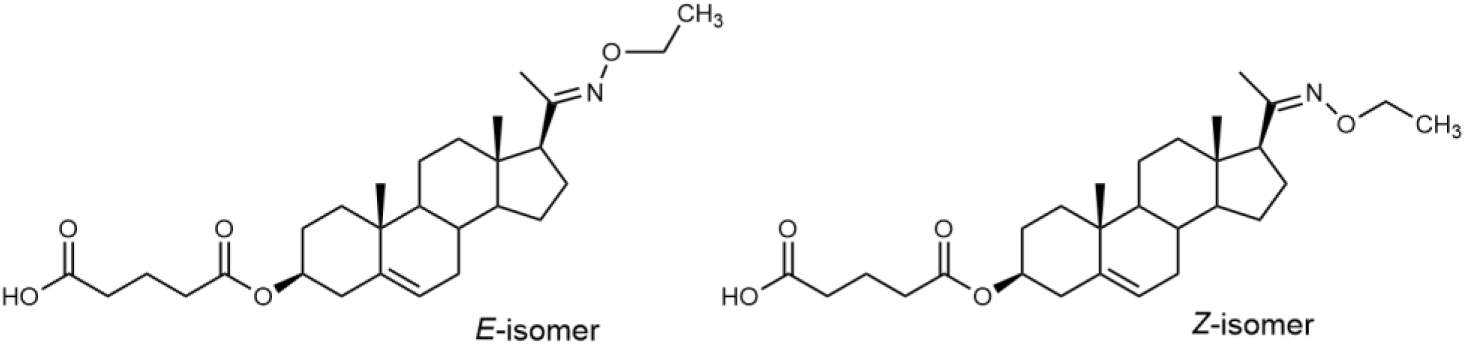
*E*- and *Z*-isomers of compound **12**.

To unambiguously assign the structure of compound **12** which could exist in interconvertible *E*- and *Z*-stereoisomerism (Fig. 6), we have performed the structural assignment of the proton and carbon signals by combining 1D-^1^H and ^13^C spectra with homonuclear 2D-H,H-COSY and 2D-H,H-ROESY, and heteronuclear 2D-H,C-HSQC and 2D-H,C-HMBC spectra on a Bruker 600 AVANCE III instrument (^1^H at 600.13 MHz and ^13^C at 150.9 MHz) with 5 mm cryoprobe in CDCl_3_ solutions at 25 °C. Experimental evidence for configuration on the C(20)=N double bond based on NOE contacts of CH_2_ or CH_3_ protons was not successful, likely due to the long distance of these protons to either C(21)H_3_ in the *E*-isomer or C(16)H and C(17)H_2_ in the *Z*-isomer. The geometry optimization on the model of both isomers suggested a slightly lower potential energy for the E-isomer, however, this difference is insufficient to serve as definitive evidence of configuration. The ^1^H and ^13^C NMR data are in Table 1.

**Table 1.** The ^13^C and ^1^H NMR data of compound 12. Coupling constants J(H,H) are given in brackets.

| Position | Type | Carbon | Proton |
| --- | --- | --- | --- |
| **1** | -CH<sub>2</sub>- | 36.96 | 1.865; 1.145 |
| **2** | -CH<sub>2</sub>- | 27.74 | 1.855; 1.585 |
| **3** | >CH-O | 74.22 | 4.62 m |
| **4** | -CH<sub>2</sub>- | 28.08 | 2.31 (2H) |
| **5** | >C= | 139.60 | N/A |
| **6** | =CH- | 122.50 | 5.375 m |
| **7** | -CH<sub>2</sub>- | 31.77 | 2.00; 1.57 |
| **8** | >CH- | 31.99 | 1.465 |
| **9** | >CH- | 50.04 | 0.995 |
| **10** | >C< | 36.62 | N/A |
| **11** | -CH<sub>2</sub>- | 20.97 | 1.57; 1.445 |
| **12** | -CH<sub>2</sub>- | 38.56 | 1.88; 1.30 |
| **13** | >C< | 43.70 | N/A |
| **14** | >CH- | 56.12 | 1.11 |
| **15** | -CH<sub>2</sub>- | 24.27 | 1.675; 1.205 |
| **16** | -CH<sub>2</sub>- | 23.16 | 2.16; 1.67 |
| **18** | -CH<sub>3</sub> | 13.19 | 0.64 |
| **19** | -CH<sub>3</sub> | 19.32 | 1.02 |
| **20** | >C=N | 157.40 | N/A |
| **21** | -CH<sub>3</sub> | 15.78 | 1.82 |
| **N-CH<sub>2</sub>-CH<sub>3</sub>** |  |  |  |
|  | O-CH<sub>2</sub>- | 68.72 | 4.085 |
|  | -CH<sub>3</sub> | 14.74 | 1.24 |
| **3-O-CO-CH<sub>2</sub>-CH<sub>2</sub>-CH<sub>2</sub>-CH<sub>3</sub>** |  |  |  |
|  | C=O | 172.27 | N/A |
|  | -CH<sub>2</sub>- | 33.50 | 2.37 (2H) |
|  | -CH<sub>2</sub>- | 19.87 | 1.95 (2H) |
|  | -CH<sub>2</sub>- | 32.86 | 2.43 (2H) |
|  | COOH | 178.27 | N/A |

### Effects of compounds 11–23 on responses of GluN1/GluN2B receptors in HEK293 cells to glutamate

To investigate the effect of compounds **11**–**23** on the activity of NMDARs, electrophysiological measurements were performed on HEK293 cells that were co-transfected with cDNAs containing genes encoding for the rat GluN1-1a and GluN2B subunits. The degree of modulation (E; potentiation/ inhibition) for the newly synthesized compounds was determined using the following formula:

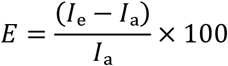

where *I_e_* is the value of the current amplitude during glutamate and compound coapplication and *I_a_* is the current amplitude value for glutamate application. The degree of potentiation (%) was determined for five compound doses, differing 100-fold in their concentration range, in individual cells, and data were fitted to the following equation:

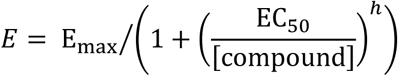

where E_max_ is the maximal value of potentiation induced by a saturating concentration of the compound, EC_50_ is the concentration of the compound that produces half-maximal potentiation of the agonist-evoked current, [compound] is the compound concentration, and *h* is the apparent Hill coefficient. The results are presented as mean ± standard error of the mean (SEM), with *n* equal to the number of independent measurements. One-way analysis of variance (ANOVA) was used for multiple comparisons (unless otherwise stated, a value of *P ≤ 0.05* was used for the determination of significance).

The overview of the concentration-dependent effects of compounds **11**–**23** on responses of GluN1/GluN2B receptors in HEK293 cells to glutamate are summarized in Table 2.

**Table 2.** Parameters from the concentration-response analysis of the effect of compounds 11–23 on glutamate-induced currents of GluN1/GluN2B receptors expressed in HEK293 cells.

| No. | E<sub>max</sub> ± SEM (%) | EC<sub>50</sub> ± SEM (μM) | h ± SEM | n |
| --- | --- | --- | --- | --- |
| PES | 116 ± 10 | 21.7 ± 1.6 | 1.5 ± 0.1 | 10 |
| 11 | 270 ± 35 | 7.9 ± 2.1* | 1.5 ± 0.1 | 5 |
| 12 | 673 ± 121*** | 8.7 ± 1.1 | 1.7 ± 0.3 | 7 |
| 13 | 111 ± 11 | 34.7 ± 4.2 | 2.8 ± 0.2** | 5 |
| 14 | N/A# | N/A# | N/A# | 7 |
| 15 | 441 ± 76** | 22.7 ± 3.1 | 1.5 ± 0.1 | 7 |
| 16 | 44 ± 5 | 63.5 ± 6.4 | 1.7 ± 0.1 | 4 |
| 17 | 503 ± 85** | 6.1 ± 0.7* | 1.6 ± 0.1 | 4 |
| 18 | NAM (7 ± 2%) at 3 μM.<br>PAM (75 ± 21% and 108 ± 17%) at 10 and 30 μM |  | N/A | 4 |
| 19 | NAM (27 ± 5%) at 3 μM.<br>PAM (81 ± 20% and 169 ± 55%) at 10 and 30 μM |  | N/A | 5 |
| 20 | 317 ± 43 | 12.6 ± 2.9 | 2.0 ± 0.2 | 5 |
| 21 | 403 ± 56 | 11.3 ± 2.0 | 1.7 ± 0.3 | 4 |
| 22 | 340 ± 22 | 7.0 ± 1.1* | 1.3 ± 0.1 | 7 |
| 23 | 451 ± 79* | 7.5 ± 1.1 | 1.7 ± 0.1 | 5 |
| One-way ANOVA | P < 0.001 | P < 0.001 | P = 0.013 |  |

PAM, positive allosteric modulator; NAM, negative allosteric modulator; NA, not analysed; #Compound **14** had no effect at the concentration range of 3-100 μM and was toxic for the HEK293 cells which precluded the analysis. One-way ANOVA on ranks with Kruskal-Wallis *post hoc* analysis versus PES; \*P>0.05; \*\*P>0.01; \*\*\*P>0.001.

We have evaluated the relationship between oxime derivatives **11**-**23** and their modulatory effect on recombinant GluN1/GluN2B receptors. Due to the disuse-dependent effect of neurosteroid-based PAMs at NMDARs (Horak, Vlcek et al. 2004, Krausova, Slavikova et al. 2018). GluN1/GluN2B receptors were activated by 1 µM glutamate, a concentration corresponding to the EC_50_ value for this agonist, to reveal the maximal modulatory effect of the tested compounds during coapplication. We focused on GluN2B-containing receptors as we previously demonstrated that pregnane-based steroids with PAM activity can compensate for NMDAR hypofunction caused by the *de novo* missense variant L825V in GluN2B subunit (Kysilov, Hrcka Krausova et al. 2022).

Our data shows that 8 out of 13 newly synthesized compounds had PAM effect at GluN1/GluN2B receptors, with increased efficacy (in case of compounds **12**, **15**, **17**, and **23** significantly) and/or potency (in case of compounds **11**, **17**, and **22** significantly) as compared to endogenous sulfate analogue PES. Three compounds – **14**, **18**, and **19** – were not identified as PAMs of NMDARs. In contrast, compounds **14**, **18**, and **19** exhibited a more complex concentration-dependent pharmacological profile, acting as PAMs at certain concentrations while displaying antagonistic effects at others. This behavior is less common but has been observed for some compounds. For example, several ligand-gated ion channels are modulated by ivermectin, including the glutamate-gated chloride channel, GABA_A_R, glycine receptor, α7-nicotinic acetylcholine receptor, and P2X4 purinergic receptor (Zemkova, Tvrdonova et al. 2014). Ivermectin enhances the activity of these channels in a concentration-dependent manner; at lower concentrations, it potentiates agonist-induced responses, whereas at higher concentrations, it can activate the channels independently of the agonist. Concentration dependence has been also shown for neurosteroids. For example, the U-shaped dose-response curve on the normalized peak amplitude of I_GABA_ has been shown the Purkinje cells of cerebellum. In particular, allopregnanolone, pregnanolone, and PAS enhanced I_GABA_ between 10 and 5000 nM. This effect was reversed at higher concentrations from 10 to 100 µM (Bukanova, Solntseva et al. 2021, Solntseva, Bukanova et al. 2022).

As mentioned above, within the structural family of this study, the majority of compounds exhibited PAM effect on NMDARs with more than 16-fold difference in the E_max_ values (44% for the least efficacious compound **16** compared to 673% for the most efficacious compound 12), with EC_50_ values varying from 6.1 μM for the most potent compound 17 to 63.5 μM for the least potent compound **16**.

First, we compared the biological activity of C-17 versus C-20 oxime derivatives using recombinant GluN1/GluN2B receptors expressed in HEK293 cells. Our findings revealed that pregnenolone-based oximation displayed overall high PAM-activity. Compound **12** emerged as the most efficacious compound in the series. In contrast, the C-17 hydroxy oxime analogue of DHEA (**14**) lost its PAM-activity and the morpholine-substituted analogue **16** proved to be the least effective derivative in the study. Next, we explored the replacement of the pregnenolone C-20 ketone group with various oxime moieties. As oximation represents an effective strategy for SAR studies, a wide range of reagents is readily available for this purpose. We selected and prepared a series of aliphatic and branched alk(en)yl oximes (**17**– **20**), along with benzyl (**21**) analogue, and morpholine analogues (**13**), in which the substituent is linked to the pregnenolone scaffold via the oxime functionality. Interestingly, within the series of alk(en)yl oximes (**12**, **13**, **17**–**20**), the methyl (**17**) and ethyl (**12**) oximes emerged as the most efficacious compounds, exhibiting E_max_ values of 673% and 503%, respectively. The corresponding EC₅₀ values were 6.1 µM for compound **17** and 8.7 µM for compound **12**. In contrast, the propyl (**18**) and isopropyl (**19**) analogues displayed a complex, concentration-dependent activity profile. Both compounds potentiated agonist-induced responses at 3 µM, but at higher concentrations (10–30 µM), their effects shifted toward inhibition. The presence of a double bond within the oxime moiety in compounds 20 (allyl) and **21** (benzyl) resulted in comparable efficacy, with E_max_ values of 317% and 403%, respectively. The corresponding EC₅₀ values were 12.6 µM for compound 20 and 11.3 µM for compound **21**. Our SAR study revealed that modifications of the oxime moiety did not result in improved efficacy compared to compound **12**. Consequently, we turned our attention to the C-3 hemiester moiety. Compounds **22** (hemisuccinate) and **23** (hemiadipate) were evaluated for their ability to modulate NMDARs. Both compounds demonstrated comparable efficacy, with E_max_ values of 340% and 451%, respectively. The corresponding EC₅₀ values were 7.0 µM for compound **22** and 7.5 µM for compound **23**. Compound **12** was identified as the most efficacious compound of the study (Fig. 7).

**Fig. 7.**
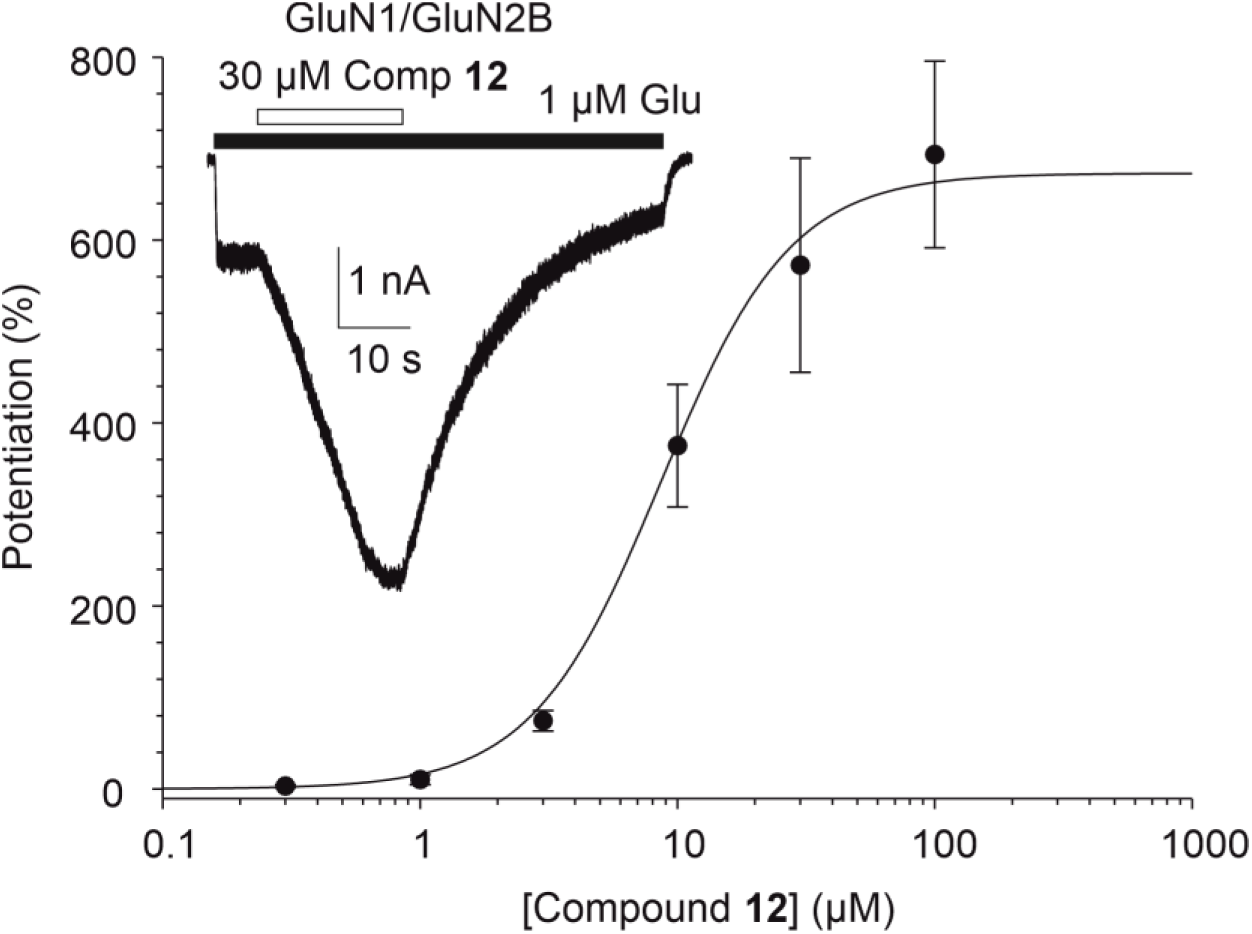
The effect of compound 12 on GluN1/GluN2B receptors. The graph shows the concentration-response curve for the effect of compound **12** at GluN1/GluN2B receptors. Data points are averaged values of potentiation at given concentration (0.3–100 µM) from 7 independent measurements; error bars represent SEM. The degree of potentiation of glutamate-induced responses recorded in the presence of compound **12** was determined in individual cells and data were fitted to the logistic equation (obtained parameters for compound **12** are indicated in Table 2). Inset shows an example of trace recorded from HEK293 cell expressing recombinant GluN1/GluN2B receptors. Compound 12 (30 µM) was applied simultaneously with 1 µM glutamate and 30 µM glycine (the duration of compound **12** and glutamate application is indicated by open and filled bars, respectively).

### In Vitro ADME Profiling – Metabolic Stability, Permeability Assessment, and Solubility

The results of *in vitro* ADME profiling are summarized in Table 3. First, the metabolic stability of compounds **11**–**23** was assessed using rat liver microsomes. Briefly, each compound was incubated with microsomes for 60 min, followed by extraction and analysis via LC-MS. Verapamil was used as a control due to its well-characterized metabolic elimination and relatively short half-life (Follath, Ha et al. 1986). Remarkably, nearly all tested compounds demonstrated high metabolic stability, with half-lives exceeding 60 min. Notably, compounds **12**, **19**–**21**, and **23** exhibited exceptional stability, showing no detectable metabolism after 60 min of incubation.

**Table 3.** ADME Assessment of compounds 11–23.

| Steroid | Stability in rat microsomes |  | PAMPA permeability |  | Solubility | Stability in rat plasma (% remaining) |  | Stability in primary rat hepatocytes |  |
| --- | --- | --- | --- | --- | --- | --- | --- | --- | --- |
|  | t<sub>1/2</sub> (min) | Cl<sub>int</sub> (μl/min/mg) | P<sub>e</sub> (cm/s) | Retention (%) | PBS, pH 7.4 (μM) | 8 h | 24 h | t<sub>1/2</sub> (min) | Cl<sub>int</sub> (μl/min/mg) |
| **11** | 58.6 ± 18.7 | 23.6 | 1.55E-04 | 87 | - | 69.9 ± 6.0 | 58.1 ± 8.1 |  |  |
| **12** | >> 60<sup>a</sup> | - | 3.89E-06 | 97 | 361 ± 41 | 85.6 ± 1.0 | 59.6 ± 0.7 | 72.4 | 19.1 |
| **13** | > 60 | - | n.d. | 95 | - | - | - | - | - |
| **14** | > 60 | - | 2.63E-05 | 17 | - | 61.1 ± 1.6 | 56.3 ± 2.5 | - | - |
| **15** | > 60 | - | 1.82E-03 | 80 | - | 82.1 ± 7.5 | 69.9 ± 5.0 | - | - |
| **16** | > 60 | - | 6.76E-05 | 17 | - | 82.1 ± 5.8 | 73.1 ± 7.8 | - | - |
| **17** | > 60 | - | 5.50E-06 | 95 | - | - | - | - | - |
| **18** | > 60 | - | n.d. | 100 | - | - | - | - | - |
| **19** | >> 60<sup>a</sup> | - | n.d. | 99 | - | - | - | - | - |
| **20** | >> 60<sup>a</sup> | - | 7.24E-06 | 99 | - | - | - | - | - |
| **21** | >> 60<sup>a</sup> | - | 1.82E-06 | 95 | - | - | - | - | - |
| **22** | 52.6 ± 3.7 | 26.3 | 1.58E-06 | 96 | 22 ± 1 | 94.7 ± 5.7 | 76.7 ± 4.7 | - | - |
| **23** | >> 60 | - | n.d. | 100 | 21 ± 1 | 85.7 ± 4.6 | 56.3 ± 3.0 | - | - |
| **Verapamil** | 43.9 ± 1.2 | 31.5 | 3.72E-04 | 33 | - | - | - | - | - |
| **Atenolol** | - | - | 1.07E-07 | 31 | - | - | - | - | - |
| **Imipramine** | - | - | - | - | - | - | - | 20.6 | 67.3 |

<sup>a</sup>The compound exhibits a half-life significantly exceeding 60 min, with no detectable decrease in intensity over time. Where referred, results are presented as mean ± SEM; n.d., not defined.

Second, the permeability of compounds **11**–**23** was evaluated using the parallel artificial membrane permeability assay (PAMPA), a widely accepted method for predicting brain penetration (Kansy, Avdeef et al. 2004, Avdeef, Bendels et al. 2007, Carrara, Reali et al. 2007, Morofuji and Nakagawa 2020). Atenolol and verapamil served as reference compounds for low and high permeability, respectively (Teksin, Seo et al. 2010). Permeability was quantified using permeability coefficients (P_e_) and mass retention, the latter representing the proportion of compound retained in the membrane. Compounds **11** and **15** exhibited permeability comparable or greater to that of verapamil, compounds **14** and **16** exhibited moderate permeability. Among the series, compound **15** was identified as the most permeable. High mass retention values (>80%) were observed for most compounds, indicating strong membrane interaction, with the exception of compounds **14** and **16**, which showed high permeability but low mass retention (17%). These findings suggest that structural modifications of the DHEA scaffold can yield derivatives with favourable permeability profiles. However, it is important to note that compounds **14** and **16** were inactive *in vitro* despite their favourable permeability characteristics.

Next, we evaluated the stability in rat plasma and thermodynamic solubility in PBS (pH 7.4) for proof-of-concept compounds (compounds **11**, **12**, **14**–**16**), as well as for the most potent compound at NMDAR-induced currents (**12**) and its analogues (**22** and **23**). All compounds showed comparable stability within 8 and 24 h. Compound **12** also demonstrated moderate metabolic stability in rat hepatocytes. Overall, the results for rat plasma stability and hepatic clearance were consistent with those observed in rat microsomes. Notably, the tested compounds exhibited unexpected stability of the hemiester bond after 8 h, suggesting that these compounds could potentially be suitable for preliminary *in vivo* studies.

Finally, to test our hypothesis that introducing an oxime ether moiety could enhance the drug-like properties of our compounds, we compared the rat plasma stability of compounds **12**, **22**, and **23** with their corresponding 20-ketone analogues (Krausova, Slavikova et al. 2018). The results, summarized in Table 4, show that the oxime ether moiety indeed improved the plasma stability of these compounds.

**Table 4.**
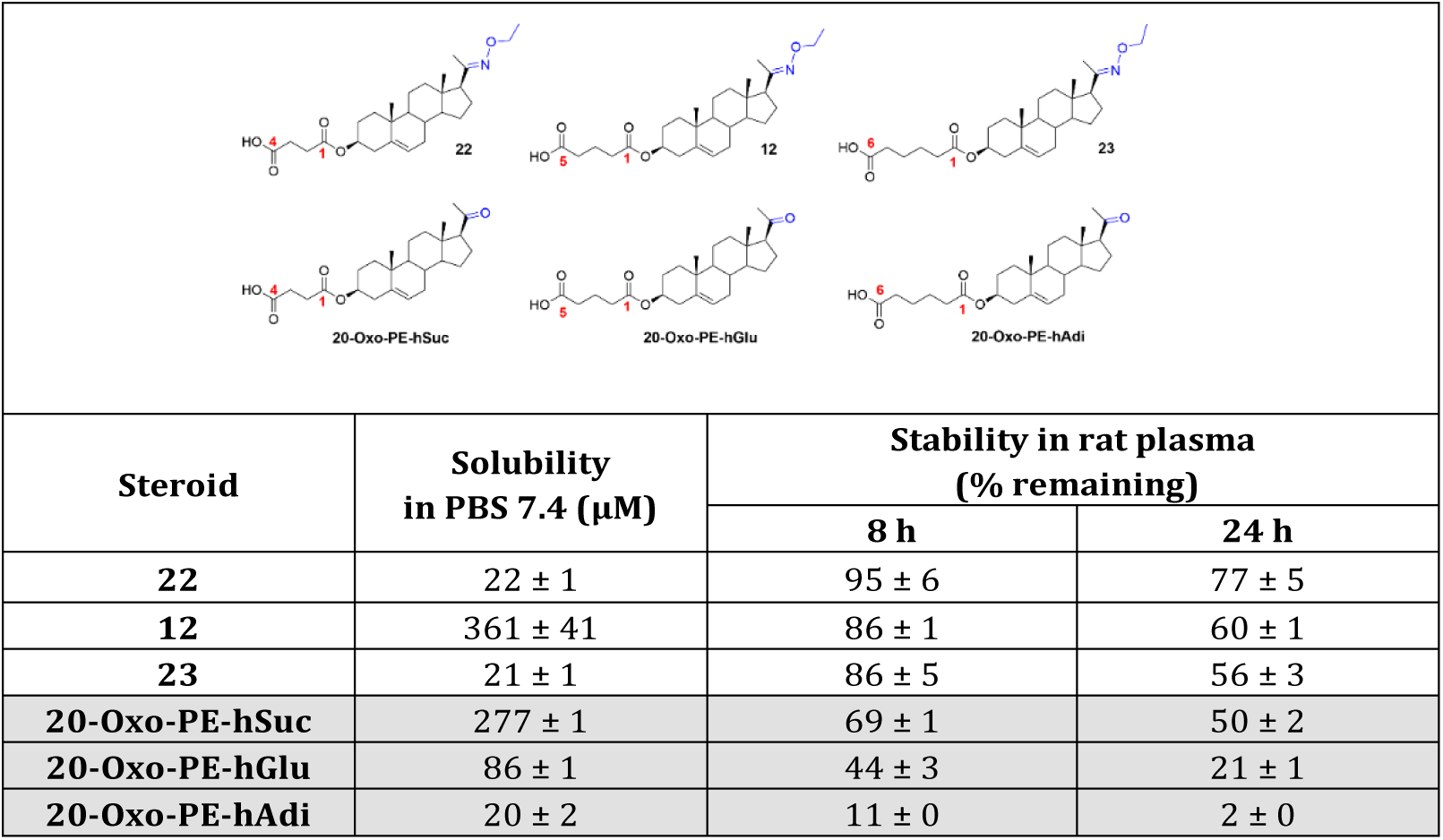
Comparison of the solubility and rat plasma stability of oxime ethers 12, 22, and 23 with that of their corresponding 20-ketone analogues.

|  |  |  |  |
| --- | --- | --- | --- |
| Steroid | Solubility<br>in PBS 7.4 (μM) | Stability in rat plasma<br>(% remaining) |  |
|  |  | 8 h | 24 h |
| **22** | 22 ± 1 | 95 ± 6 | 77 ± 5 |
| **12** | 361 ± 41 | 86 ± 1 | 60 ± 1 |
| **23** | 21 ± 1 | 86 ± 5 | 56 ± 3 |
| **20-Oxo-PE-hSuc** | 277 ± 1 | 69 ± 1 | 50 ± 2 |
| **20-Oxo-PE-hGlu** | 86 ± 1 | 44 ± 3 | 21 ± 1 |
| **20-Oxo-PE-hAdi** | 20 ± 2 | 11 ± 0 | 2 ± 0 |

We evaluated compounds **11**–**23** in a series of *in vitro* assays designed to assess key parameters relevant to their bioavailability profiles. Our results indicate that substitution of the steroidal scaffold with an oxime or oxime ether moiety yielded compounds with unexpectedly high stability in both rat microsomes and rat plasma, despite the presence of a hemiester group at the C-3 position. This is particularly notable, as the hemiester moiety is generally considered susceptible to rapid hydrolysis by carboxylesterases, as reported in the literature (Schottler and Krisch 1974). Furthermore, our experiments demonstrated that certain lipophilic steroidal derivatives (compound **12**) can demonstrate excellent thermodynamic solubility (>350 µM) atypical for steroidal compounds.

## Experimental

### General

Dichloromethane was refluxed over phosphorous pentoxide under inert atmosphere for 30 min followed by distillation. The chemicals and reagents were purchased from commercial suppliers and used without further purification. Melting points were determined with a micromelting point apparatus (Hund/Wetzlar, Germany) and are uncorrected. For elemental analysis, PE 2400 Series II CHNS/O Analyzer (Perkin Elmer, MA, USA) was used, with microbalance MX5 (Met-tler Toledo, Switzerland). For measurement of optical rotation, AUTOPOL IV (Rudolph Research Analytical, NJ, USA) was used, all samples were measured at 20 °C, at a given concentration in a given solvent at 589 nm. Analytical samples were dried over phosphorus pentoxide at 50 °C/100 Pa. The HRMS spectra were performed with LTQ Orbitrap XL (Thermo Fischer Scientific, MA, USA) in ESI mode. Thin-layer chromatography (TLC) was performed on silica gel (Merck, 60 µm). Proton and carbon NMR spectra were measured in a Bruker AVANCE-400 FT NMR spectrometer (400 MHz, 101 MHz) in CDCl_3_, with tetramethylsilane as the internal standard. Chemical shifts are given in ppm (δ scale). Coupling constants (J) and widths of multiplets (W) are given in Hz. IR spectra were recorded on a Nicolet 6700 spectrometer (wavenumbers in cm-1). MS High-resolution MS spectra were performed with a Q-Tof microspectrometer (Waters). LC-MS was measured by two methods. Method A: using LC-MS system (C4 column Phenomenex 250 mm length, 2 mm width, Jupiter 300-5 C4, TSP quaternary pump P4000, TSP autosampler AS3000) equipped with UV/VIS detector (UV6000LP, UV/VIS setup wavelength range 190-700 nm, bandwidth 1 nm, step 1 nm, and scan rate 1 Hz) and ion-trap mass spectrometer Advantage (positive ion mode with m/z range from 220 to 1000 Da). Solvent A was 98% water/ 2% acetonitrile and B was 95% acetonitrile/ 3% isopropanol/ 2% water with 5mM ammonium formate both. Gradient setup: 0-25-30-30.1-45min 50-100-100-50-50% solvent B and flow rate 150 µL/min. Method B: Analysis was carried out on the UHPLC-MS system Nexera LC-40 (Shimadzu, Japan) equipped with ELS, and MS detectors. Ions were detected: MS, positive DUIS (ESI/APCI) ion mode, with m/z range from 250 to 1000 Da, interface temp. 350 °C, desolvation line temp. 250 °C, nebulizing gas flow 1.5 L/min, heat block temp. 400 °C, drying gas 15 L/min. ELSD, desolvation temp. 60°C, gain 10. Solvent A was water/methanol/formic acid (950:50:1), and solvent B was acetonitrile. Analysis was performed in an isocratic mode with 80% of solvent B, and a flow rate of 0.5 mL/min, column: Shim-pack Scepter, C8-120, 1.9 µm, 100 × 2.1 mm (Shimadzu). The sample was prepared by dissolving the material (1 mg) in methanol (1 mL, LCMS grade) and then sonicated for 5 min. Injection volume varied from 0.1 to 0.4 µL. The percentage purity of compounds was calculated from the ratio of peaks in the ELSD chromatogram. The purity of the final compounds was assessed by a combination of NMR and based on LC-HR-MS analysis or elemental analysis, and the results showed they were greater than 95%.

### Chemistry

General procedure: Synthesis of oxime ethers with O-alkyl hydroxylamine hydrochloride and sodium acetate. A mixture of *O*-alkyl hydroxylamine hydrochloride (0.58 mmol), sodium acetate (NaOAc) (1.05 mmol), and steroid (0.23 mmol) was dissolved in an aqueous ethanol solution (EtOH/H₂O, 10:1, 20 mL). The reaction mixture was heated at 95 °C, and the progress of the reaction was monitored periodically by TLC. If the conversion was low after 24 hours, additional O-alkyl hydroxylamine hydrochloride and NaOAc were added, and the mixture was heated at 95 °C for a further 24 hours. After cooling to room temperature, the reaction mixture was poured into water (50 mL) and extracted with dichloromethane (3 × 50 mL). The combined organic layers were washed with brine (50 mL), dried over Na₂SO₄, and concentrated under reduced pressure. The crude product was purified by column chromatography.

### Synthesis of compounds 11–28

#### 20-Oxo-pregn-5-en-3β-yl hemiglutarate O-methyloxime (**11**)

Compound **11** was prepared according to the *General procedure*. Starting from compound **25** (150 mg, 0.35 mmol), compound **11** (129 mg, 82%) was obtained as by column chromatography (5-10% methanol in dichloromethane). Recrystallization from acetone/heptane afforded LC-MS purity of 99%. Mp 185–186 °C. [α]_D_ –13.6 (*c* 0.2, CHCl_3_). ^1^H NMR (400 MHz, CDCl_3_): δ 0.62 (s, 3H, H-18), 1.01 (s, 3H, H-19), 1.94 (s, 3H, H-21), 1.96–1.99 (m, 2H, HOOC-CH_2_-<u>CH_2_</u>-CH_2_), 2.38 (t, *J* = 7.1 Hz, 2H, HOOC-<u>CH_2_</u>-CH_2_-CH_2_), 2.43 (t, *J* = 7.0 Hz, 2H, HOOC-CH_2_-CH_2_-<u>CH_2_</u>), 4.60–4.71 (m, 1H, H-3), 5.37 (d, *J* = 5.8 Hz, 1H, H-5). ^13^C NMR (101 MHz, CDCl_3_): δ 178.21 (COOH), 172.50 (ester CO), 159.49 (C=N-OH), 139.73 (C-5), 122.57 (C-6), 73.79 (C-3), 56.74, 56.02, 50.02, 44.37, 38.49, 38.32, 37.08, 36.81, 33.38, 32.72, 32.22, 31.90, 27.85, 24.43, 23.44, 21.08, 20.30, 19.56, 16.00, 13.38. IR spectrum (CHCl_3_): 3591, 3274 (oxime OH), 3510, 3097 (COOH), 1723 (C=O), 1655 (C=C). MS (negative ESI): *m/z* 445.3 (35%, M), 444.3 (100%, M–H). HR-MS (negative ESI) *m/z*: For C_26_H_38_O_5_N [M–H] calcd, 444.2755; found, 444.2753.

#### 20-Oxo-pregn-5-en-3β-yl hemiglutarate O-ethyloxime (**12**)

Compound **12** was prepared according to *General procedure*. Starting from compound **25** (100 mg, 0.23 mmol), compound **12** (98 mg, 89%) was obtained as an oily compound by column chromatography (10-20% acetone in dichloromethane) with LC-MS purity of was 99%. [α]_D_ –13.7 (*c* 0.3, CHCl_3_). ^1^H NMR (400 MHz, CDCl_3_): δ 0.64 (s, 3H, H-18), 1.02 (s, 3H, H-19), 1.22 (t, 3H, *J* = 7.0 Hz, O-CH_2_-<u>CH_3_</u>), 1.81 (s, 3H, H-21), 1.93–1.98 (m, 2H, HOOC-CH_2_-<u>CH_2_</u>-CH_2_), 2.18 (s, 3H, O-CH_2_-<u>CH_3_</u>), 2.37 (t, *J* = 7.3 Hz, 2H, HOOC-C<u>H_2_</u>-CH_2_-CH_2_), 2.43 (t, *J* = 7.3 Hz, 2H, HOOC-CH_2_-CH_2_-<u>CH_2_</u>), 4.03–4.12 (m, 2H, O-<u>CH_2_</u>-CH_3_), 4.59–4.65 (m, 1H, H-3), 5.37 (d, *J* = 5.0 Hz, 1H, H-6). ^13^C NMR (101 MHz, CDCl_3_): δ 178.46 (COOH), 172.43 (ester CO), 157.31 (C=N-O), 139.77 (C-5), 122.67 (C-6), 74.19 (C-3), 68.82 (O-<u>CH_2_</u>-CH_3_), 56.87, 56.30, 50.22, 43.79, 38.76, 38.24, 37.13, 36.78, 33.67, 33.03, 32.16, 31.94, 31.06, 27.91, 24.43, 23.30, 21.14, 20.06, 19.48, 15.93, 14.92, 13.33 (O-CH_2_-<u>CH_3_</u>). IR spectrum (CHCl_3_): 3517, 3098 (COOH), 2970, 2945 (CH_3_), 2676 (OH), 1723, 1714 (C=O), 1666 (C=C), 1626 (C=N). MS (ESI): *m/z* 496.3 (20%, M+Na), 474.3 (100%, M+H). HR-MS (ESI) *m/z*: For C_28_H_44_O_5_N [M-H] calcd, 472.3068; found, 472.3060.

#### 20-Oxo-pregn-5-en-3β-yl hemiglutarate O-(3-morpholinopropyl)oxime (**13**)

Compound **13** was prepared according to the *General procedure*. Starting from compound **25** (200 mg, 0.46 mmol), compound **13** (162 mg, 62%) was obtained as an oily compound by column chromatography (10-20% acetone in dichloromethane) with LC-MS purity of was 99%. [α]_D_ –19.6 (*c* 0.15, CHCl_3_). ^1^H NMR (400 MHz, CDCl_3_): δ 0.62 (s, 3H, H-18), 1.02 (s, 3H, H-19), 1.80 (s, 3H, H-21), 1.91–1.96 (m, 2H, HOOC-CH_2_-<u>CH_2_</u>-CH_2_), 2.30–2.34 (m, 2H, HOOC-<u>CH_2_</u>-CH_2_-CH_2_), 2.36–2.38 (m, 2H, HOOC-CH_2_-CH_2_-<u>CH_2_</u>), 2.64–2.69 (m, 4H, N-(<u>CH_2_</u>-CH_2_)_2_-O), 2.75–2.79 (m, 2H, O-CH_2_-<u>CH_2_</u>-N), 3.73–3.79 (m, 4H, N-(CH_2_-<u>CH_2_</u>)_2_-O), 4.23 (t, *J* = 5.6 Hz, 2H, O-<u>CH_2_</u>-CH_2_-N), 4.58–4.66 (m, 1H, H-3), 5.37 (d, *J* = 5.9 Hz, 1H, H-6). ^13^C NMR (101 MHz, CDCl_3_): δ 178.31 (COOH), 172.61 (ester CO), 158.01 (C=N-O), 139.80 (C-5), 122.60 (C-6), 74.04 (C-3), 70.35 (O-<u>CH_2_</u>-CH_2_-N), 66.39 (2 x C, N-(CH_2_-<u>CH_2_</u>)_2_-O), 57.28 (2 x C, N-(<u>CH_2_</u>-CH_2_)_2_-O), 56.86, 56.31, 53.59, 50.20, 43.81, 38.84, 38.26, 37.16, 36.78, 33.90, 32.18, 31.93, 27.93, 24.39, 23.23, 21.96, 21.17, 20.44, 19.49, 16.21, 13.40. IR spectrum (CHCl_3_): 3513, 3115, (COOH), 1722 (C=O), 1660 (C=C), 1467, 1455, 1397, 1190, 1138, 967 (morpholine). MS (negative ESI): *m/z* 558.5 (36%, M), 557.5 (100%, M–H). HR-MS (negative ESI) *m/z*: For C_32_H_49_O_6_N_2_ [M–H] calcd, 557.3596; found, 557.3594.

#### 17-Oxo-androst-5-en-3β-yl hemiglutarate O-methyloxime (**14**)

Compound **14** was prepared according to the *General procedure*. Starting from compound **26** (150 mg, 0.37 mmol), compound **14** (132 mg, 85%) was obtained as by column chromatography (5-12.5% methanol in dichloromethane). Recrystallization from acetone/heptane afforded LC-MS purity of 99%. Mp 210–212 °C. [α]_D_ –49.0 (*c* 0.2, CHCl_3_). ^1^H NMR (400 MHz, CDCl_3_): δ 0.93 (s, 3H, H-18), 1.04 (s, 3H, H-19), 1.91–2.01 (m, 2H, COOH-CH_2_-<u>CH_2_</u>-CH_2_), 2.35–2.40 (m, 2H, HOOC-<u>CH_2_</u>-CH_2_-CH_2_), 2.39–2.44 (m, 2H, COOH-CH_2_-CH_2_-<u>CH_2_</u>), 4.55–4.68 (m, 1H, H-3), 5.39 (d, *J* = 4.0 Hz, 1H, H-5). ^13^C NMR (101 MHz, CDCl_3_): δ 177.90 (COOH), 172.48 (ester CO), 171.59 (C=N-OH), 139.96 (C-5), 122.15 (C-6), 73.93 (C-3), 54.14, 50.23, 44.17, 38.20, 37.06, 36.84, 33.85, 33.75, 33.12, 31.42, 31.35, 27.89, 25.65, 23.37, 20.63, 20.28, 19.48, 17.03. IR spectrum (CHCl_3_): 3586, 3296 (oxime OH), 3513, 3122 (COOH), 1723 (C=O), 1682 (C=C). MS (negative ESI): *m/z* 417.3 (29%, M), 416.3 (100%, M–H). HR-MS (negative ESI) *m/z*: For C_24_H_34_O_5_N [M–H] calcd, 416.2442; found, 416.2441.

#### 17-Oxo-androst-5-en-3β-yl hemiglutarate O-ethyloxime (**15**)

Compound **15** was prepared according to the *General procedure*. Starting from compound **26** (150 mg, 0.37 mmol), compound **15** (152 mg, 92%) was obtained as by column chromatography (CH_2_Cl_2_/acetone/acetic acid (10:1:0 to 1:1:0.1)). Recrystallization from acetone/heptane afforded LC-MS purity of 99%. Mp 73–75 °C. [α]_D_ –37.7 (*c* 0.2, CHCl_3_). ^1^H NMR (400 MHz, CDCl_3_): δ 0.91 (s, 3H, H-18), 1.03 (s, 3H, H-19), 1.23 (t, *J* = 7.0 Hz, 3H, O-CH_2_-C<u>H_3_</u>), 1.74–1.87 (m, 2H, HOOC-CH_2_-<u>CH_2_</u>-CH_2_), 1.89–2.09 (m, 2H, HOOC-<u>CH_2_</u>-CH_2_-CH_2_), 2.38–2.52 (m, 2H, HOOC-CH_2_-CH_2_-<u>CH_2_</u>), 4.04–4.08 (m, 2H, O-<u>CH_2_</u>-CH_3_), 4.56–4.64 (m, 1H, H-3), 5.37 (d, *J =* 4.1 Hz, 1H, H-6). ^13^C NMR (101 MHz, CDCl_3_): δ 179.15 (COOH), 172.90 (ester CO), 170.15 (C=N-O), 140.02 (C-5), 122.25 (C-6), 73.99 (C-3), 68.91 (O-<u>CH_2_</u>-CH_3_), 54.26, 50.37, 43.81, 43.81, 38.23, 37.09, 36.86, 34.22, 32.03, 31.54, 31.42, 27.87, 25.90, 23.47, 20.71, 19.49, 17.18, 14.83, 14.25. IR spectrum (CHCl_3_): 3516, 3088 (COOH), 2970, 2950 (CH_3_), 2873, 2861 (CH_2_), 2650 (OH), 1722, 1713 (C=O), 1670 (C=C), 1655 (C=N-O), 1050, 1043 (O-Et). MS (negative ESI): *m/z* 444.3 (100%, M–H). HR-MS (negative ESI) *m/z*: For C_26_H_38_O_5_N [M–H] calcd, 444.2755; found, 444.2752.

#### 17-Oxo-androst-5-en-3β-yl hemiglutarate O-(3-morpholinopropyl)oxime (**16**)

Compound **16** was prepared according to the *General procedure*. Starting from **26** (200 mg, 0.49 mmol), compound **16** (202 mg, 77%) – a mixture of *E* and *Z*-isomers was obtained as an oily compound by column chromatography (5-12.5% methanol in dichloromethane) with LC-MS purity of was 99%. ^1^H NMR (400 MHz, CDCl_3_): δ 0.93 (s, 3H, H-18), 1.06 (s, 3H, H-19), 2.34–2.40 (m, 6H, HOOC-<u>CH_2_</u>-<u>CH_2_</u>-<u>CH_2_</u>), 2.64–2.75 (m, 4 H, N-(<u>CH_2_</u>-CH_2_)_2_-O), 2.82 (t, *J* = 5.6 Hz, 2H, O-CH_2_-<u>CH_2_</u>-N), 3.77–3.82 (m, 4H, N-(CH_2_-<u>CH_2_</u>)_2_-O), 4.13–4.30 (m, 2H, O-<u>CH_2_</u>-CH_2_-N), 4.58–4.68 (m, 1H, H-3), 5.40 (d, *J* = 5.0 Hz, 1H, H-6). ^13^C NMR (101 MHz, CDCl_3_): δ (mixture of isomers, m=minor), peak for COOH is not visible, 172.67 (ester CO), 170.99 (C=N-O-), 140.00 (C-5), 122.19 (C-6), 73.94 (C-3), 70.27 (O-<u>CH_2_</u>-CH_2_-N), 66.38 (m), 66.26 (2C, N-(CH_2_-<u>CH_2_</u>)_2_-O), 57.16 (2C, N-(<u>CH_2_</u>-CH_2_)_2_-O), 57.04 (m), 55.31 (m), 54.25, 53.56 (m), 53.51, 50.32, 50.05 (m), 46.06 (m), 43.97, 38.24, 37.09, 37.02 (m), 36.86, 36.81 (m), 34.16, 33.90, 33.73 (m), 31.50, 31.40, 30.99 (m), 29.84 (m), 29.40 (m), 29.27, 27.88, 26.17, 24.06 (m), 23.44, 22.83 (m), 20.88 (m), 20.68, 20.49, 19.49 (m), 19.41, 17.20, 14.26 (m), 14.01. IR spectrum (CHCl_3_): 3515, 3115 (COOH), 1723 (C=O), 1668 (C=C), 1467, 1455, 1189, 1143, 1027, 970, 610 (morpholine), 1234, 1116 (ester C-O). MS (negative ESI): *m/z* 530.4 (38%, M), 529.4 (100%, M–H). HR-MS (negative ESI) *m/z*: For C_30_H_45_O_6_N_2_ [M–H] calcd, 529.3283; found, 529.3282.

#### 20-Oxo-pregn-5-en-3β-yl hemiglutarate O-methyloxime (**17**)

Compound **17** was prepared according to *General procedure I - Synthesis of oxime ethers with O-alkyl hydroxylamine hydrochloride and sodium acetate*. Starting from compound **25** (130 mg, 0.30 mmol), compound **17** (101 mg, 73%) was obtained as an amorphous compound by column chromatography (dichloromethane/acetone/acetic acid, 4:1:0 to 0:1:0.1). Recrystallization from acetone/heptane afforded LC-MS purity of 99%. Mp 138–140 °C. [α]_D_ –26.8 (*c* 0.17, CHCl_3_). ^1^H NMR (400 MHz, CDCl_3_): δ 0.64 (s, 3H, H-18), 1.02 (s, 3H, H-19), 1.81 (s, 3H, H-21), 2.29–2.34 (m, 2H, HOOC-CH_2_-<u>CH_2_</u>-CH_2_), 2.37 (t, *J* = 7.3 Hz, 2H, HOOC-<u>CH_2_</u>-CH_2_-CH_2_), 2.43 (t, *J* = 5.0 Hz, 2H, HOOC-CH_2_-CH_2_-<u>CH_2_</u>), 3.83 (s, 3H, O-<u>CH_3_</u>), 4.58–4.65 (m, 1H, H-3), 5.38 (d, *J* = 5.0 Hz, 1H, H-6). ^13^C NMR (101 MHz, CDCl_3_): δ 178.24 (COOH), 172.46 (ester CO) 157.69 (C=N-O), 139.78 (C-5), 122.66 (C-6), 74.18 (C-3), 61.33 (O-<u>CH_3_</u>), 56.76, 56.30, 50.21, 43.78, 38.77, 38.24, 37.14, 36.78, 33.69, 33.06, 32.15, 31.93, 27.91, 24.41, 23.23, 21.14, 20.09, 19.48, 15.77, 13.30. IR spectrum (CHCl_3_): 3516, 3090 (COOH), 2967, 2943 (CH_3_), 2670 (OH), 1726, 1713 (C=O), 1670 (C=C), 1626 (C=N). MS (negative ESI): *m/z* 458.3 (100%, M–H). HR-MS (negative ESI) *m/z*: For C_27_H_40_O_5_N [M–H] calcd, 458.2912; found, 458.2909.

#### 20-Oxo-pregn-5-en-3β-yl hemiglutarate O-propyloxime (**18**)

Compound **18** was prepared according to the *General procedure*. Starting from compound **25** (130 mg, 0.30 mmol), compound **22** (81 mg, 55%) was obtained as an amorphous compound by column chromatography (5-10% methanol in dichloromethane). Recrystallization from acetone/heptane afforded LC-MS purity of 99%. Mp 70.5–72 °C. [α]_D_ –27.7 (*c* 0.2, CHCl_3_). ^1^H NMR (400 MHz, CDCl_3_): δ 0.63 (s, 3H, H-18), 0.92 (t, *J* = 7.4 Hz, 3H O-CH_2_-CH_2_-<u>CH_3_</u>), 1.02 (s, 3H, H-19), 1.82 (s, 3H, H-21), 1.91–1.97 (m, 2H, HOOC-CH_2_-<u>CH_2_</u>-CH_2_), 2.37 (t, *J* = 7.3 Hz, 2H, HOOC-<u>CH_2_</u>-CH_2_-CH_2_), 2.43 (t, *J* = 7.3 Hz, 2H, HOOC-CH_2_-CH_2_-<u>CH_2_</u>), 3.98 (td, *J* = 6.7, 2.1 Hz, 2H, O-<u>CH_2_</u>-CH_2_), 4.62 (dt, *J* = 11.6, 6.8 Hz, 1H, H-3), 5.38 (d, *J* = 5.1 Hz, 1H, H-5). ^13^C NMR (101 MHz, CDCl_3_): δ 178.10 (COOH), 172.45 (ester CO), 157.20 (C=N-O), 139.78 (C-5), 122.68 (C-6), 75.00 (O-<u>CH_2_</u>-CH_2_), 74.19 (C-3), 56.88, 56.30, 50.23, 43.80, 38.78, 38.25, 37.14, 36.79, 33.69, 33.03, 32.17, 31.95, 27.92, 24.43, 23.28, 22.69, 21.15, 20.10, 19.48, 15.89, 13.34, 10.56 (O-CH_2_-CH_2_-<u>CH_3_</u>). IR spectrum (CHCl_3_): 3516, 3088 (COOH), 2672 (OH), 1727, 1713 (C=O), 1671 (C=C), 1626 (C=N). MS (negative ESI): *m/z* 487.3 (28%, M), 486.3 (100%, M–H). HR-MS (negative ESI) *m/z*: For C_29_H_44_O_5_N [M–H] calcd, 486.3225; found, 486.3218.

#### 20-Oxo-pregn-5-en-3β-yl hemiglutarate O-isopropyloxime (**19**)

Compound **19** was prepared according to *General procedure I - Synthesis of oxime ethers with O-alkyl hydroxylamine hydrochloride and sodium acetate*. Starting from compound **25** (130 mg, 0.30 mmol), compound **22** (72 mg, 49%) was obtained as an amorphous compound by column chromatography (5-10% methanol in dichloromethane). Recrystallization from acetone/heptane afforded LC-MS purity of 98%. Mp 114–116 °C. [α]_D_ +12 (*c* 0.2, CHCl_3_). ^1^H NMR (400 MHz, CDCl_3_): δ 0.64 (s, 3H, H-18), 1.02 (s, 3H, H-19), 1.21 (dd, *J* = 6.2, 1.0 Hz, 6H, O-CH(<u>CH_3_)_2_</u>), 1.80 (s, 3H, H-21), 1.93–2.02 (m, 2H, HOOC-CH_2_-<u>CH_2_</u>-CH_2_), 2.37 (t, *J =* 7.3 Hz, 2H, HOOC-<u>CH_2_</u>-CH_2_-CH_2_), 2.43 (t, *J =* 7.3 Hz, 2H, HOOC-CH_2_-CH_2_-<u>CH_2_</u>), 4.28 (hept, *J* = 6.2 Hz, 1H, O-<u>CH</u>(CH_3_)_2_), 4.56–4.68 (m, 1H, H-3), 5.38 (d, *J* = 5.0 Hz, 1H, H-6). ^13^C NMR (101 MHz, CDCl_3_): δ 177.94 (COOH), 172.45 (ester CO), 156.58 (C=N-O), 139.78 (C-5), 122.70 (C-6), 74.42 (O-<u>CH</u>(CH_3_)_2_), 74.20 (C-3), 57.01, 56.32, 50.25, 43.78, 38.79, 38.25, 37.15, 36.79, 33.69, 32.99, 32.18, 31.96, 27.92, 24.43, 23.36, 21.97, 21.85, 21.16, 20.09, 19.48, 16.03, 13.35. IR spectrum (CHCl_3_): 3517, 3089 (COOH), 2672 (OH), 1754, 1725, 1713 (C=O), 1671 (C=C), 1629 (C=N). MS (negative ESI): *m/z* 486.3 (100%, M–H). HR-MS (negative ESI) *m/z*: For C_29_H_44_O_5_N [M–H] calcd, 486.3225; found, 486.3219.

#### 20-Oxo-pregn-5-en-3β-yl hemiglutarate O-(2-propenyl)oxime (**20**)

Compound **20** was prepared according to the *General procedure*. Starting from compound **25** (100 mg, 0.23 mmol), compound **20** (71 mg, 63%) was obtained as an amorphous compound by column chromatography (10-35% acetone in dichloromethane). Recrystallization from acetone/heptane afforded LC-MS purity of 99%. Mp 91–93 °C. [α]_D_ –21.8 (*c* 0.2, CHCl_3_). ^1^H NMR (400 MHz, CDCl_3_): δ 0.64 (s, 3H, H-18), 1.02 (s, 3H, H-19), 1.84 (s, 3H, H-21), 1.91–1.99 (m, 2H, HOOC-CH_2_-<u>CH_2_</u>-CH_2_), 2.37 (t, *J* = 7.3 Hz, 2H, HOOC-<u>CH_2_</u>-CH_2_-CH_2_), 2.43 (t, *J* = 7.3 Hz, 2H, HOOC-CH_2_-CH_2_-<u>CH_2_</u>), 4.54 (dt, *J* = 5.6, 1.5 Hz, 2H, N-O-<u>CH_2_</u>), 4.57–4.68 (m, 1H, H-3), 5.16 (dt, *J* = 10.5, 1.5 Hz, 1H, CH=<u>CH_2_</u>), 5.26 (dq, *J* = 17.3, 1.7 Hz, 1H, CH=<u>CH_2_</u>), 5.37 (dd, *J* = 4.5, 2.8 Hz, 1H, H-6), 5.99 (ddt, *J* = 17.3, 10.7, 5.5 Hz, 1H, <u>CH</u>=CH_2_). ^13^C NMR (101 MHz, CDCl_3_): δ 178.49 (COOH), 172.43 (ester CO), 157.88 (C=N-O), 139.77 (C-5), 135.06 (O-CH_2_-<u>CH</u>=CH_2_), 122.66 (C-6), 116.79 (O-CH_2_-CH=<u>CH_2_</u>), 74.35 (N-O-<u>CH_2_</u>-CH=CH_2_), 74.19 (C-3), 56.83, 56.29, 50.21, 43.80, 38.76, 38.24, 37.13, 36.78, 33.67, 33.04, 32.15, 31.93, 27.91, 24.41, 23.26, 21.14, 20.05, 19.47, 16.00, 13.35. IR spectrum (CHCl_3_): 3516, 3083 (COOH), 3011 (=CH), 1724, 1713 (C=O), 1671 (C=C), 1647 (vinyl C=C), 1624 (C=N). MS (ESI): *m/z* 486.3 (100%, M+H). HR-MS (ESI) *m/z*: For C_29_H_44_O_5_N [M+H] calcd, 486.3214; found, 486.3214.

#### 20-Oxo-pregn-5-en-3β-yl hemiglutarate O-(phenylmethyl)oxime (**21**)

Compound **21** was prepared according to the *General procedure*. Starting from compound **25** (100 mg, 0.23 mmol), compound **21** (111 mg, 90%) was obtained as an oily compound by column chromatography (6-25% acetone in dichloromethane) with the LC-MS purity of 99%. [α]_D_ –11.6 (*c* 0.3, CHCl_3_). ^1^H NMR (400 MHz, CDCl_3_): 0.58 (s, 3H, H-18), 1.01 (s, 3H, H-19), 1.85 (s, 3H, H-21), 1.93–1.98 (m, 2H, HOOC-CH_2_-<u>CH_2_</u>-CH_2_), 2.37 (t, *J* = 7.3 Hz, 2H, HOOC-<u>CH_2_</u>-CH_2_-CH_2_), 2.42 (t, *J* = 7.3 Hz, 2H, HOOC-CH_2_-CH_2_-<u>CH_2_</u>), 4.55–4.68 (m, 1H, H-3), 5.08 (s, 2H, N-O-CH_2_-), 5.37 (dd, *J =* 4.5, 2.8 Hz, 1H, H-6), 7.28–7.37 (m, 5H, phenyl). ^13^C NMR (101 MHz, CDCl_3_): δ 178.16 (COOH), 172.47 (ester CO), 158.11 (C=N-O), 139.76 (C-5), 138.83 (arom), 128.37 (arom), 128.30 (arom), 128.21 (arom), 128.09 (arom), 127.55 (arom), 122.63 (C-6), 75.44 (N-O-<u>CH_2_</u>), 74.15 (C-3), 56.86, 56.29, 50.19, 43.80, 38.75, 38.23, 37.11, 36.75, 33.69, 33.03, 32.14, 31.91, 27.89, 24.38, 23.21, 21.12, 20.09, 19.46, 16.19, 13.26. IR spectrum (CHCl_3_): 3513 (COOH), 3089, 3065, 3030, 1496, 1454, 1185, 1157, 917, 839, 699 (phenyl), 2672 (OH), 1728, 1713 (C=O), 1665 (C=C), 1630 (C=N). MS (ESI): *m/z* 536.3 (100%, M+H). HR-MS (ESI) *m/z*: For C_33_H_46_O_5_N [M+H] calcd, 536.3370; found, 536.3370.

#### 20-Oxo-pregn-5-en-3β-yl hemisuccinate O-ethyloxime (**22**)

Compound **22** was prepared according to *General procedure I - Synthesis of oxime ethers with O-alkyl hydroxylamine hydrochloride and sodium acetate*. Starting from compound **27** (208 mg, 0.50 mmol), compound **22** (164 mg, 71%) was obtained as an amorphous compound by column chromatography (10-50% acetone in dichloromethane). Recrystallization from acetone/heptane afforded LC-MS purity of 99%. Mp 151–152 °C. [α]_D_ –20.2 (*c* 0.15, CHCl_3_). ^1^H NMR (400 MHz, CDCl_3_): δ 0.64 (s, 3H, H-18), 1.02 (s, 3H, H-19), 1.23 (t, 3H, *J* = 7.0 Hz, O-CH_2_-<u>CH_3_</u>), 1.82 (s, 3H, H-21), 2.60 (t, *J* = 6.1 Hz, 2H, HOOC-<u>CH_2_</u>-CH_2_), 2.65 (t, *J* = 6.1 Hz, 2H, HOOC-CH_2_-<u>CH_2_</u>), 4.04–4.11 (m, 2H, O-<u>CH_2_</u>-CH_3_), 4.58–4.68 (m, 1H, H-3), 5.37 (d, *J* = 5.0 Hz, 1H, H-5). ^13^C NMR (101 MHz, CDCl_3_): δ 177.48 (COOH), 171.82 (ester CO), 157.25 (C=N-O), 139.73 (C-5), 122.71 (C-6), 74.61 (C-3), 68.82 (O-<u>CH_2_</u>-CH_3_), 56.89, 56.31, 50.22, 43.79, 38.77, 38.15, 37.12, 36.78, 32.16, 31.95, 29.48, 29.22, 27.83, 24.43, 23.30, 21.15, 19.48, 15.94, 14.92, 13.33 (O-CH_2_-<u>CH_3_</u>). IR spectrum (CHCl_3_): 3517, 3100 (COOH), 2672, 2577 (OH), 1728 (ester C=O), 1717 (C=O), 1671 (C=C), 1628 (C=N). MS (negative ESI): *m/z* 459.3 (26%, M), 458.3 (100%, M–H). HR-MS (negative ESI) *m/z*: For C_27_H_40_O_5_N [M–H] calcd, 458.2912; found, 458.2911.

#### 20-Oxo-pregn-5-en-3β-yl hemiadipate O-ethyloxime (**23**)

Compound **23** was prepared according to *General procedure I - Synthesis of oxime ethers with O-alkyl hydroxylamine hydrochloride and sodium acetate*. Starting from compound **28** (148 mg, 0.33 mmol), compound **23** (94 mg, 58%) was obtained as an amorphous compound by column chromatography (15-30% acetone in dichloromethane) with LC-MS purity of 99%. Mp 113– 115 °C. [α]_D_ –21.4 (*c* 0.17, CHCl_3_). ^1^H NMR (400 MHz, CDCl_3_): δ 0.64 (s, 3H, H-18), 1.02 (s, 3H, H-19), 1.23 (t, *J* = 7.0 Hz, 3H, O-CH_2_-<u>CH_3_</u>), 1.81 (s, 3H, H-21), 1.84–1.87 (m, 2H, HOOC-CH_2_-CH_2_-<u>CH_2_</u>-CH_2_), 2.28–2.38 (m, 6H, HOOC-<u>CH_2_-CH_2_-</u>CH_2_<u>-CH_2_</u>), 4.07 (q, *J =* 7.0 Hz, 2H, O-<u>CH_2_</u>-CH_3_), 4.62 (td, *J =* 8.6, 7.8, 3.3 Hz, 1H, H-3), 5.37 (d, *J* = 4.9 Hz, 1H, H-5). ^13^C NMR (101 MHz, CDCl_3_): δ 178.58 (COOH), 172.87 (ester CO), 157.25 (C=N-O), 139.82 (C-5), 122.62 (C-6), 74.02 (C-3), 68.81 (O-<u>CH_2_</u>-CH_3_), 56.89, 56.30, 50.23, 43.78, 38.77, 38.25, 37.15, 36.79, 34.38, 33.68, 32.16, 31.94, 27.92, 24.53, 24.42, 24.24, 23.29, 21.14, 19.48, 15.93, 14.92, 13.33 (O-CH_2_-<u>CH_3_</u>). IR spectrum (CHCl_3_): 3517, 3091 (COOH), 2670, 2565 (OH), 1736 (acid C=O), 1725 (ester C=O), 1713 (C=O), 1670 (C=C). MS (negative ESI): *m/z* 487.3 (14%, M), 486.3 (100%, M–H). HR-MS (negative ESI) *m/z*: For C_29_H_44_O_5_N [M–H] calcd, 486.3225; found, 486.3227.

#### 20-(Ethyloxyimino)-pregn-5-en-3β-ol (**12**)

Compound **24** was prepared according to the *General procedure*. Starting from pregnenolone (100 mg, 0.32 mmol), compound **24** (112 mg, 98%) was obtained as an amorphous solid compound by column chromatography (10-20% acetone in dichloromethane). Recrystallization from acetone/heptane afforded LC-MS purity of was 99%. Mp 111–113 °C. [α]_D_ –28.7 (*c* 0.3, CHCl_3_). ^1^H NMR (400 MHz, CDCl_3_): δ 0.64 (s, 3H, H-18), 1.01 (s, 3H, H-19), 1.23 (t, 3H, *J* = 7.0 Hz, O-CH_2_-<u>CH_3_</u>), 1.82 (s, 3H, H-21), 3.52 (tdd, *J* = 11.0, 5.2, 4.1 Hz, 1H, H-3), 4.07 (q, *J* = 7.0 Hz, 2H, O-<u>CH_2_</u>-CH_3_), 5.35 (dt, *J* = 5.3, 2.0 Hz, 1H, H-6). ^13^C NMR (101 MHz, CDCl_3_): δ 157.29 (C=N-O), 140.93 (C-5), 121.68 (C-6), 71.89 (C-3), 68.82 (O-<u>CH_2_</u>-CH_3_), 56.89, 56.37, 50.33, 43.80, 42.42, 38.81, 37.41, 36.69, 32.20, 31.95, 31.78, 24.44, 23.29, 21.19, 19.57, 15.91, 14.92, 13.34 (O-CH_2_-<u>CH_3_</u>). IR spectrum (CHCl_3_): 3607 (OH), 3009 (=CH), 1667 (C=C), 1620 (C=N). MS (ESI): *m/z* 360.3 (100%, M+H). HR-MS (ESI) *m/z*: For C_23_H_38_O_2_N [M+H] calcd, 360.2897; found, 360.2897.

Alternative synthesis: A stirred solution of pregnenolone (500 mg, 1.5 mmol) in dry pyridine (5 mL) and dry triethyl amine (5 mL) was treated with *O*-ethylhydroxylamine hydrochloride (318 mg, 3.15 mmol). The progress of the reaction was repeatedly checked by TLC. After 70 h, the reaction mixture was quenched with water with crushed ice, a white precipitate was collected, washed with water, and dried. The crude material was purified by column chromatography (10-25% acetone in dichloromethane) affording compound **24** (568 mg, 92%).

#### 20-Oxo-pregn-5-en-3β-yl hemiglutarate (**25**)

Compound **25** was prepared according to the literature (Krausova, Slavikova et al. 2018) with LC-MS purity of 99%. Mp 139–141 °C, lit.(Krausova, Slavikova et al. 2018) 137–139°C. [α]_D_ +9.3 (*c* 0.150, CHCl_3_), lit. (Krausova, Slavikova et al. 2018) +9.1 (*c* 0.3, CHCl_3_). ^1^H NMR (400 MHz, CDCl_3_): δ 0.62 (s, 3H, H-18), 1.00 (s, 3H, H-19), 1.90–1.98 (m, 2H, HOOC-CH_2_-<u>CH_2_</u>-CH_2_), 2.11 (s, 3H, H-21), 2.36 (t, *J* = 7.3 Hz, 2H, HOOC-<u>CH_2_</u>-CH_2_-CH_2_), 2.42 (t, *J* = 7.3 Hz, 2H, HOOC-CH_2_-CH_2_-<u>CH_2_</u>), 2.52 (1H, t, *J* = 8.9, H-17), 4.53–4.66 (m, 1H, H-3), 5.36 (d, *J* = 5.1, 1H, H-6). ^13^C NMR (101 MHz, CDCl_3_): δ 209.88 (C=O), 178.72 (COOH), 172.43 (ester CO), 139.71 (C-5), 122.49 (C-6), 74.09 (C-3), 63.80, 56.95, 50.00, 44.12, 38.90, 38.18, 37.10, 36.72, 33.65, 33.09, 31.93, 31.88, 31.65, 27.86, 24.60, 22.96, 21.15, 20.03, 19.42, 13.33.

#### 17-Oxo-androst-5-en-3β-yl hemiglutarate (**26**)

Compound **26** was prepared according to the literature (Krausova, Slavikova et al. 2018) with LC-MS purity of 99%. Mp 120–121 °C, lit.(Krausova, Slavikova et al. 2018) 126–128°C. [α]_D_ +0.0 (*c* 0.164, CHCl_3_), lit. (Krausova, Slavikova et al. 2018) +0.0 (*c* 0.2, CHCl_3_). ^1^H NMR (400 MHz, CDCl_3_): δ 0.88 (s, 3H, H-18), 1.04 (s, 3H, H-19), 1.92–1.98 (m, 2H, HOOC-CH_2_-<u>CH_2_</u>-CH_2_), 2.37 (t, *J* = 7.3 Hz, 2H, HOOC-<u>CH_2_</u>-CH_2_-CH_2_), 2.41–2.45 (m, 2H, HOOC-CH_2_- CH_2_-<u>CH_2_</u>), 4.58–4.67 (m, 1H, H-3), 5.40 (1H, d, *J* = 4.0, H-6). ^13^C NMR (101 MHz, CDCl_3_): δ ketone at 221 is not visible (the scale of the measurement was below 219.5 PPM), 178.56 (COOH), 172.40 (ester CO), 140.00 (C-5), 122.05 (C-6), 73.99 (C-3), 51.84, 50.27, 47.68, 38.21, 37.05, 36.87, 35.98, 33.65, 33.06, 31.60, 31.54, 30.91, 27.85, 22.02, 20.46, 20.05, 19.48, 13.68.

#### 20-Oxo-pregn-5-en-3β-yl hemisuccinate (**27**)

Compound **27** was prepared according to the literature(Krausova, Slavikova et al. 2018) with LC-MS purity of 99%. Mp 164–165 °C, lit.(Krausova, Slavikova et al. 2018) 161–163°C. [α]_D_ +11.6 (*c* 0.2, CHCl_3_), lit.(Krausova, Slavikova et al. 2018) +14.3 (*c* 0.2, CHCl_3_). ^1^H NMR (400 MHz, CDCl_3_): δ 0.63 (s, 3H, H-18), 1.02 (s, 3H, H-19), 2.12 (s, 3H, H-21), 2.53 (t, *J* = 8.9, 1H, H-17), 2.60 (ddd, *J* = 7.7, 5.8, 1.5 Hz, 2H, HOOC-CH_2_-<u>CH_2_</u>), 2.66–2.70 (m, 2H, HOOC-CH_2_- CH_2_-<u>CH_2_</u>), 4.59–4.68 (m, 1H, H-3), 5.37–5.39 (m, 1H, H-6). ^13^C NMR (101 MHz, CDCl_3_): δ 209.81 (C=O), 177.35 (COOH), 171.70 (ester CO), 139.70 (C-5), 122.58 (C-6), 74.55 (C-3), 63.83, 56.99, 50.02, 44.14, 38.93, 38.11, 37.11, 36.74, 31.96, 31.91, 31.69, 29.38, 29.02, 27.80, 24.63, 22.98, 21.18, 19.44, 13.37.

#### 20-Oxo-pregn-5-en-3β-yl hemiadipate (**28**)

Compound **28** was prepared according to the literature (Krausova, Slavikova et al. 2018) with LC-MS purity of 99%. Mp 135–138 °C, lit.(Krausova, Slavikova et al. 2018) 135–136°C. [α]_D_ +10.5 (*c* 0.2, CHCl_3_), lit. (Krausova, Slavikova et al. 2018) +3.0 (*c* 0.2, CHCl_3_). ^1^H NMR (400 MHz, CDCl_3_): δ 0.60 (s, 3H, H-18), 0.99 (s, 3H, H-19), 1.80–1.87 (m, 2H, H-adipate), 2.09 (s, 3H, H-21), 2.25–2.32 (m, 4H, H-adipate), 2.51 (1H, t, *J* = 8.9, H-17), 4.53–4.62 (m, 1H, H-3), 5.33–5.36 (m, 1H, H-6). ^13^C NMR (101 MHz, CDCl_3_): δ 209.67 (C=O), 176.04 (COOH), 172.90 (ester CO), 139.77 (C-5), 122.36 (C-6), 73.78 (C-3), 63.75, 56.90, 49.95, 44.06, 38.85, 38.15, 37.07, 36.67, 34.39, 33.76, 31.89, 31.83, 31.62, 27.82, 24.57, 24.55, 24.40, 22.89, 21.10, 19.38, 13.29.

## Biological assays methods

### Transfection and maintenance of the HEK293 cells

HEK293 cells (American Type Culture Collection, ATCC No. CRL-1573, Rockville, MD, USA) were cultured in Opti-MEM I (Invitrogen, Carlsbad, CA, USA) supplemented with 5% fetal bovine serum (FBS; PAN Biotech, Aidenbach, Germany) at 37°C in 5% CO₂. 24 h before transfection, HEK293 cells were plated in 24-well plates at a density of 2 × 10⁵ cells per well. The next day, the cells were transfected with cDNA encoding rat GluN1-1a (GluN1; GenBank accession number)(Hollmann, Boulter et al. 1993) and GluN2B (GenBank accession number)(Monyer, Sprengel et al. 1992) subunits (in the pCI-neo expression vector), along with green fluorescent protein (GFP; in the pQBI 25 vector, Takara, Tokyo, Japan). Briefly, equal amounts (200 ng) of cDNAs encoding GluN1-1a, GluN2B, and GFP were mixed with 0.6 μl of Matra-A reagent (IBA, Gottingen, Germany) in 50 μL of Opti-MEM I and added to confluent HEK293 cells cultured in 24-well plates. After trypsinization, the cells were resuspended in Opti-MEM I containing 1% FBS, supplemented with 20 mM MgCl₂, 1 mM D,L-2-amino-5-phosphonovaleric acid, 3 mM kynurenic acid, and 1 μM ketamine, and plated on 30 mm glass coverslips coated with collagen and poly-L-lysine. Transfected cells were identified by GFP epifluorescence. Electrophysiology experiments were performed 24–48 h after transfection.

Electrophysiology at HEK293 cells. Whole-cell current recordings, voltage-clamped at a holding potential of −60 mV, were performed at room temperature using a patch-clamp amplifier (Axopatch 200B; Molecular Devices, Sunnyvale, CA, USA) after capacitance and series resistance (<10 MΩ) compensation (80–90%). NMDAR current responses were low-pass filtered at 2 kHz, digitally sampled at 5 kHz, and analysed using pClamp software (version 10.6; Molecular Devices). Patch pipettes (3–5 MΩ), pulled from borosilicate glass, were filled with an intracellular solution (ICS) containing (in mM): 15 CsCl, 10 BAPTA, 3 MgCl₂, 1 CaCl₂, 120 gluconic acid, 10 HEPES, and 2 ATP-Mg salt (pH adjusted to 7.2 with CsOH). The extracellular solution (ECS) contained (in mM): 160 NaCl, 2.5 KCl, 0.2 EDTA, 10 glucose, 10 HEPES, and 0.7 CaCl₂ (pH adjusted to 7.3 with NaOH). NMDAR responses were induced by 1 μM glutamate and 30 μM glycine. 1% DMSO was present in all control and testing solutions. Solution applications were performed using a microprocessor-controlled multibarrel fast-perfusion system with a solution exchange rate around the cells of ∼10 ms (Vyklicky, Korinek et al. 2016).

## *In Vitro* ADME Profiling

### Microsomal stability

Microsomal stability assay was performed using the 1 mg/mL rat pooled liver microsomal preparation (Thermo Scientific) and 10 µM compounds in 90 mM TRIS-Cl buffer pH 7.4 containing 2 mM NADPH and 2 mM MgCl2 for 0, 30, and 60 min at 37°C. Reference compound Verapamil was also tested to provide positive control data for this assay. The reactions were terminated by the addition of four volumes of ice-cold methanol, mixed vigorously and left at −20°C for 1h. After that, the samples were centrifuged and the supernatants were analysed by means of ECHO-MS® System (Sciex, Framingham, MA, USA). Zero time points were prepared by adding ice-cold methanol to the mixture of the compound with cofactors prior to the addition of microsomes. The microsomal half-lives (t1/2) were calculated using the equation t1/2=0.693/*k*, where *k* is the slope found in the linear fit of the natural logarithm for the fraction remaining of the parent compound vs. incubation time. Intrinsic clearance (CL_int_) was calculated using the following formula:

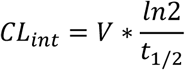

where *V* is the incubation volume per milligram of microsomal protein (µL/mg) and t_1/2_ is the microsomal half-life.

### PAMPA permeability

Permeation experiments were conducted in polystyrene 96-well filter plates with PVDF membrane pre-coated with structured layers of phospholipids (Corning Gentest^TM^) according to the manufacturer’s instructions. Briefly, the plate was warmed to room temperature and meanwhile, the solutions of the compounds were diluted from 10 mM DMSO stocks to TRIS-HCl buffer pH 7.4 to reach 100 μM final concentration. The DMSO concentration in the samples was 1%. Then, 300 μL of these solutions were transferred into the donor (receiver) plate wells and 200 μL of the (TRIS) buffer were added in the acceptor (filter) plate wells. The filter plate and receiver plate were assembled and incubated for 5 h on an orbital shaker (300 rpm). After this, concentration of the tested substances in both the donor and acceptor compartments were determined with use of LC/MS. Liquid chromatography was performed using ExionLC AD pump and autosampler from Sciex (Framingham, MA, USA). The mobile phase was (A) 0.1% formic acid in water and (B) 0.1% formic acid in acetonitrile with the following gradient: 0 – 1 min 2 % (B); 1 – 5 min 2 – 98 % (B); 5 – 6 min 98 % (B); 6 – 8 min 2 % (B). The mobile phase flow rate was 300 μL/min. The column synergi fusion-RP from phenomemex (Torrance, CA, USA) was used and was kept at 40°C during the analysis. Actual compound concentration in the samples was quantified by means of multiple reaction monitoring (MRM) mass spectrometry using the Sciex QTRAP® 6500+. Permeability coefficients P_e_ (cm/s) and mass retention R (%) were calculated using the following formulas:

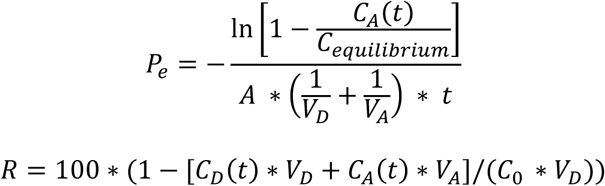

where *C_0_* is the initial compound concentration in donor well (mM), *C_D_(t)* is the compound concentration in donor well at time *t* (mM), *C_A_(t)* is the compound concentration in acceptor well at time *t* (mM), *V_D_* is the donor well volume (0.3 mL), *V_A_* is the acceptor well volume (0.2 mL), C_equilibrium_ =, *A* is the filter area (0.3 cm^2^) and *t* is the incubation time (18000 s = 5h).

### Stability in Rat Hepatocytes

Primary rat hepatocytes were isolated from 12-week old male Wistar rats (n=3) by two-step collagenase liver perfusion. Tribromoethanol was used as anesthetic agent at a dose of 250 mg/kg. Briefly, rat liver was first perfused for 3 min with a pre-perfusing solution (HBSS w/o Ca^2+^ and Mg^2+^, 20 mM HEPES pH 7.4, 0,5 mM EDTA), then for 3 min with washing solution (HBSS, 20 mM HEPES pH 7.4), and then for 15 min with perfusing solution (HBSS, 20 mM HEPES pH 7.4, 5 mM CaCl_2_, 5 mM MgCl_2_, 0.016 mg/mL collagenase Type II). Flow rate was maintained at 7 mL/min and all solutions were kept at 37°C. After *in situ* perfusion, the liver was excised, the liver capsule was opened and cells were suspended in William’s Medium E and filtered through a 70 µm membrane. Dead cells were removed by Percoll centrifugation (Percoll density: 1.06 g/mL, 50xg, 10 min, 20°C) and additional centrifugation in William’s Medium E (50xg, 5 min). Cells were subsequently diluted in William’s Medium E and their viability and cell suspension density were determined by Trypan Blue exclusion using a hemocytometer. Then, 10 mM DMSO stock solution of each test compound was diluted to 6 µM (2x concentration; final DMSO concentration – 0.25%) using William’s Medium E to create the working samples. Aliquots (50 µL) of the hepatocyte suspension were added to each test well of a 96-well plate immediately followed by the addition of 50 µL aliquot of the test compound or control solutions (final hepatocytes density – 0.5×10^6^/mL). The samples for each time point (0, 5, 10, 20, 40, 60, and 120 min) were prepared in duplicates for all the test and reference compounds. Incubations were done at 37 °C, 5% CO_2_, and 95% relative humidity in CO_2_ incubator. At appropriate time-points, 40 µL aliquots were removed from the wells and placed in 1.1 mL microtubes containing 200 µL of methanol and used for LC-MS/MS analysis.

Elimination constant (*k_el_*), half-life (*t_1/2_*) and intrinsic clearance (*Cl_int_*) were determined in plots of ln (percent remaining of parent compound) versus time, using linear regression analysis:

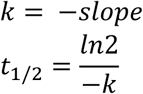

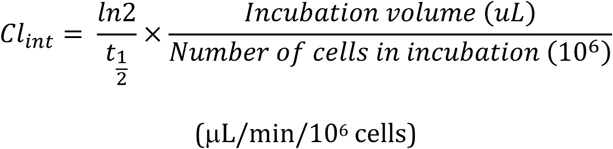

### Stability in rat plasma

Compounds were prepared as 1.6 mg/mL stock solutions in methanol. Then, 20 µL of stock solution was added to 1000 µL of human plasma, maintained at 37°C. Aliquots (50 µL) withdrawn at 0, 8, and 24 h were analyzed by HPLC. An aliquot of plasma was extracted with methanol (450 µL) containing an internal standard and the solution was vortexed (20 s) and centrifuged at 13500 rpm for 10 min. The supernatant was transferred to an autosampler vial and 10 µL was injected into an LC-MS system. The samples were analyzed using an Agilent 6230 TOF LC/MS. Samples were separated on a Waters ACQUITY UPLC CSH Phenyl-Hexyl column (100×2.1, 130Å, 1.7 µm) at a flow rate 0.3 mL/min. The concentration of mobile phase B (0.1% formic acid in acetonitrile) was gradually increased from 10 to 100 % in mobile phase A (0.1% formic acid in water) over 7 min. The mass spectrometry instrument was operated in a negative ion mode with a voltage of +3.00 kV applied to the capillary. The temperature, the flow rate of the nitrogen drying gas, the pressure of the nitrogen nebulizing gas, and the flow rate of the sheath gas were set at 325°C, 10 L/min, 40 psi, 390°C, and 11 L/min, respectively. Results are represented as a percentage of a compound remaining in spiked plasma.

### Thermodynamic solubility

The reference samples were prepared as follows: a weight corresponding to the 2.5 μmol was dissolved in 250 μL of DMSO resulting to the concentration of 10 mM stock solution. Each calibration level standard was prepared by diluting the stock solution in a mixture of PBS and acetonitrile (50:50 *v/v*). The samples for testing were prepared as follows: the volume of 1000 μL of PBS was added to the testing tube containing a weight of sample corresponding to the 0.4 mM. The mixture was shaken at 1000 RPM at 20°C for 24 h. The mixture was then filtered using a syringe polypropylene 0.45 μm filter. Samples and calibration standards are prepared using an automated system of robotic arm (PAL-RTC, Switzerland). Then, the sample was analysed on chromatographic system (Vanquish UHPLC, Thermo Fisher Scientific, Germany) connected to the diode array detector and consequently to the charged aerosol detector (both Vanquish, Thermo Fisher Scientific, Germany), Thermo Scientific Accucore C8 (100×3 mm; 2.6 μm) column, using gradient elution with acetonitrile/methanol/5mM formate buffer (pH = 3.00); flow: 1.0 cm^3^ min^-1^; column temperature: 40°C; CAD temperature: 30°C; sample volume: 5 mm^3^.

## Conclusions

A series of novel C-17 and C-20 steroidal oximes and oxime ethers were designed, synthesized, and evaluated as PAMs of GluN1/GluN2B NMDA receptors using patch-clamp technique. SAR analysis demonstrated that oxime ether modification at the C-20 position significantly influenced both efficacy and potency, with several compounds being more potent or efficacious than the endogenous reference compound, pregnenolone sulfate (**PES**). Notably, compounds **12** and **17** exhibited the highest efficacy (E_max_ = 673% and 503%, respectively) and potent activity in the low micromolar range (EC₅₀ = 8.7 µM and 6.1 µM, respectively).

Pregnenolone derivatives within this study consistently provided superior PAM activity compared to their DHEA analogues. Certain derivatives (e.g., compounds **18** and **19**) displayed concentration-dependent biphasic modulation, suggesting complex interaction with the NMDAR. Additionally, extension of the C-3 hemiester moiety (compounds **22** and **23**) did not yield further improvement in PAM mode over compound **12**. *In vitro* ADME profiling confirmed that the oxime ether scaffold affords favourable drug-like properties, including excellent metabolic and plasma stability, and enhanced solubility. Compound **12** was identified as the lead candidate based on its balanced profile of pharmacological activity and physicochemical properties.

These findings establish the oxime ether functionality as a viable bioisosteric strategy for steroidal NMDAR modulators and support further optimization of compound **12** as a lead for central nervous system-targeted therapeutic development.

## Author contributions

SKA – Data curation, Formal analysis, Investigation, Methodology; BHK – Conceptualization, Data curation, Formal analysis, Investigation, Methodology, Writing – original draft, Writing – review & editing; BK – Data curation, Formal analysis, Investigation, Methodology; KK – Data curation, Formal analysis, Investigation, Methodology; RS – Data curation, Formal analysis, Investigation, Methodology; MS – Data curation, Formal analysis, Investigation, Methodology; MB – Data curation, Formal analysis, Investigation, Methodology; HMK – Conceptualization, Data curation, Formal analysis, Funding acquisition, Investigation, Methodology, Project administration, Resources, Supervision, Validation Visualization, Writing – original draft, Writing – review & editing; LV – Conceptualization, Funding acquisition, Project administration, Resources, Supervision, Writing – original draft, Writing – review & editing; EK – Conceptualization, Funding acquisition, Project administration, Resources, Supervision, Writing – original draft, Writing – review & editing.

## Data availability

The data supporting this article have been included as part of the ESI.^†^

## Acknowledgements

This work was supported by the Czech Science Foundation (GACR): 23-04922S, the Academy of Sciences of the Czech Republic (RVO 61388963 and RVO 67985823), and by the Martina Roeselová Memorial Fellowship granted by the IOCB Tech Foundation. This work was also carried out as part of an ongoing research effort aimed at sustaining and developing the scientific the long-term objectives of the research program “PharmaBrain”, No. CZ.02.1.01/0.0/0.0/16_025/0007444, funded by the European Regional Development Fund – ERDF/ESF.

