## Supporting Info for "Positive Modulators of *N*-Methyl-*D*-Aspartate Receptor: Structure-Activity Relationship Study on Steroidal C-17 and C-20 Oxime Ethers"

### *Supporting information*

[a]Institute of Organic Chemistry and Biochemistry of the Czech Academy of Sciences, Flemingovo nam. 2, Prague 6 – Dejvice, 16610, Czech Republic
[b]Institute of Physiology of the Czech Academy of Sciences, Videnska 1083, Prague 4, 14220, Czech Republic
[c]University of Chemistry and Technology, Technická 5, Prague 6 – Dejvice, 166 28 Czech Republic

#### Table of Contents

**$^1$H and $^{13}$C NMR spectra of compounds 11-24 ...... 2**
**HR-MS and LC-MS spectra of compounds 11-28...... 11**

### $^{1}$H and $^{13}$C NMR spectra of compounds **11-24**

**Figure S1**. $^{1}$H NMR and $^{13}$C NMR and spectra of compound **11 (SKA-299)**

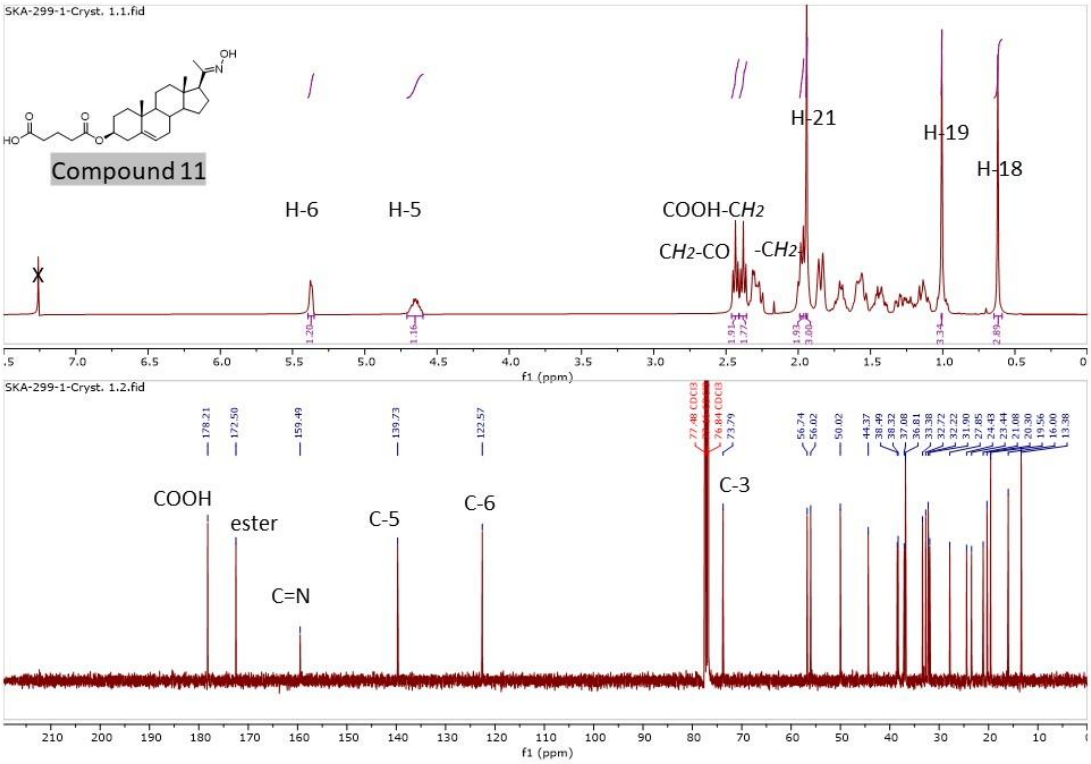


**Figure S2**. $^{1}$H NMR and $^{13}$C NMR and spectra of compound **12 (SKA-368)**

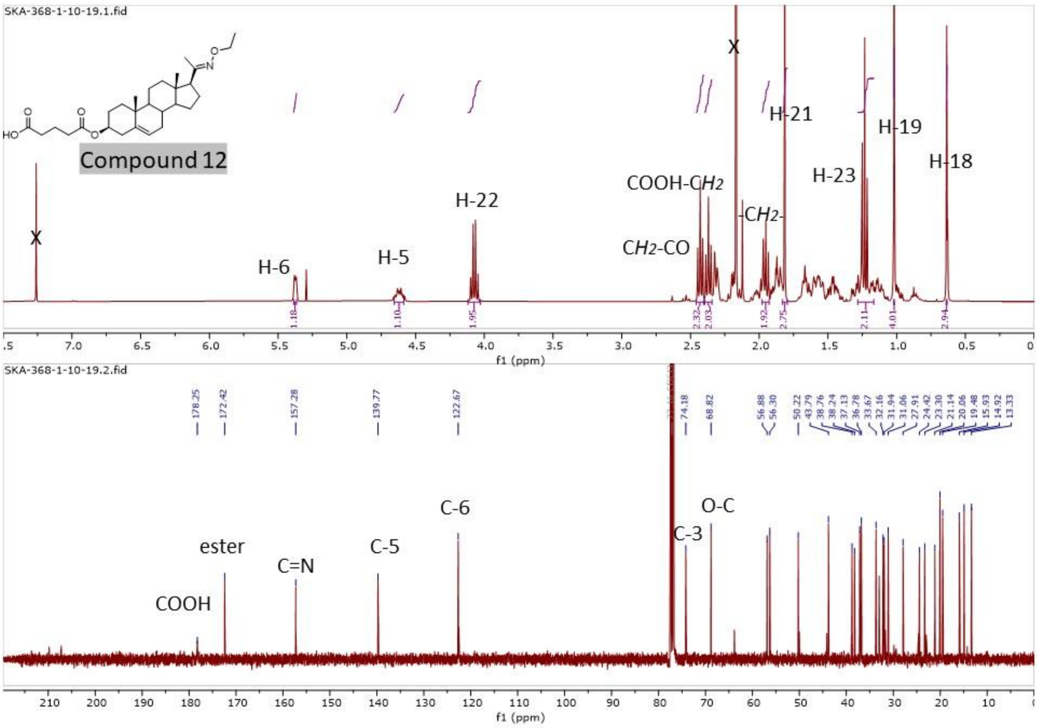

**Figure S3**. $^{1}$H NMR and $^{13}$C NMR and spectra of compound **13 (SKA-304)**

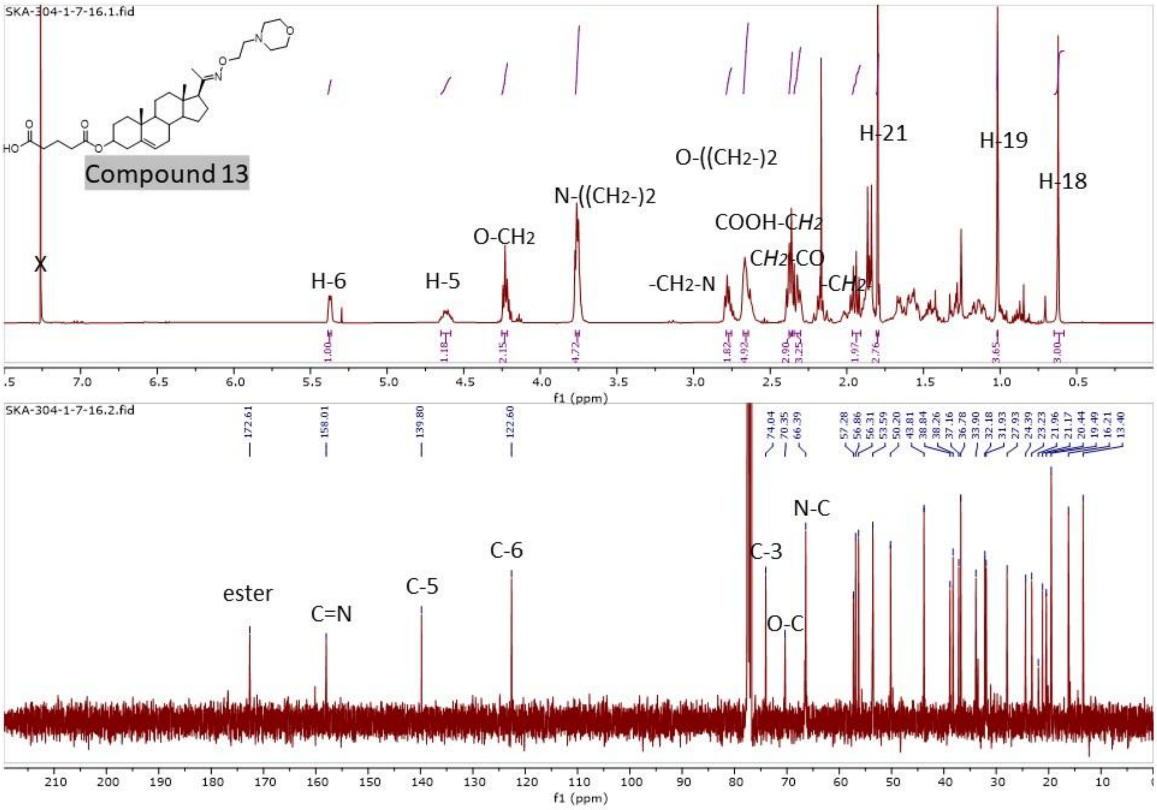


**Figure S4**. $^{1}$H NMR and $^{13}$C NMR and spectra of compound **14**

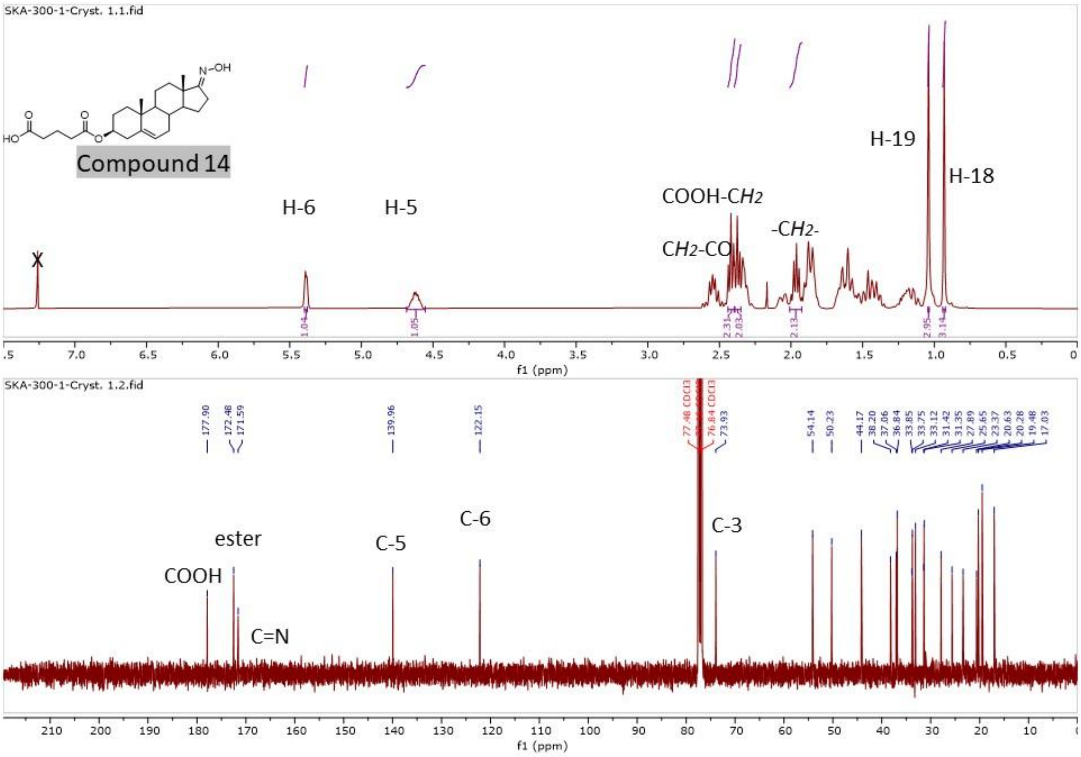

**Figure S5**. $^{1}H$ NMR and $^{13}C$ NMR and spectra of compound **15**

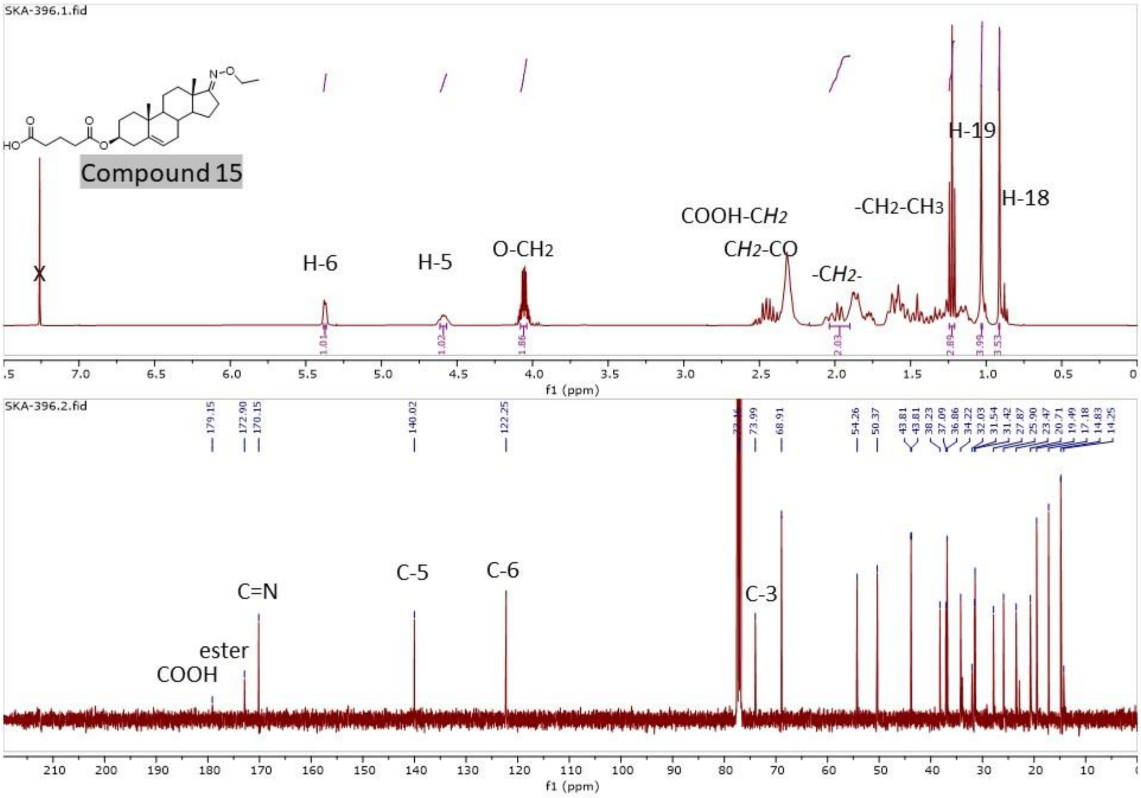


**Figure S6**. $^{1}H$ NMR and $^{13}C$ NMR and spectra of compound **16**

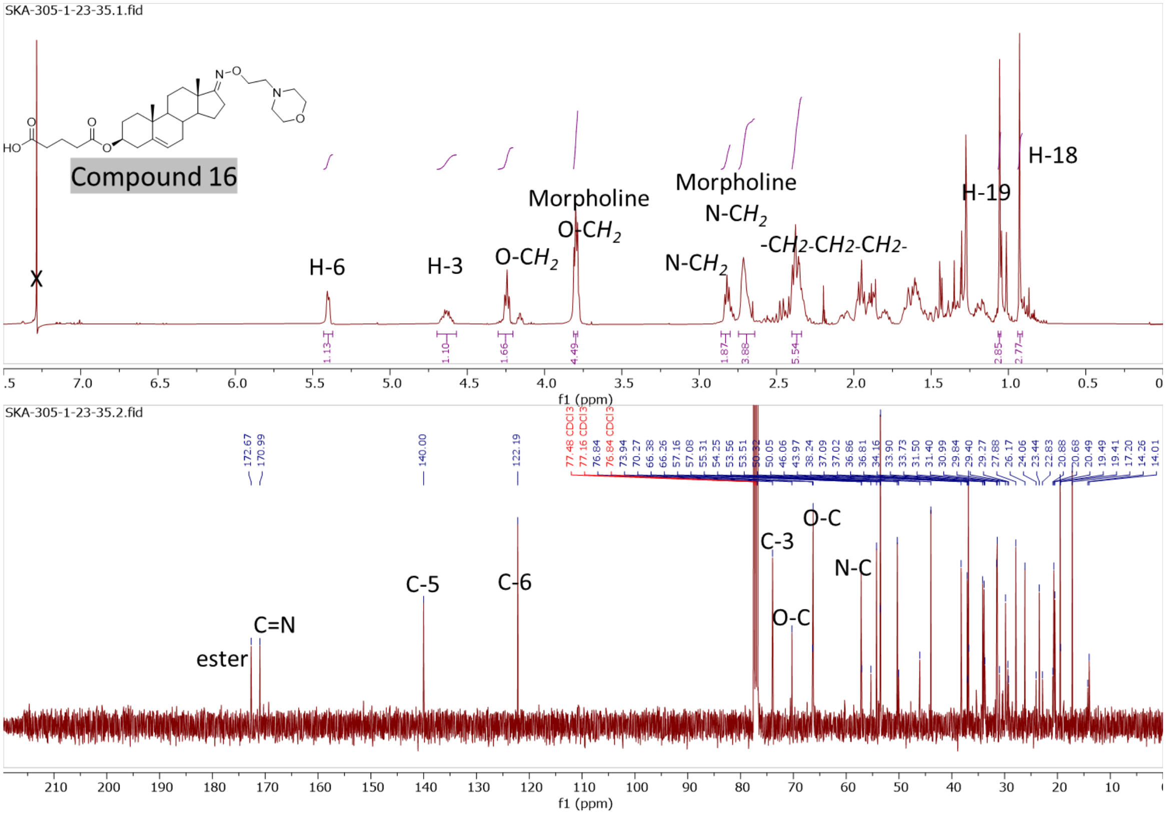

**Figure S7**. [1]H NMR and [13]C NMR and spectra of compound **17**

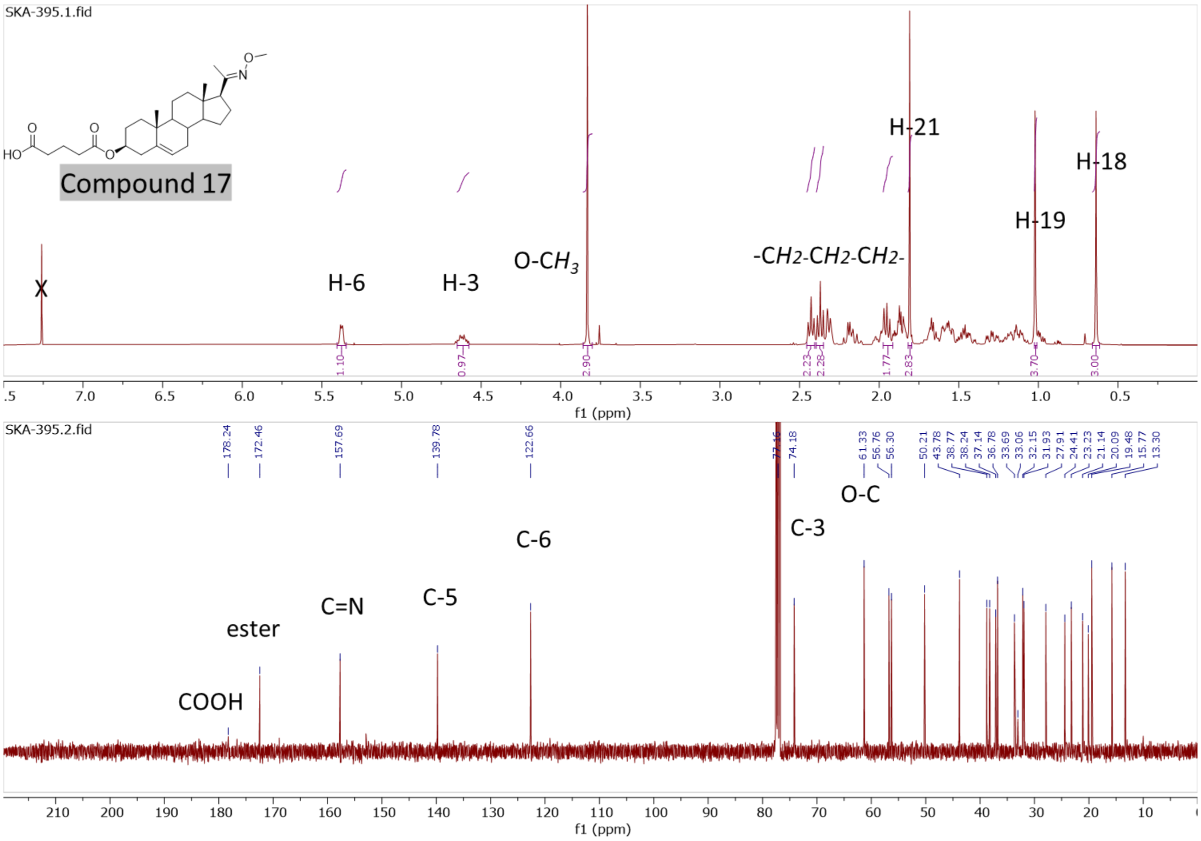


**Figure S8**. [1]H NMR and [13]C NMR and spectra of compound **18**

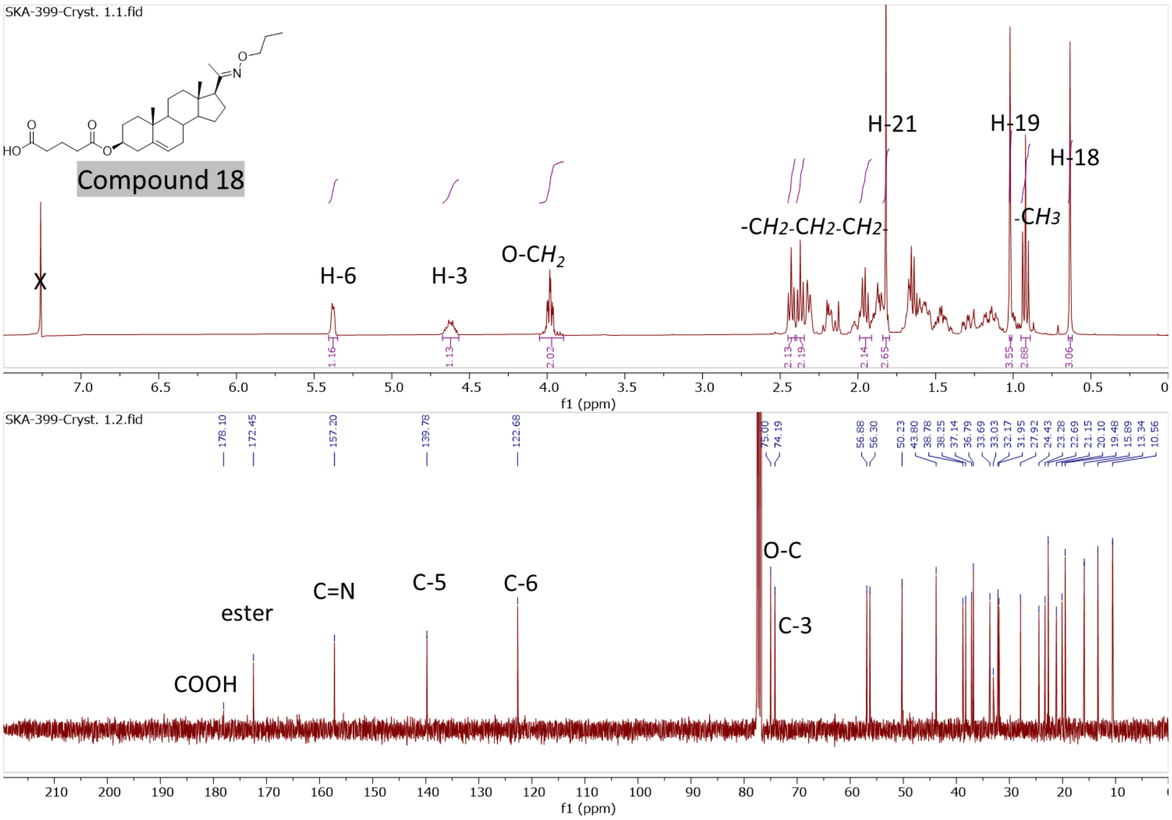

**Figure S9**. $^{1}$H NMR and $^{13}$C NMR and spectra of compound **19**

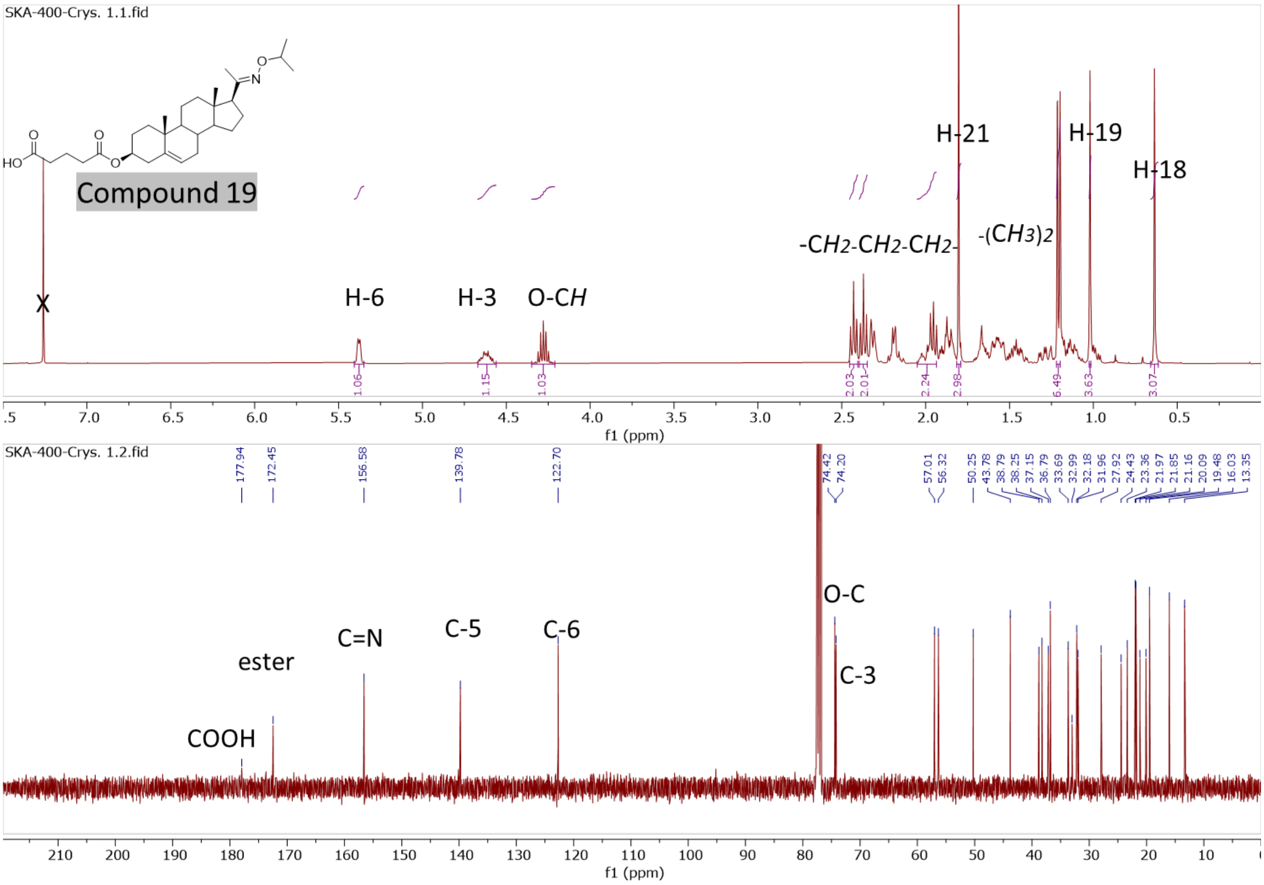


**Figure S10**. $^{1}$H NMR and $^{13}$C NMR and spectra of compound **20**

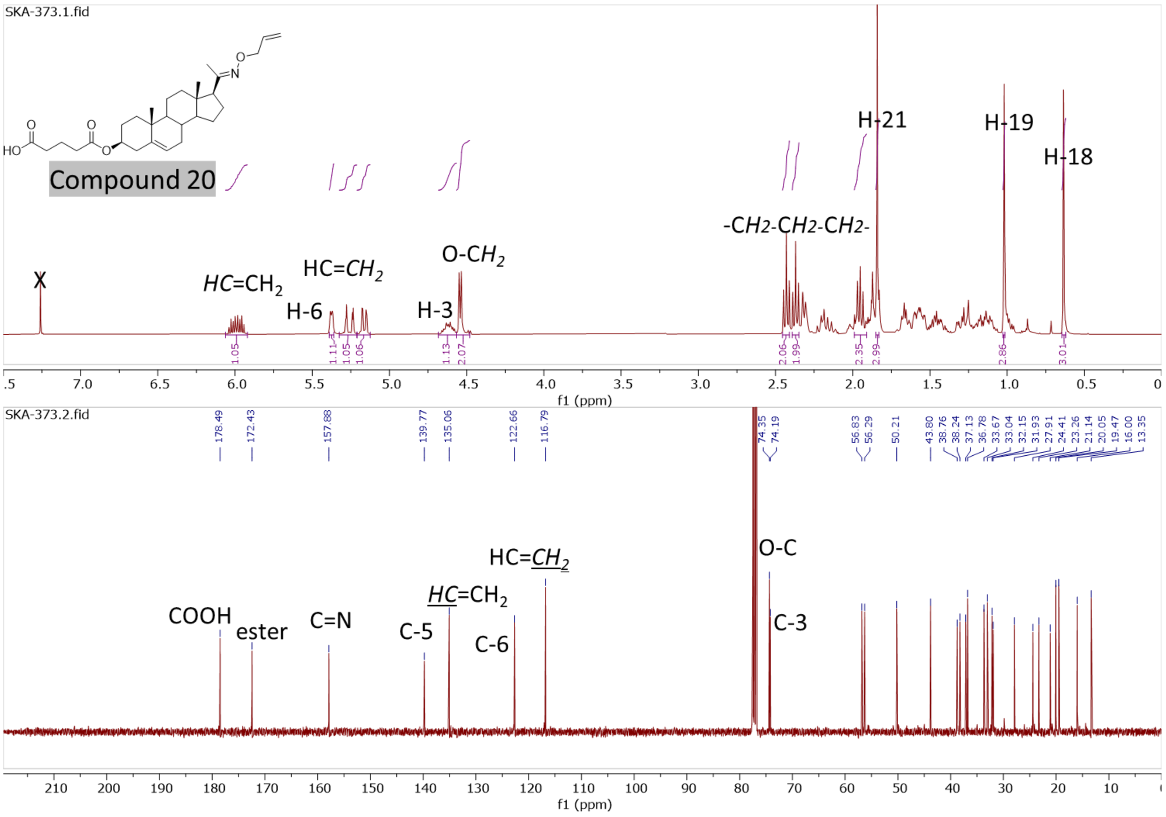

**Figure S11**. [1]H NMR and [13]C NMR and spectra of compound **21**

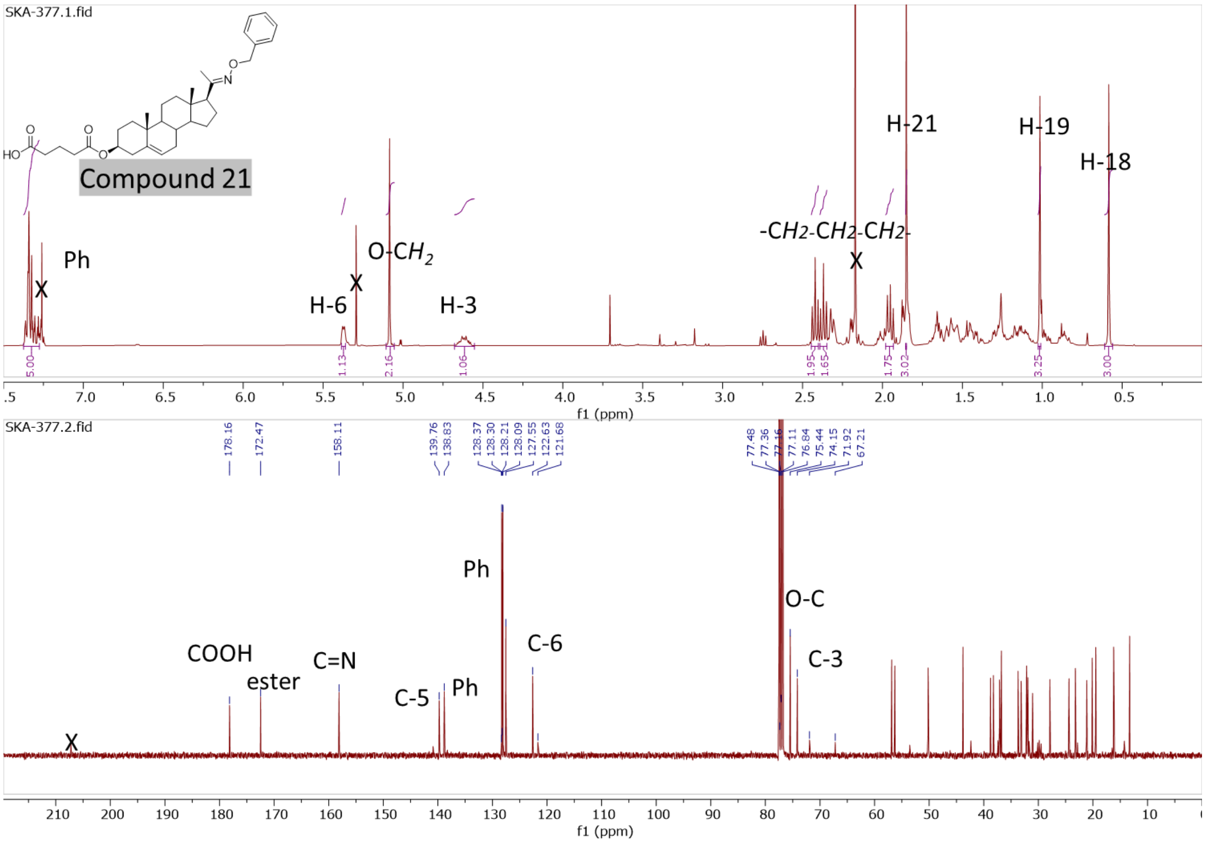


**Figure S12**. [1]H NMR and [13]C NMR and spectra of compound **22**

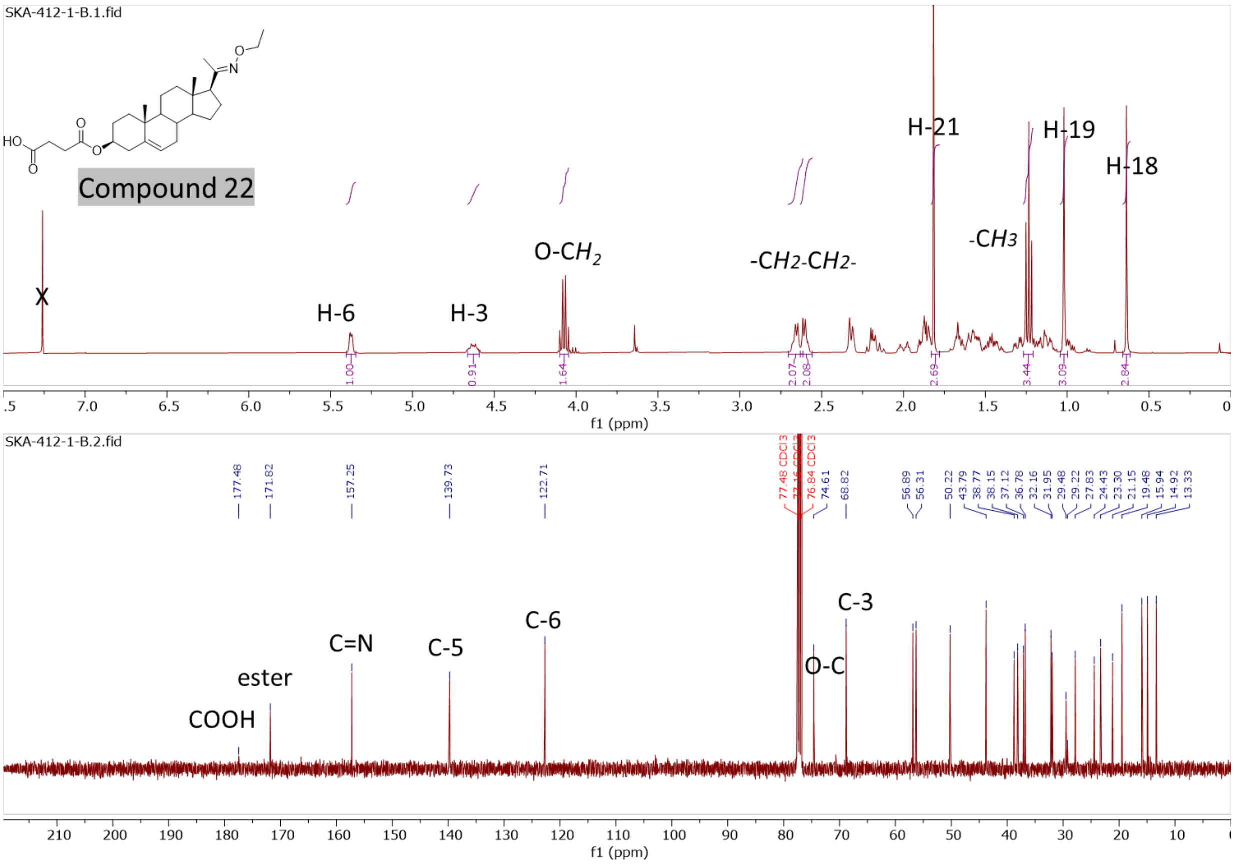

**Figure S13**. [1]H NMR and [13]C NMR and spectra of compound **23**

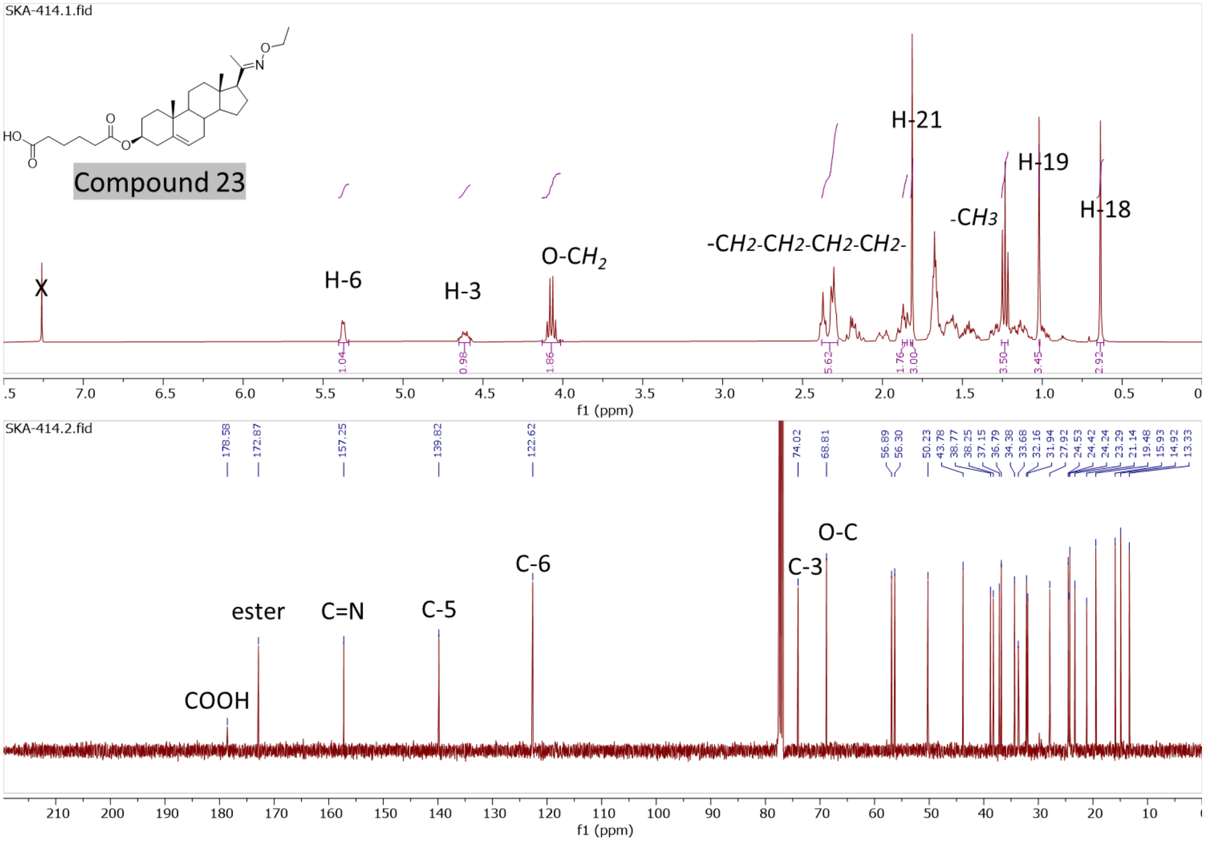


**Figure S14**. [1]H NMR and [13]C NMR and spectra of compound **24**

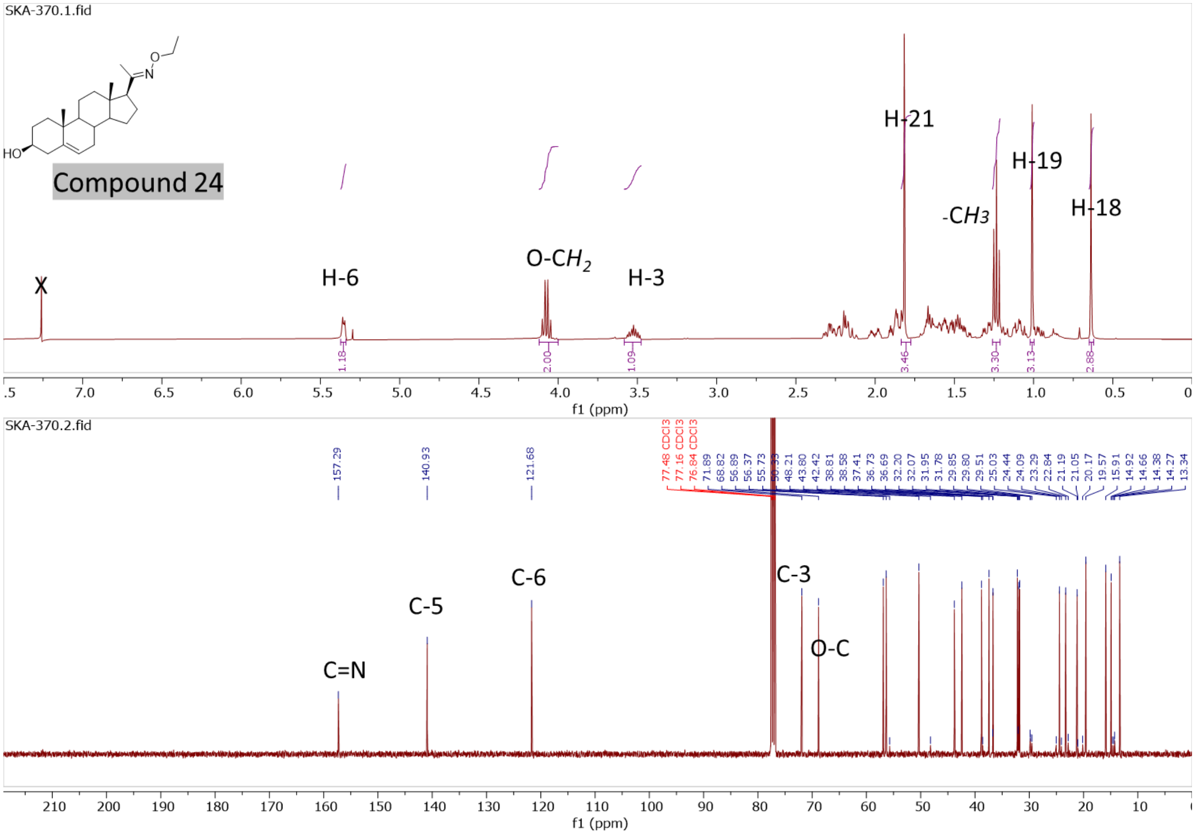

**Figure S15**. $^{1}$H NMR and $^{13}$C NMR and spectra of compound **25**

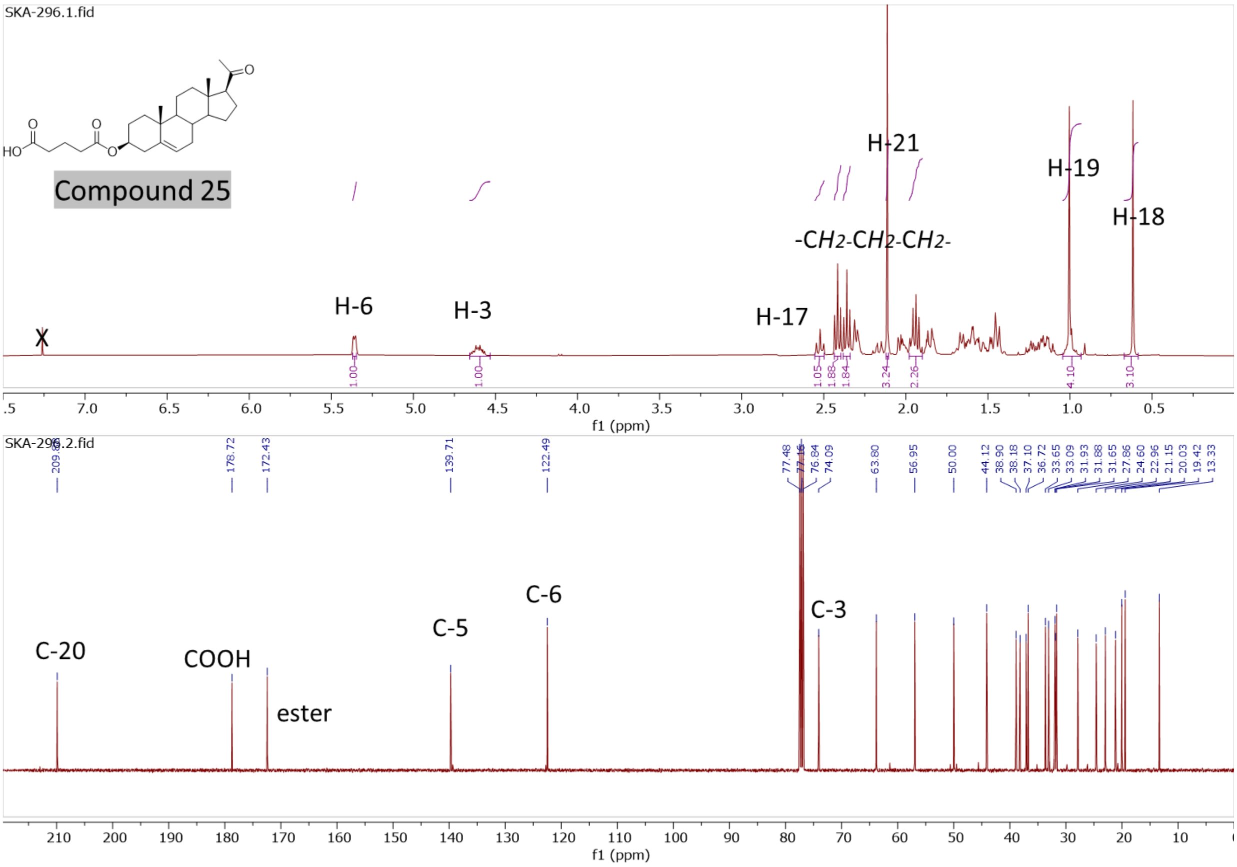


**Figure S16**. $^{1}$H NMR and $^{13}$C NMR and spectra of compound **26**

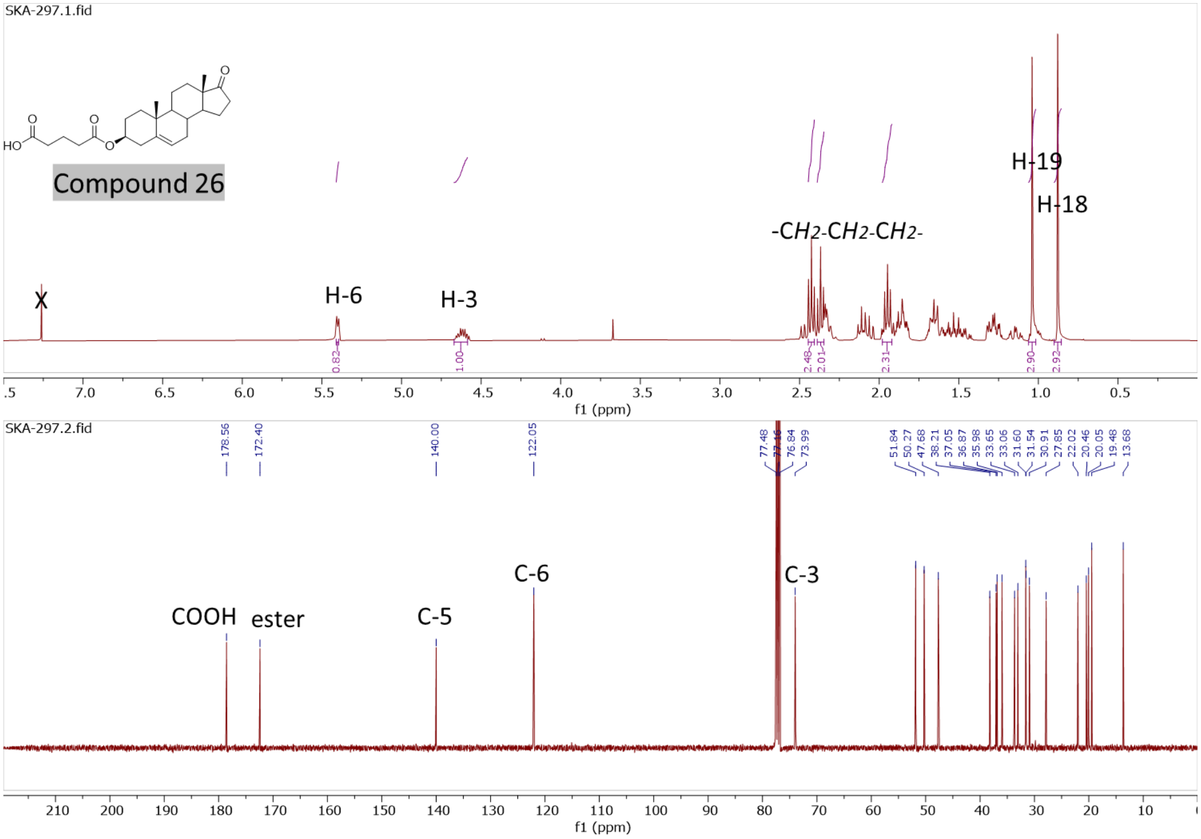

**Figure S17**. $^{1}$H NMR and $^{13}$C NMR and spectra of compound **27**

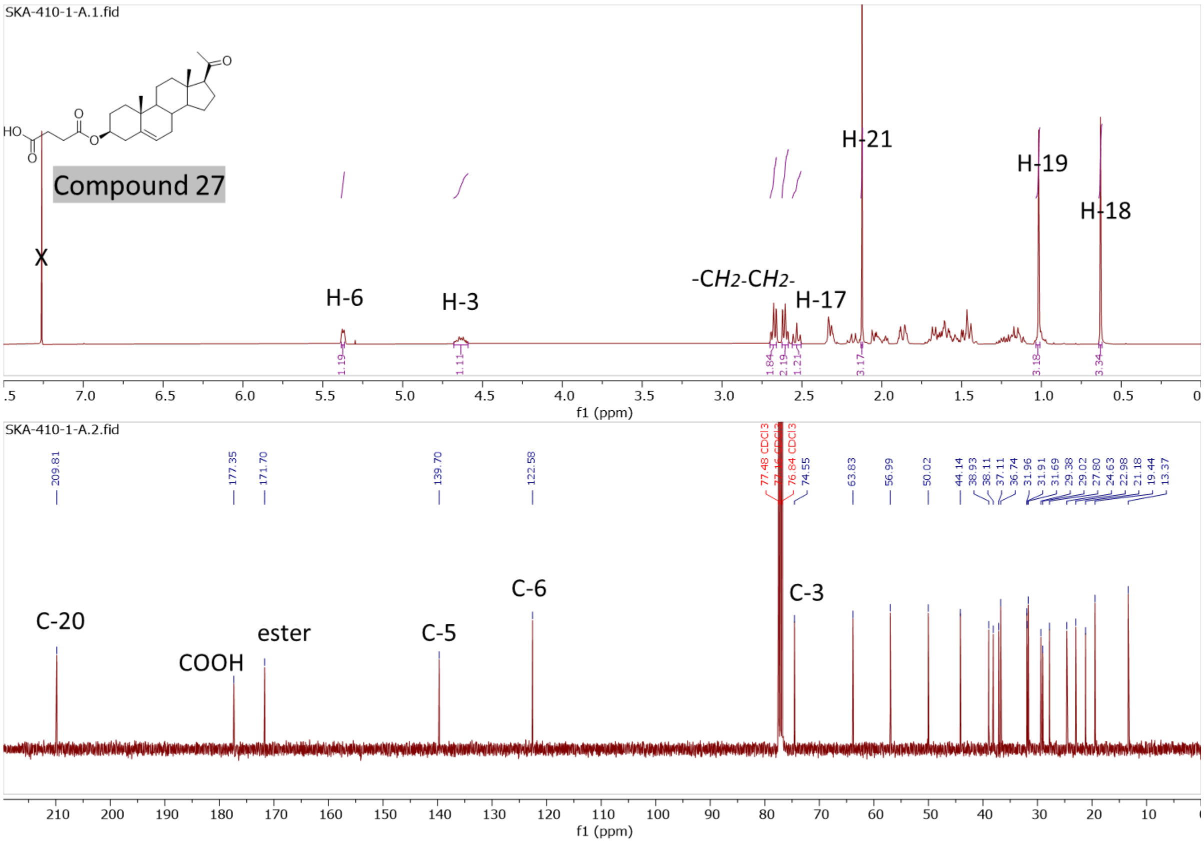


**Figure S18**. $^{1}$H NMR and $^{13}$C NMR and spectra of compound **28**

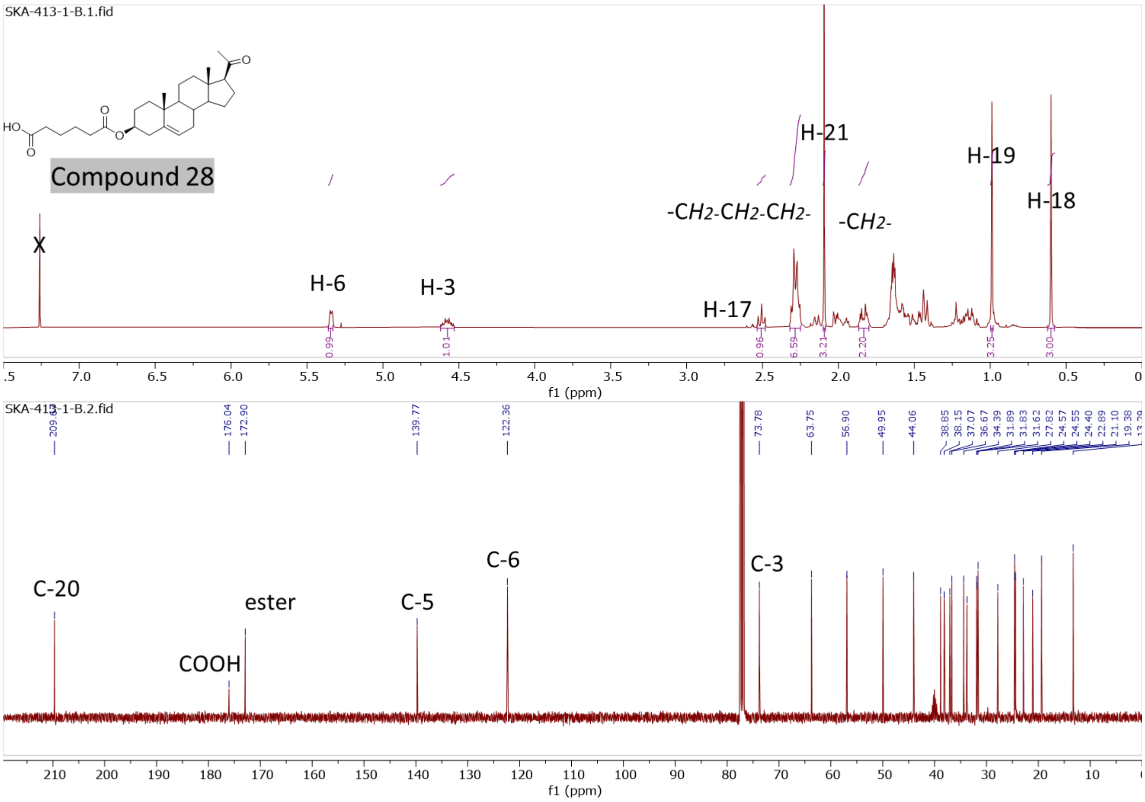

### HR-MS and LC-MS spectra of compounds **11**-**28**

**Figure S19**. HR-MS and LC-MS spectra of compound **11**

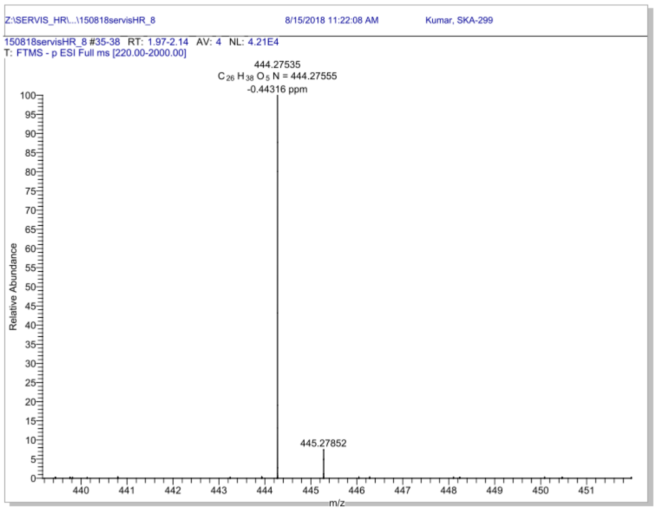


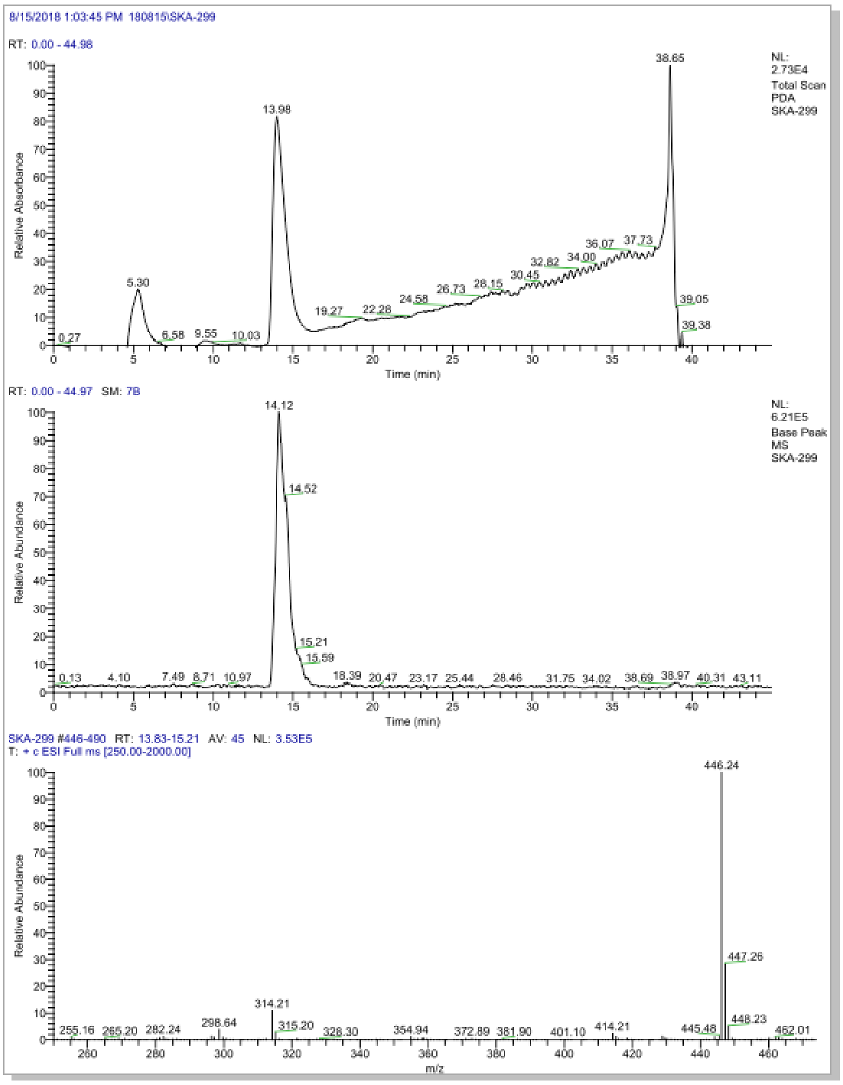

**Figure S20**. HR-MS and LC-MS spectra of compound **12**

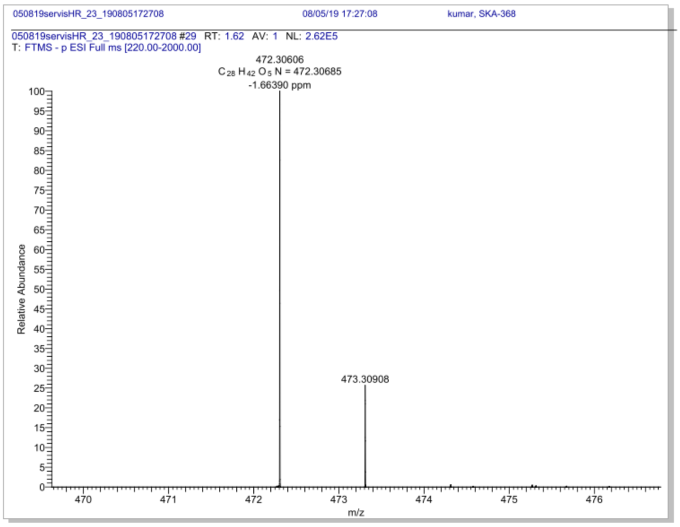


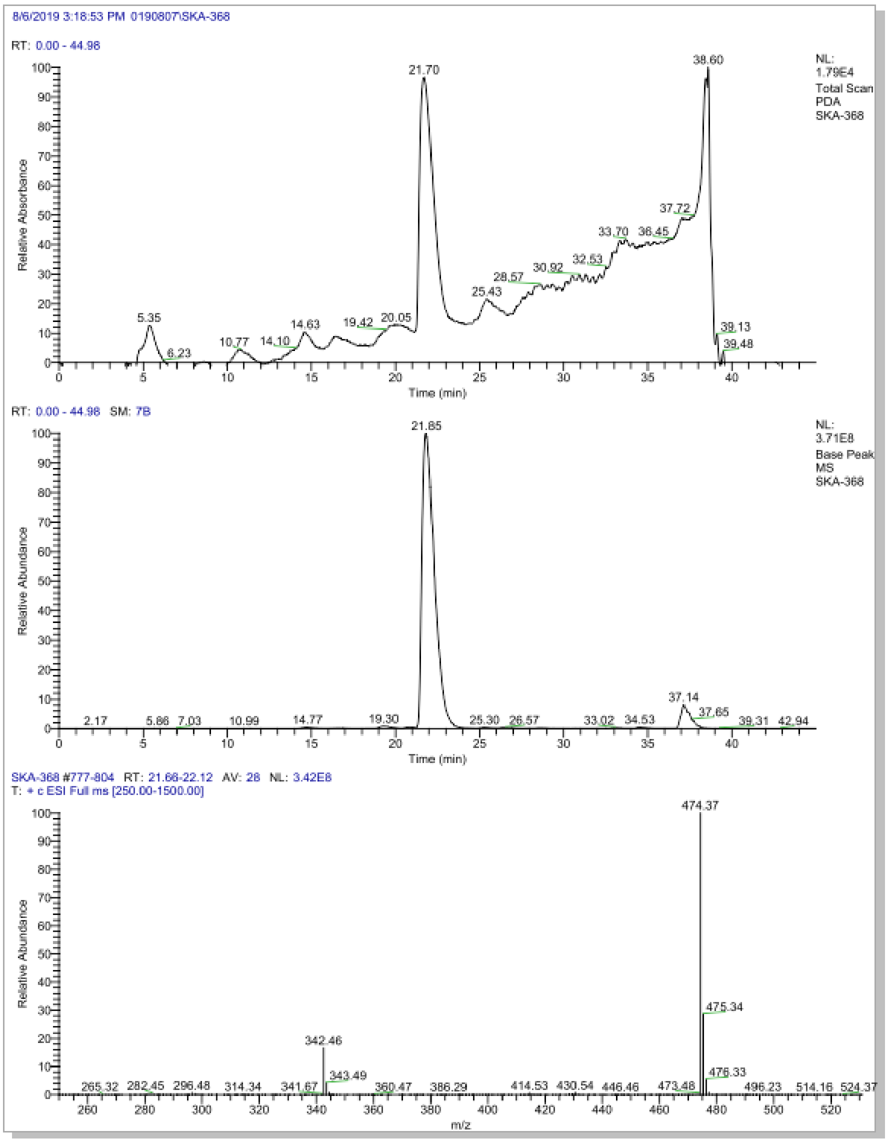

**Figure S21**. HR-MS and LC-MS spectra of compound **13**

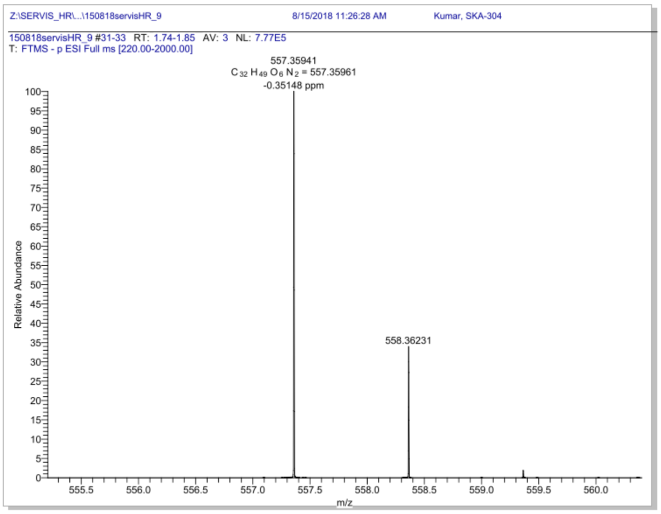


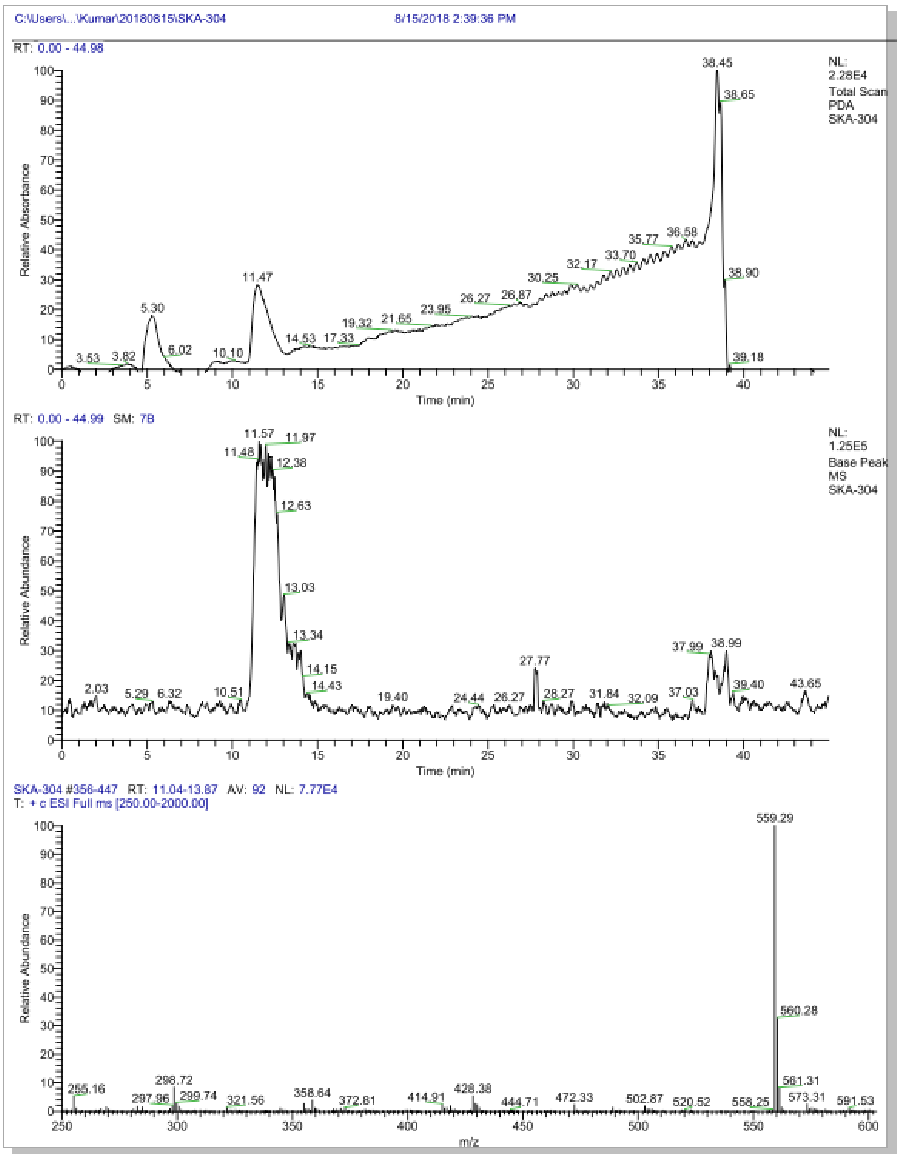

**Figure S22**. HR-MS and LC-MS spectra of compound **14**

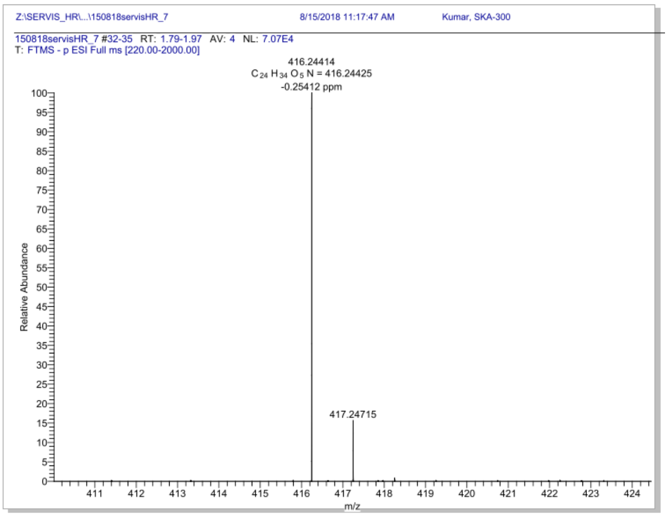


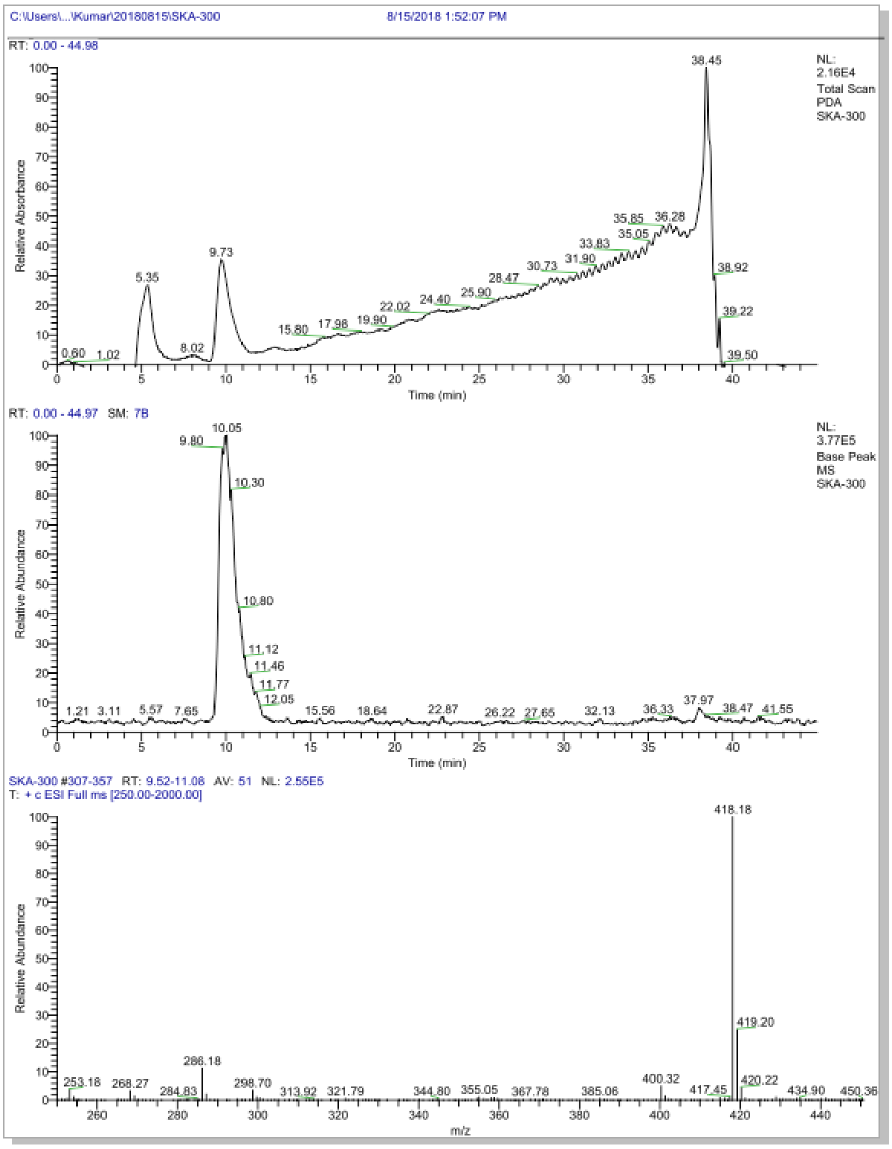

**Figure S23**. HR-MS and LC-MS spectra of compound **15**

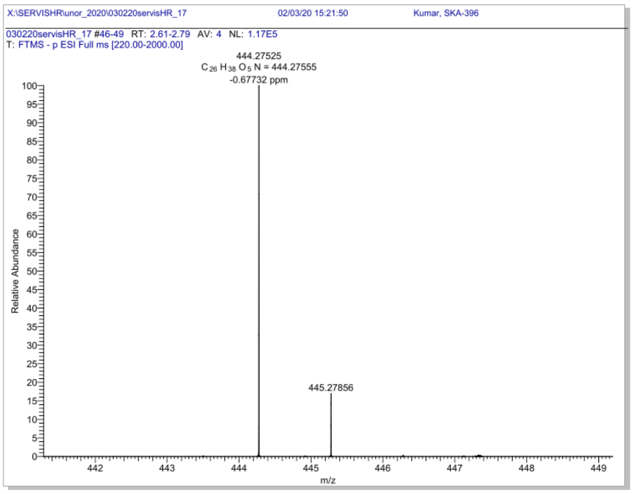


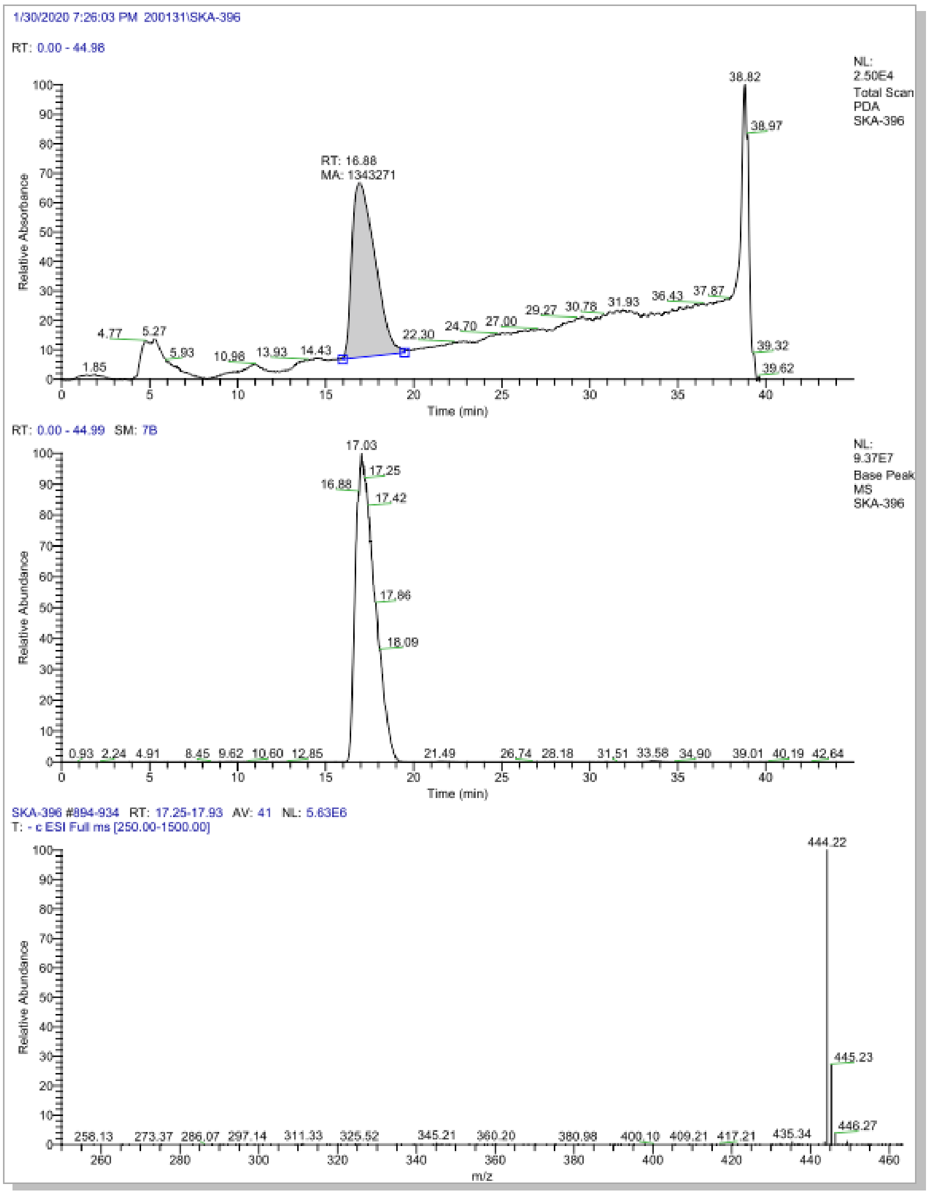

**Figure S24**. HR-MS and LC-MS spectra of compound **16**

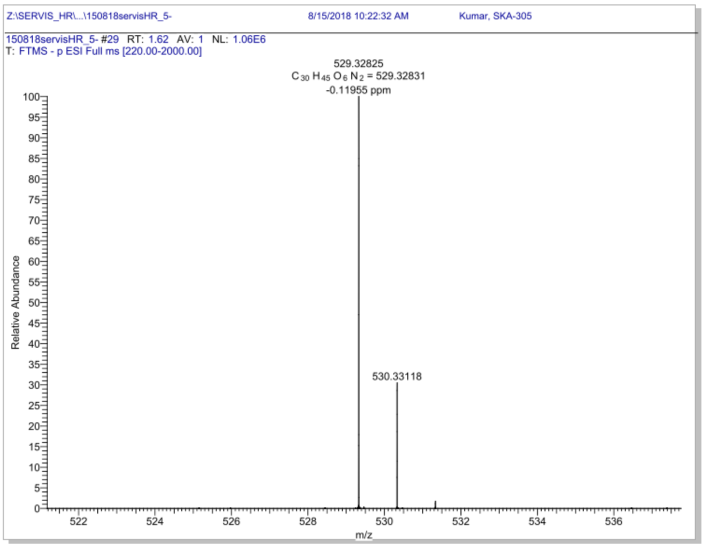


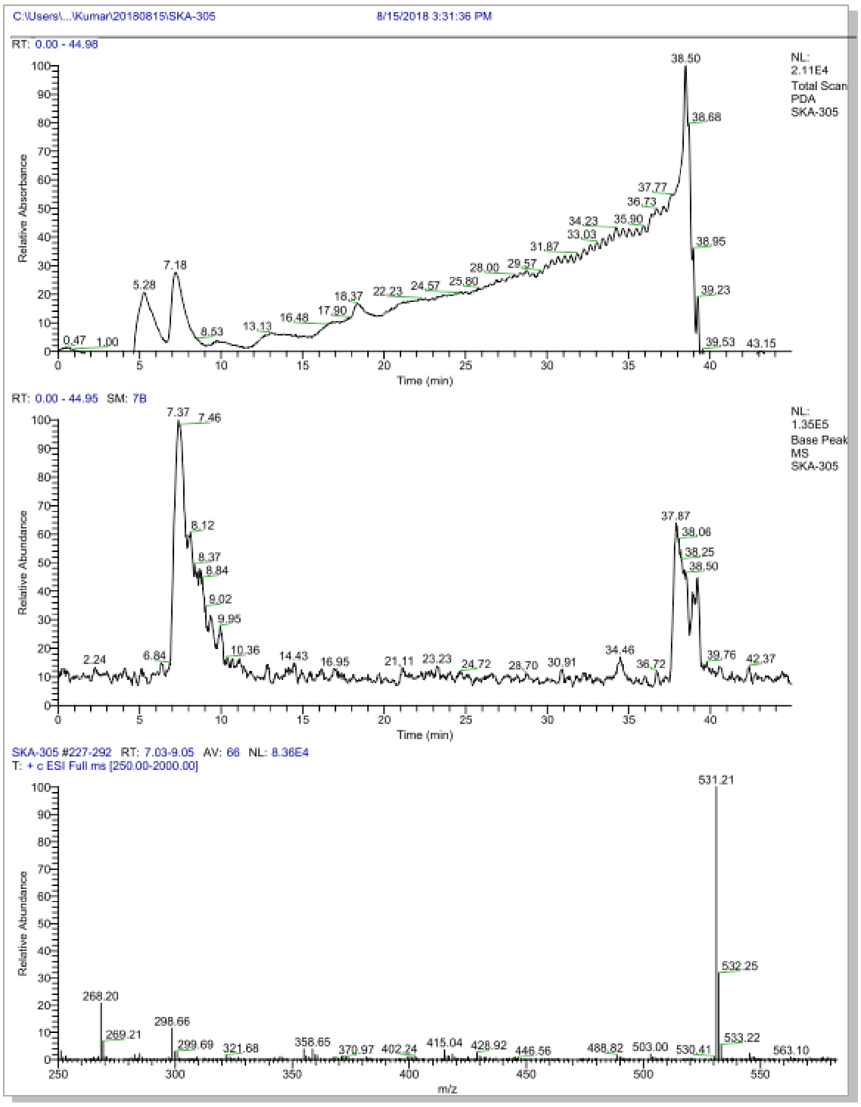

**Figure S25**. HR-MS and LC-MS spectra of compound **17**

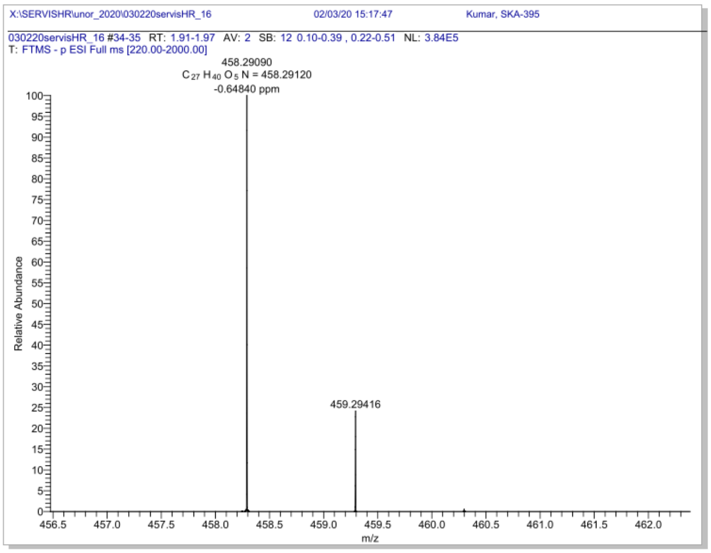


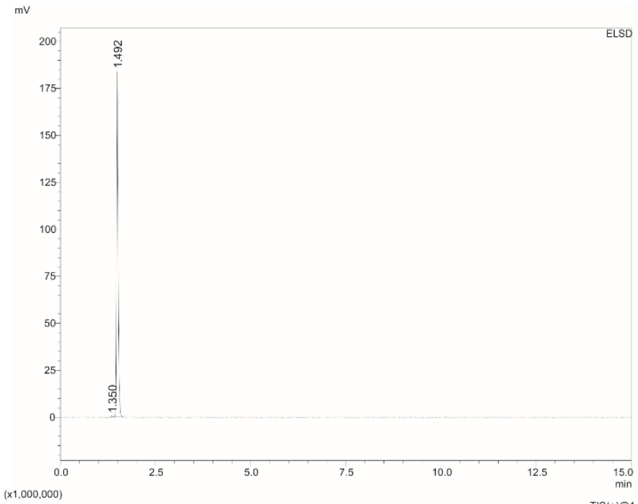


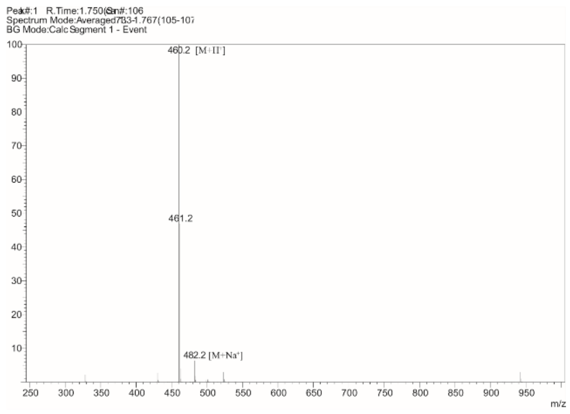


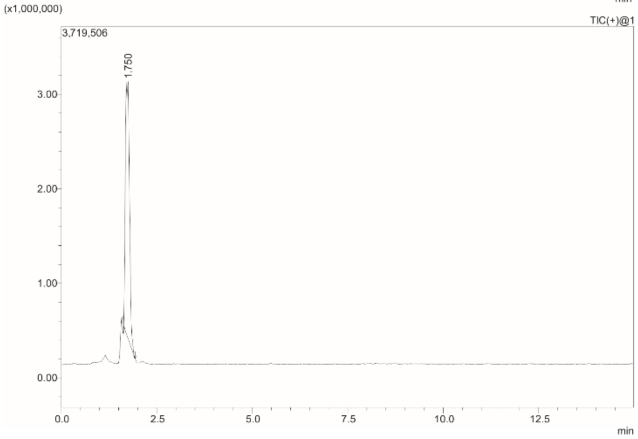


Box :
Position :
Method Filename : C8-80,20.lcm
Vial # : 1-2
Injection Volume : 0.3 uL
Date Acquired : 25-Aug-22 12:30:51

Comment : Isocratic, A: H2O, MeOH, FA (950:50:1); B: ACN; A:B (80:20), 0.5 mL/min
Column, Shim-pack Scepter, C8-120, 1.9 um, 2.1x100 mm, (Shimadzu)

ELSD

| Ret. Time | Area | Area% |
|---|---|---|
| 1.350 | 2255 | 0.410 |
| 1.492 | 548194 | 99.590 |
| | 550450 | 100.000 |

**Figure S26**. HR-MS and LC-MS spectra of compound **18**

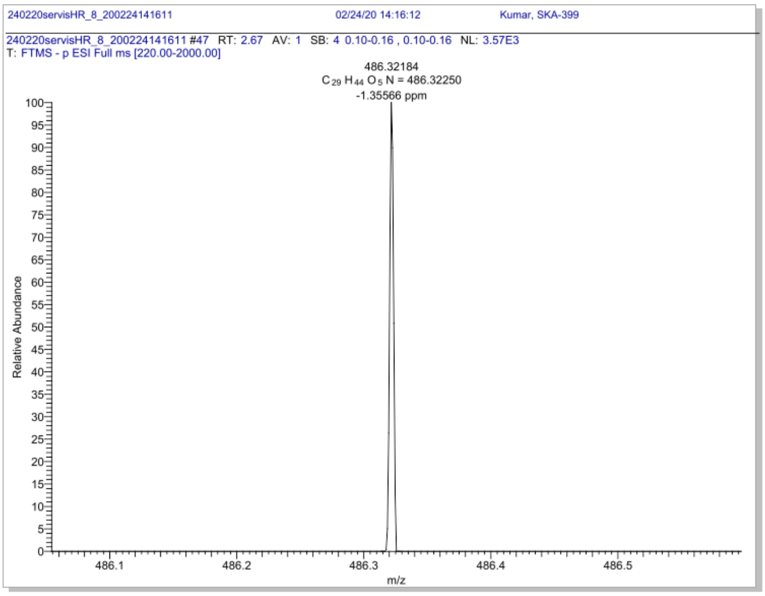


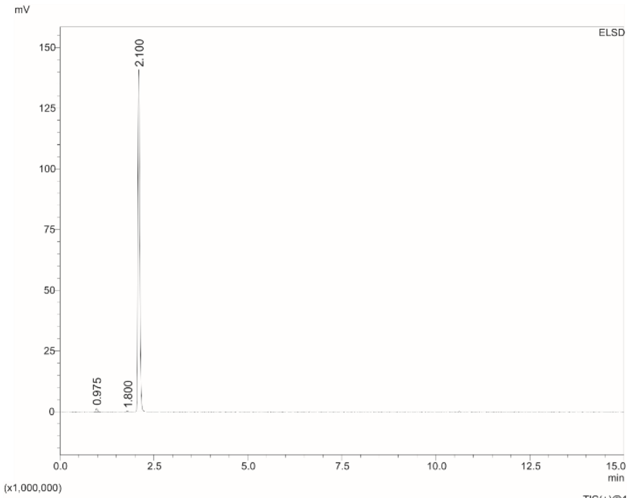


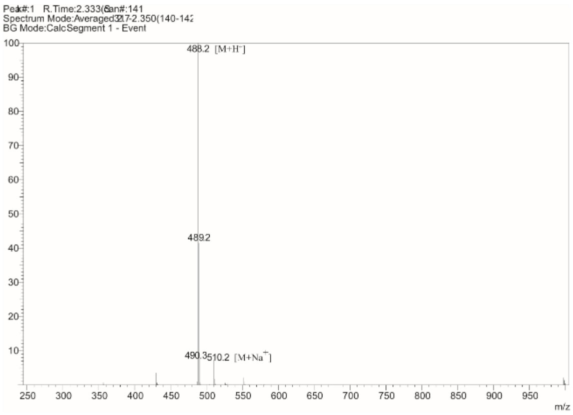


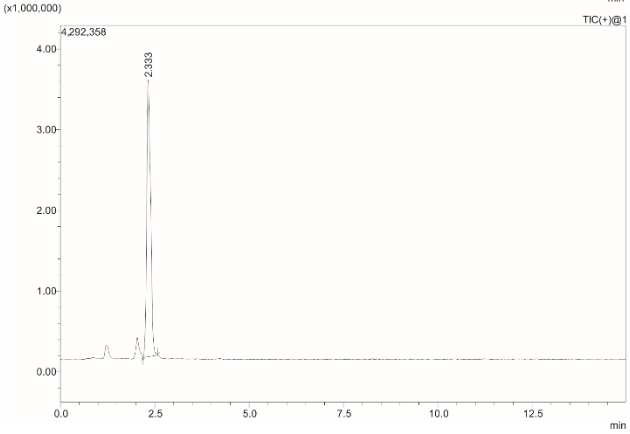


| | |
|---|---|
| Box | : |
| Position | : |
| Method Filename | : C8-80,20.lcm |
| Vial # | : 1-3 |
| Injection Volume | : 0.1 uL |
| Date Acquired | : 25-Aug-22 13:16:04 |
| Comment | : Isocratic, A: H2O, MeOH, FA (950:50:1); B: ACN; A:B (80:20), 0.5 mL/min Column, Shim-pack Scepter, C8-120, 1.9 um, 2.1x100 mm, (Shimadzu) |

ELSD

| Ret. Time | Area | Area% |
|---|---|---|
| 0.975 | 4439 | 0.918 |
| 1.800 | 868 | 0.179 |
| 2.100 | 478375 | 98.903 |
| | 483681 | 100.000 |

**Figure S27**. HR-MS and LC-MS spectra of compound **19**

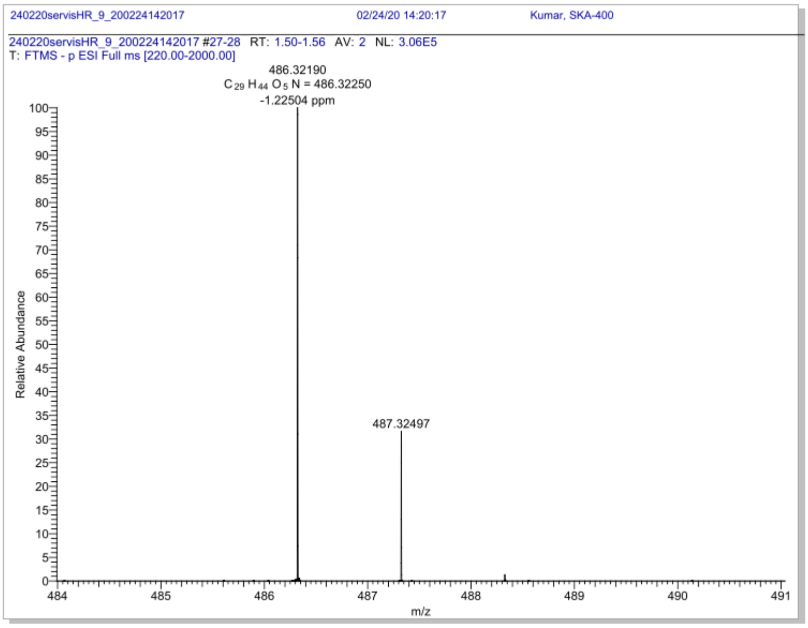


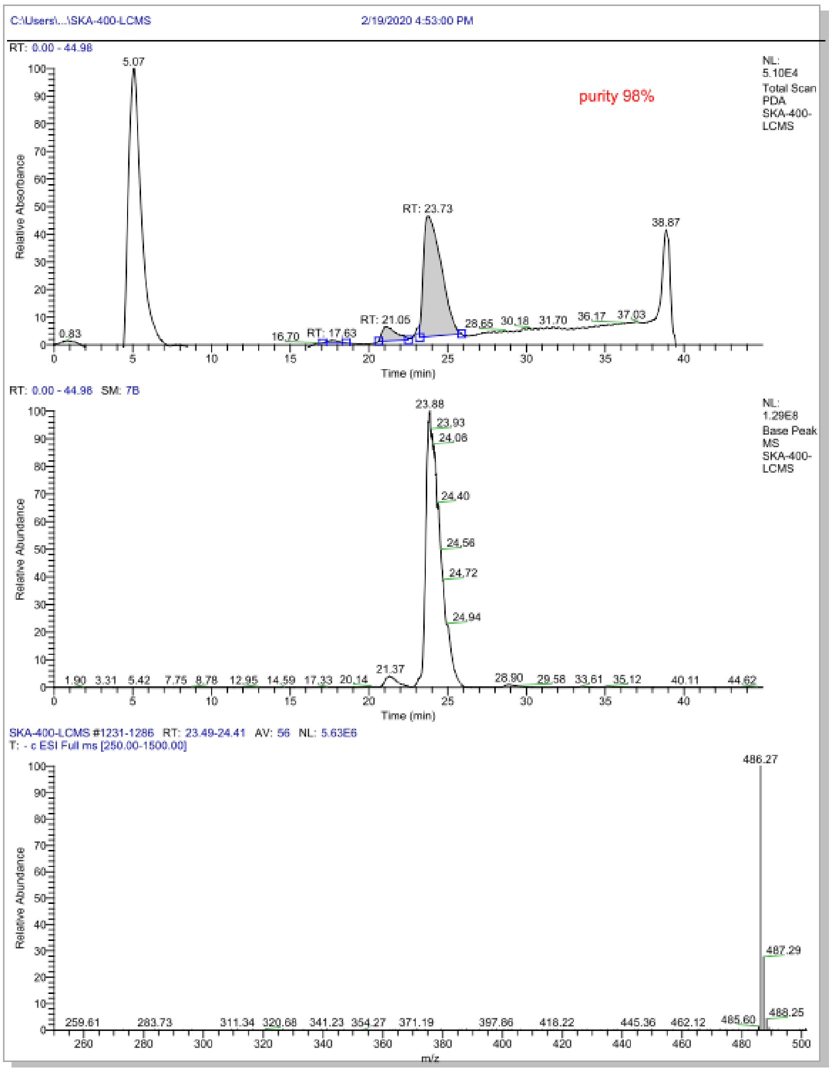

**Figure S28**. HR-MS and LC-MS spectra of compound **20**

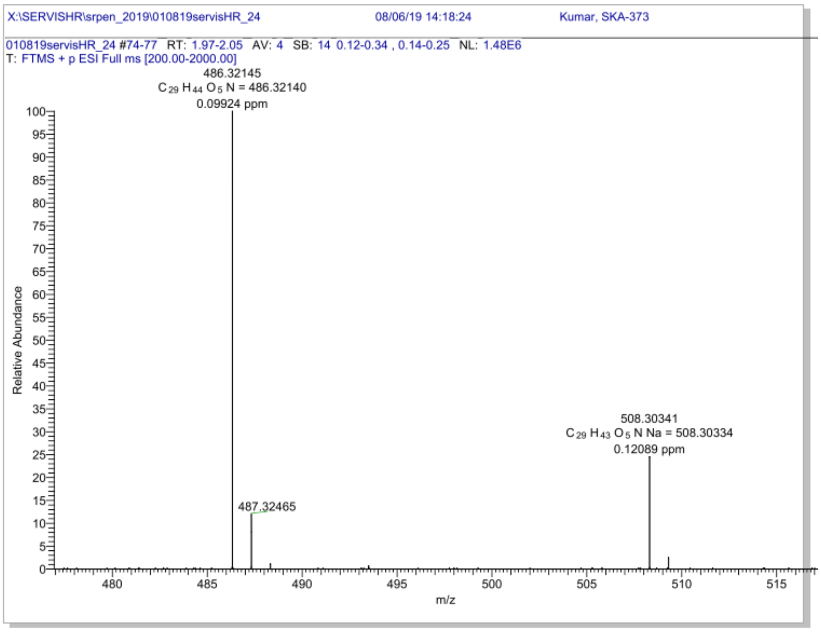


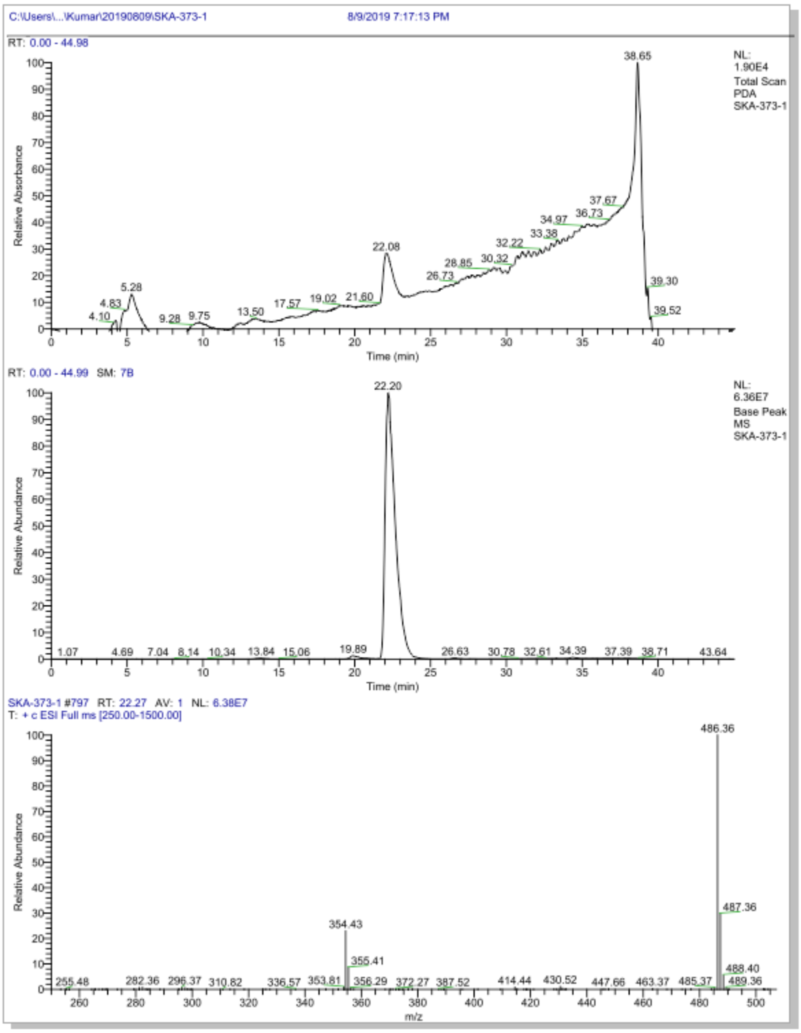

**Figure S29**. HR-MS and LC-MS spectra of compound **21**

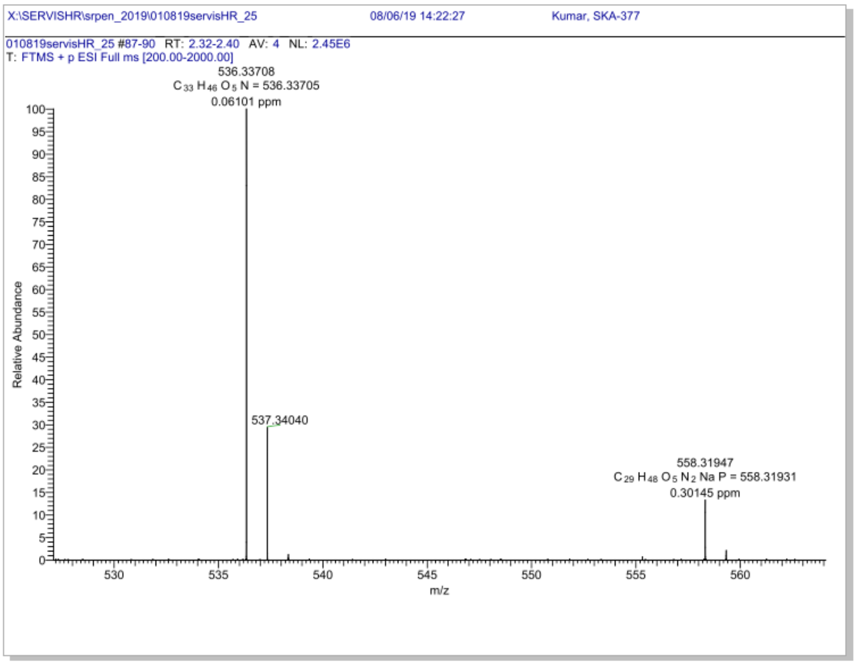


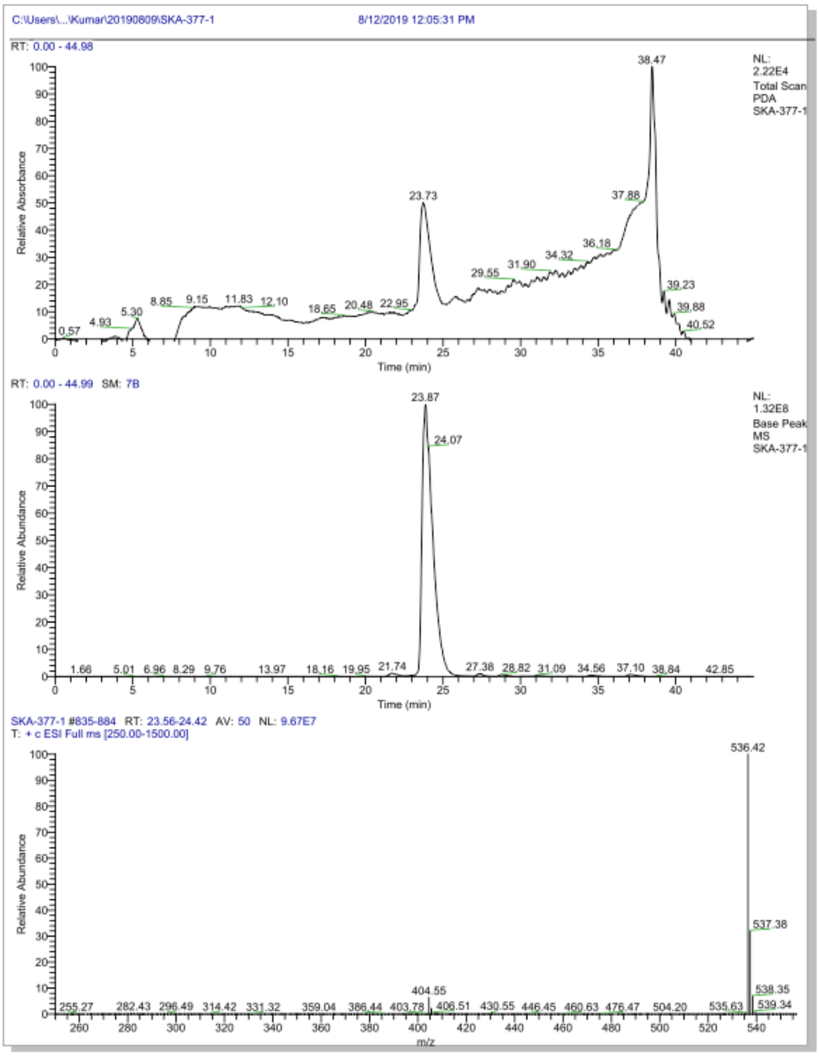

**Figure S30**. HR-MS and LC-MS spectra of compound **22**

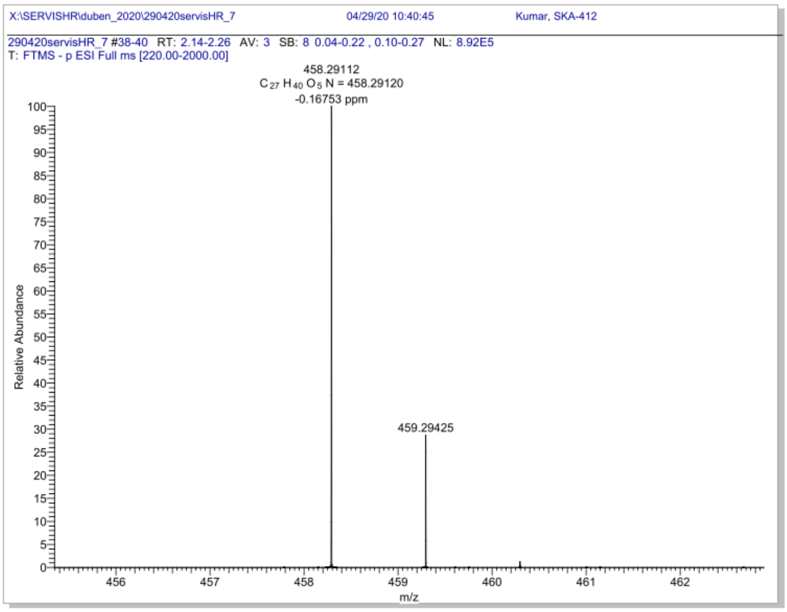


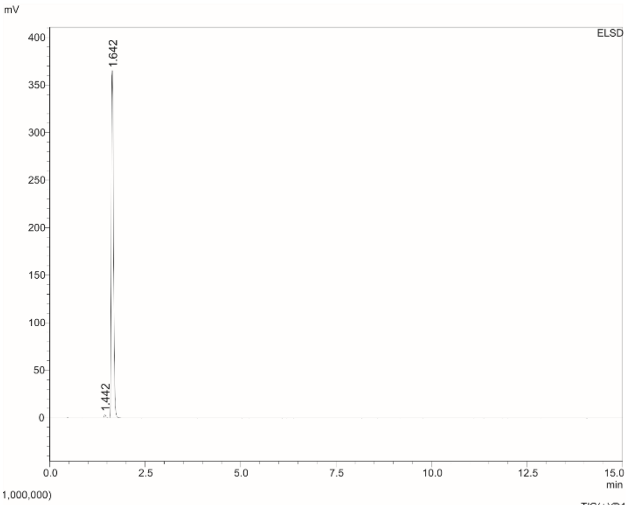


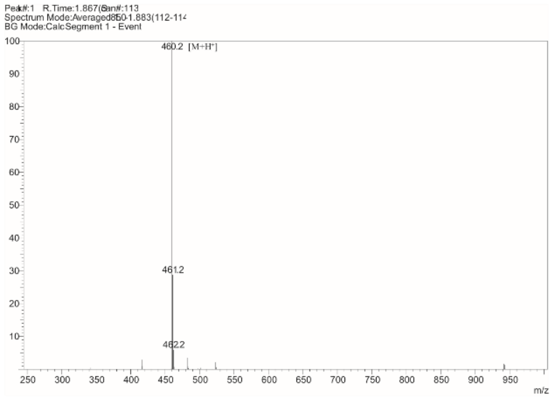


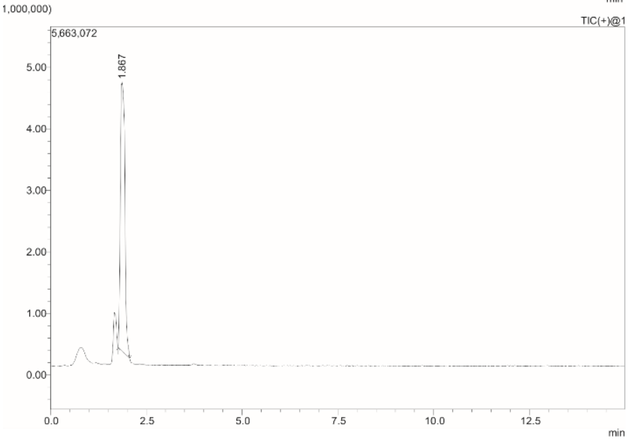


Box :
Position :
Method Filename : C8-80,20.lcm
Vial # : 1-4
Injection Volume : 0.3 uL
Date Acquired : 25-Aug-22 13:31:26

Comment : Isocratic, A: H2O, MeOH, FA (950:50:1); B: ACN; A:B (80:20), 0.5 mL/min Column, Shim-pack Scepter, C8-120, 1.9 um, 2.1x100 mm, (Shimadzu)

ELSD

| Ret. Time | Area | Area% |
|---|---|---|
| 1.442 | 10867 | 0.718 |
| 1.642 | 1502855 | 99.282 |
| | 1513722 | 100.000 |

**Figure S31**. HR-MS and LC-MS spectra of compound **23**

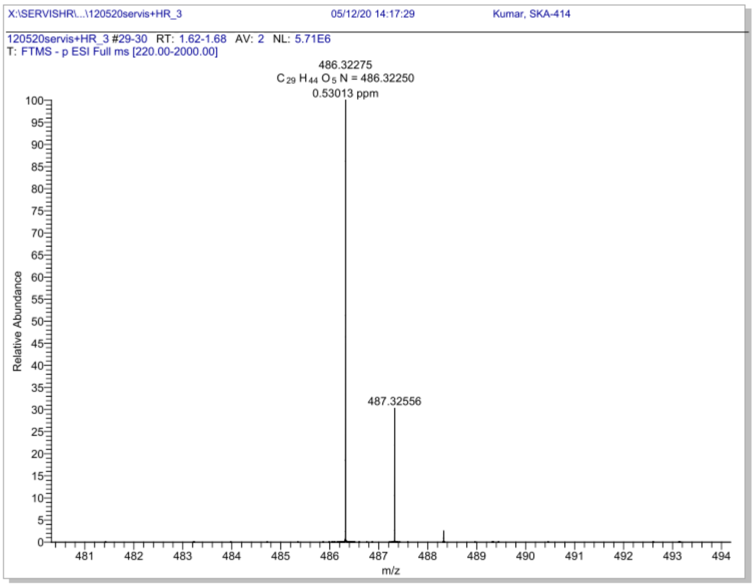


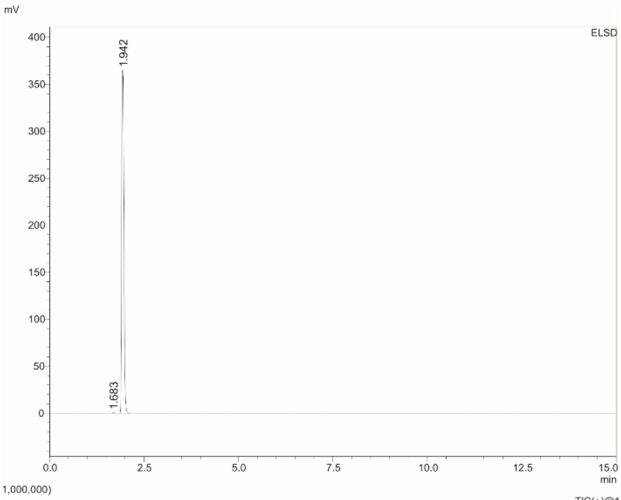


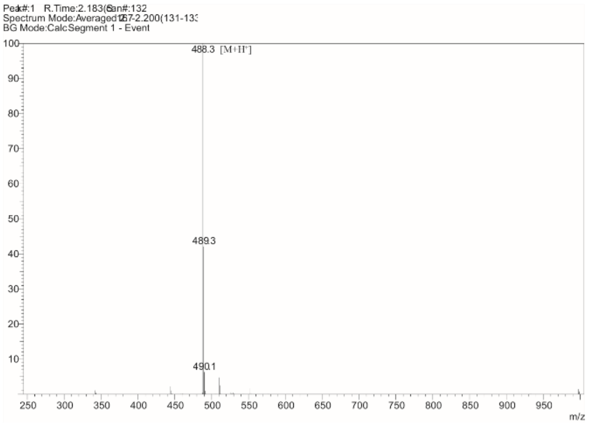


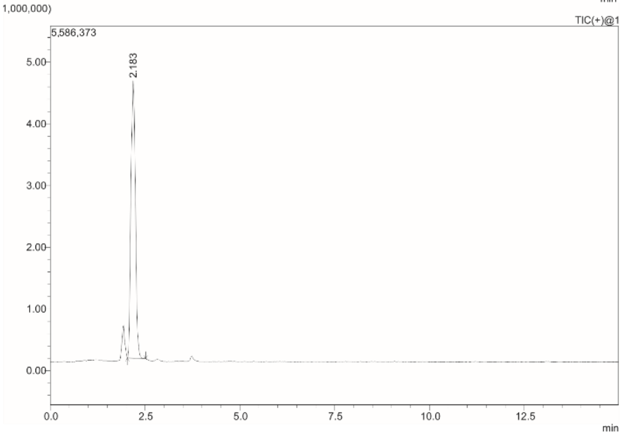


Box :
Position :
Method Filename : C8-80,20.lcm
Vial # : 1-5
Injection Volume : 0.3 uL
Date Acquired : 25-Aug-22 13:46:49

Comment : Isocratic, A: H2O, MeOH, FA (950:50:1); B: ACN; A:B (80:20), 0.5 mL/min
Column, Shim-pack Scepter, C8-120, 1.9 um, 2.1x100 mm, (Shimadzu)

ELSD

| Ret. Time | Area | Area% |
|---|---|---|
| 1.683 | 2353 | 0.157 |
| 1.942 | 1498508 | 99.843 |
| | 1500861 | 100.000 |

**Figure S32**. HR-MS and LC-MS spectra of compound **24**

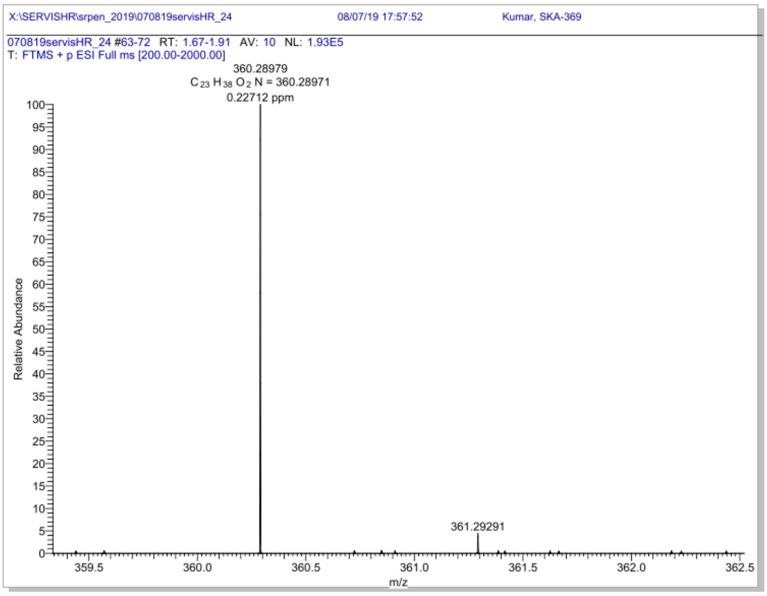


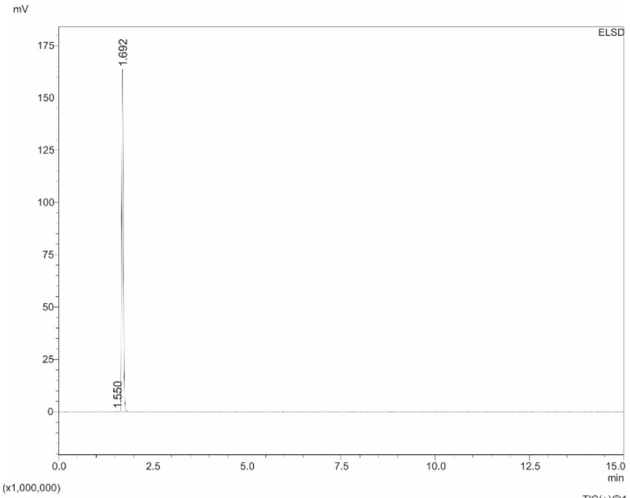


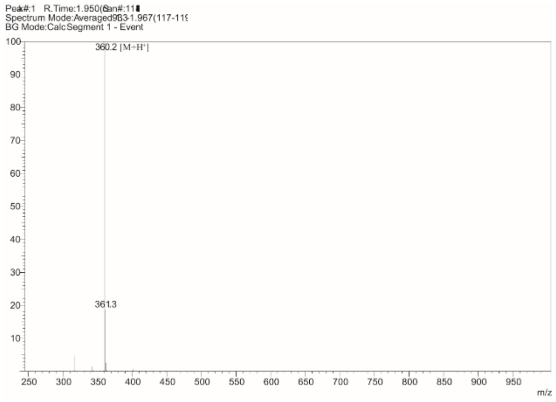


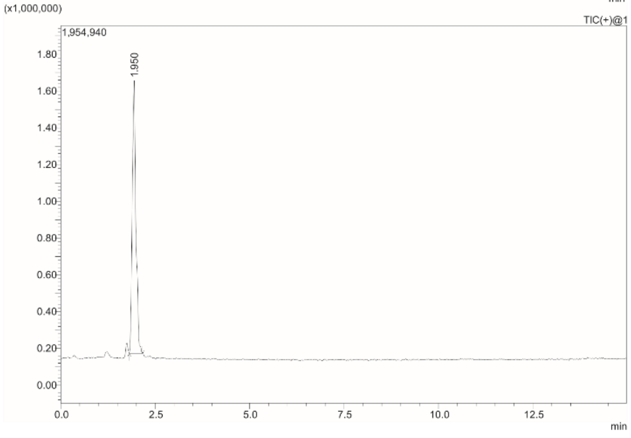


Box :
Position :
Method Filename : C8-80,20.lcm
Vial # : 1-1
Injection Volume : 0.4 uL
Date Acquired : 25-Aug-22 12:15:28

Comment : Isocratic, A: H2O, MeOH, FA (950:50:1); B: ACN; A:B (80:20), 0.5 mL/min Column, Shim-pack Scepter, C8-120, 1.9 um, 2.1x100 mm, (Shimadzu)

ELSD

| Ret. Time | Area | Area% |
|---|---|---|
| 1.550 | 215 | 0.044 |
| 1.692 | 493433 | 99.956 |
| | 493648 | 100.000 |

**Figure S33**. HR-MS and LC-MS spectra of compound **25**

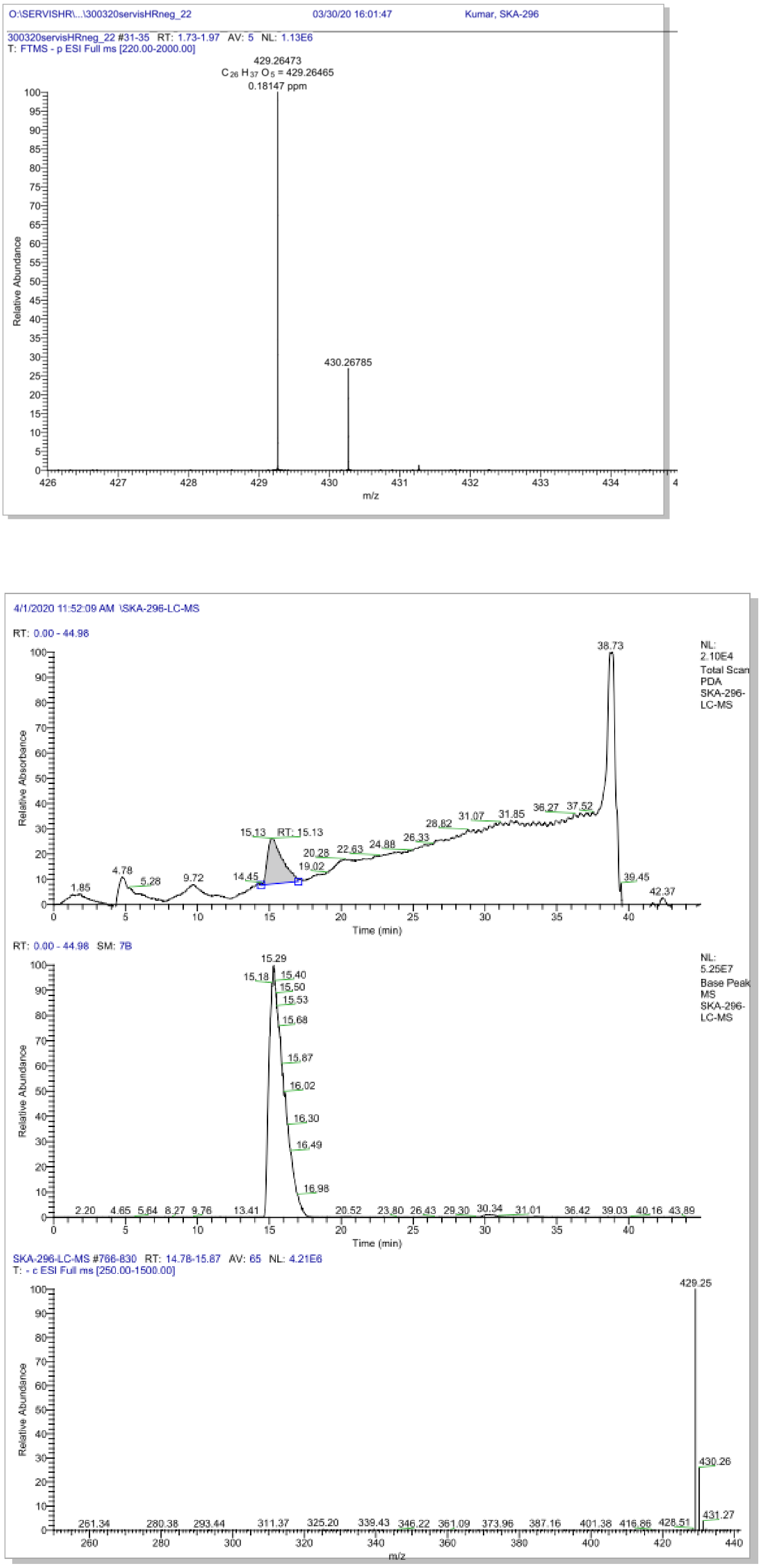

**Figure S34**. HR-MS and LC-MS spectra of compound **26**

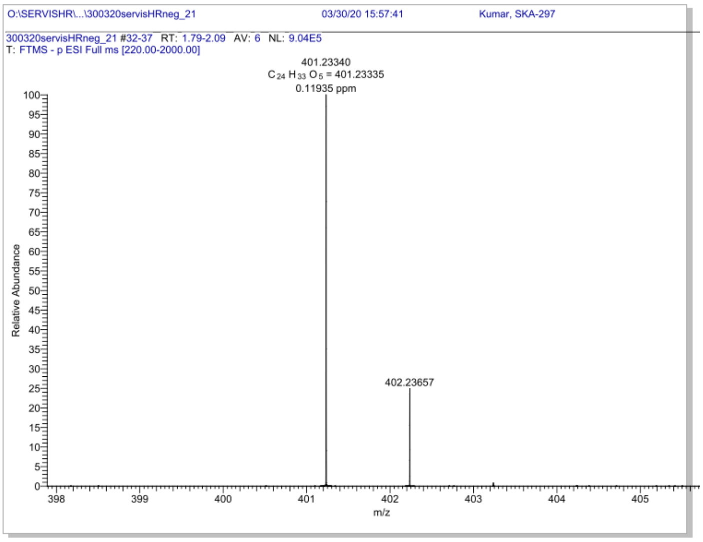


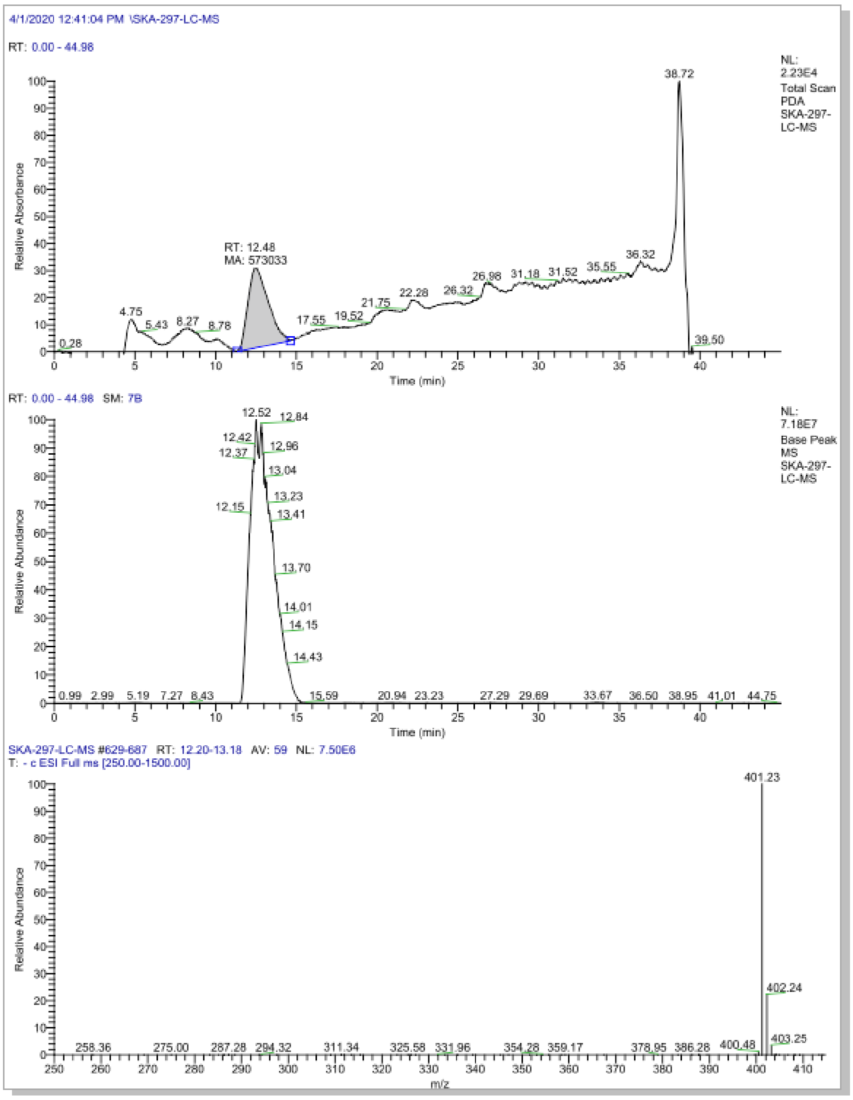

**Figure S35**. HR-MS and LC-MS spectra of compound **27**

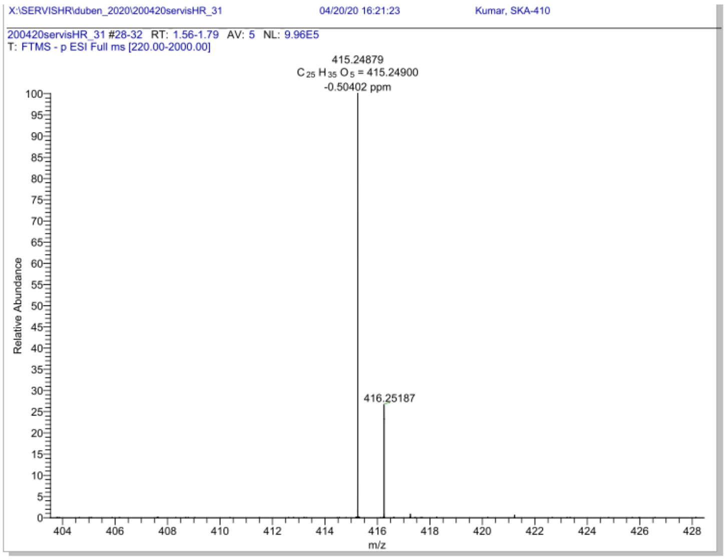


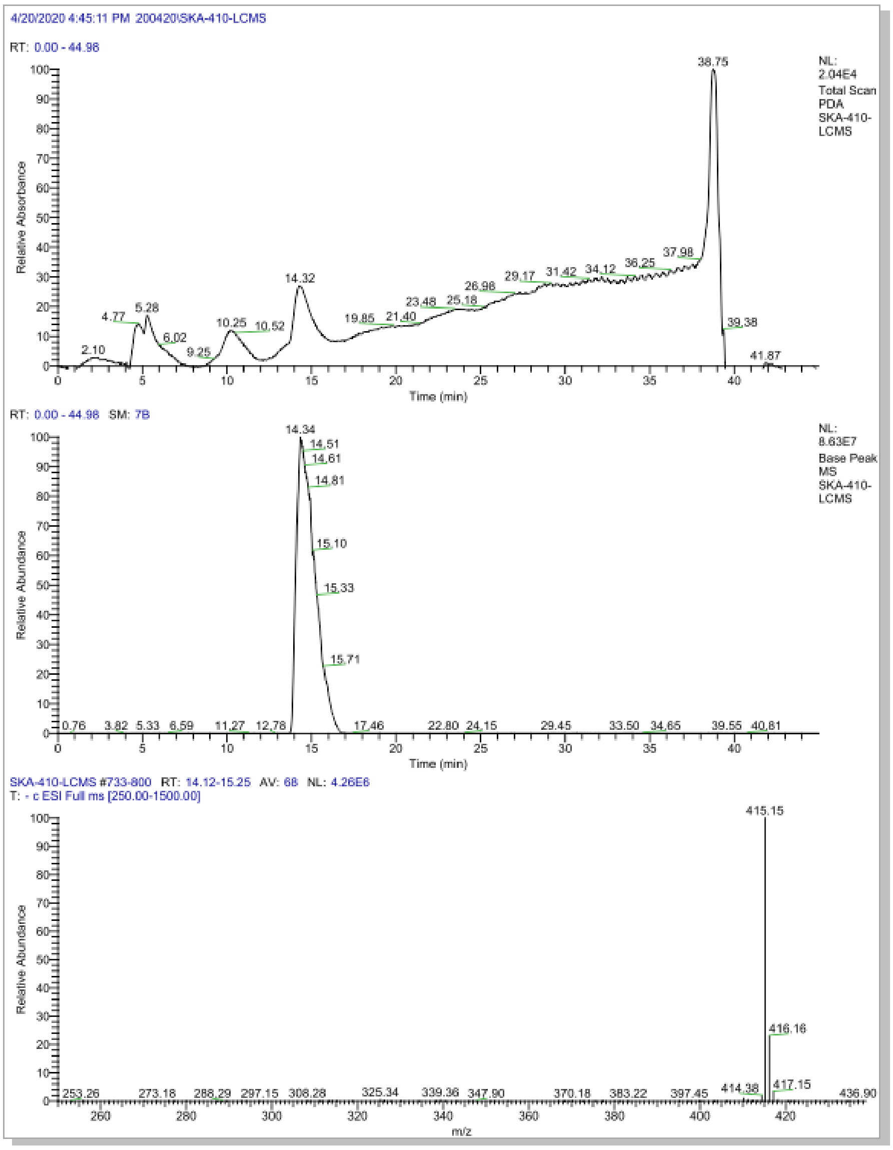

**Figure S36**. HR-MS and LC-MS spectra of compound **28**

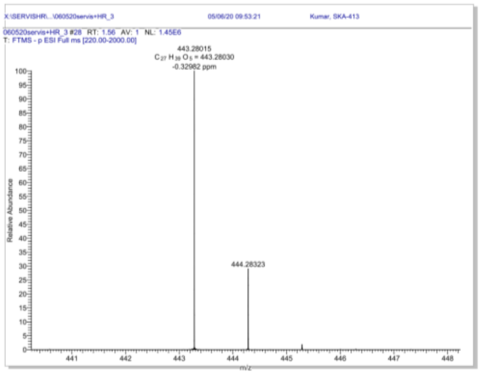


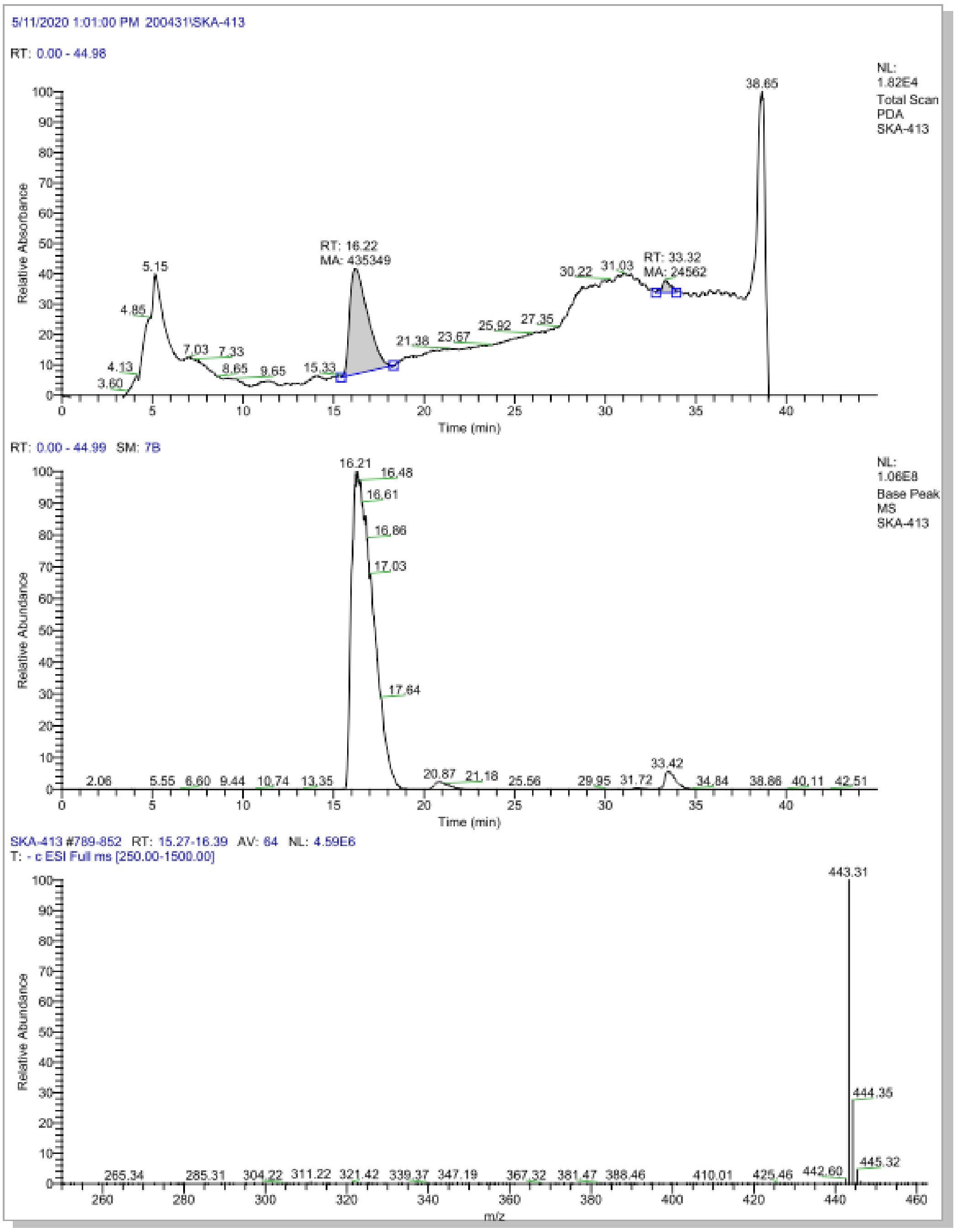
